## Supplementary material for "Uncertainty and precaution in hunting wolves twice in a year": Supp Info S1

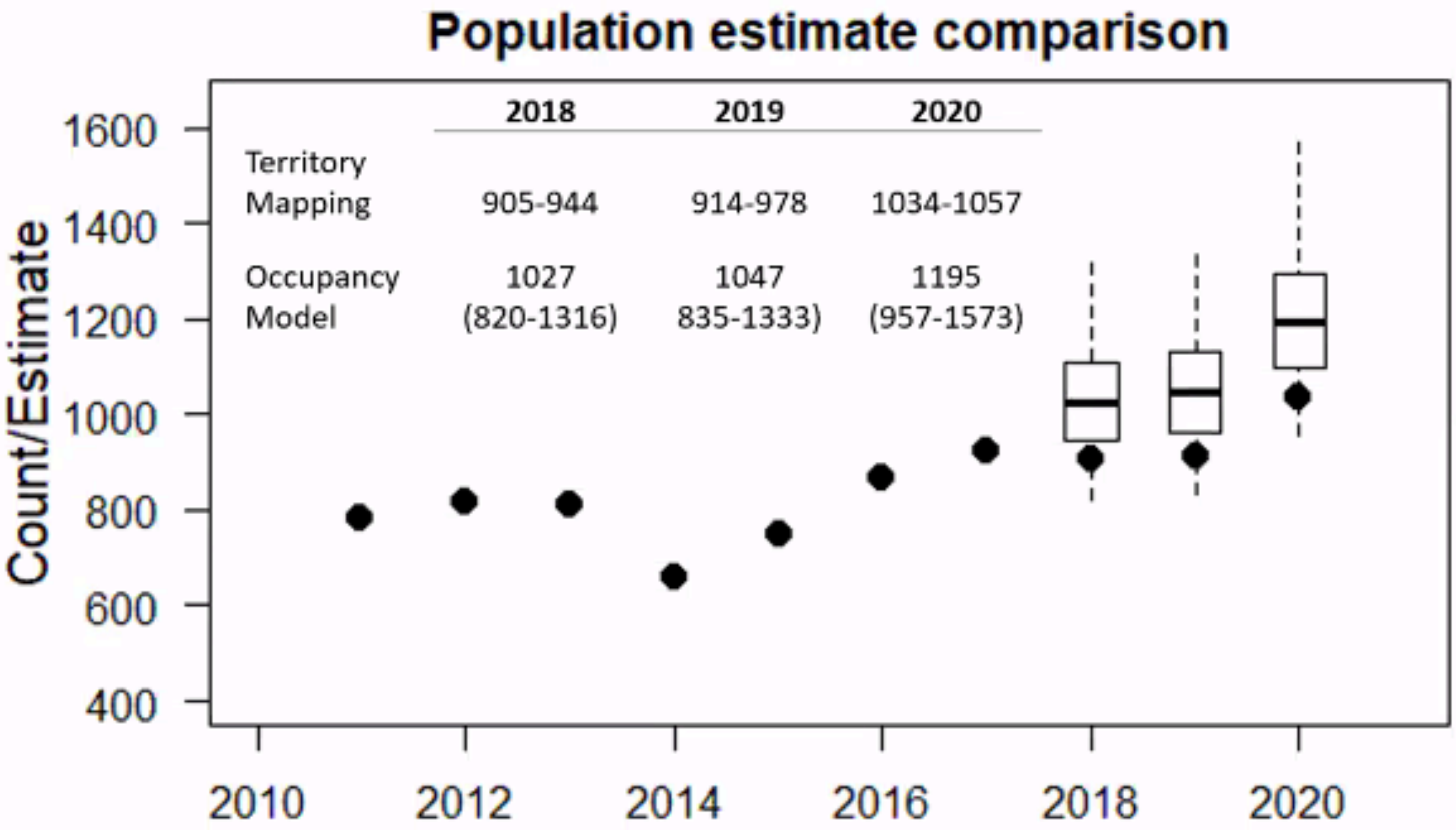
Population estimate comparison
2018
2019
2020
Territory Mapping
905-944
914-978
1034-1057
Occupancy Model
1027 (820-1316)
1047 835-1333)
1195 (957-1573)
Count/Estimate
1600
1400
1200
1000
800
600
400
2010
2012
2014
2016
2018
2020

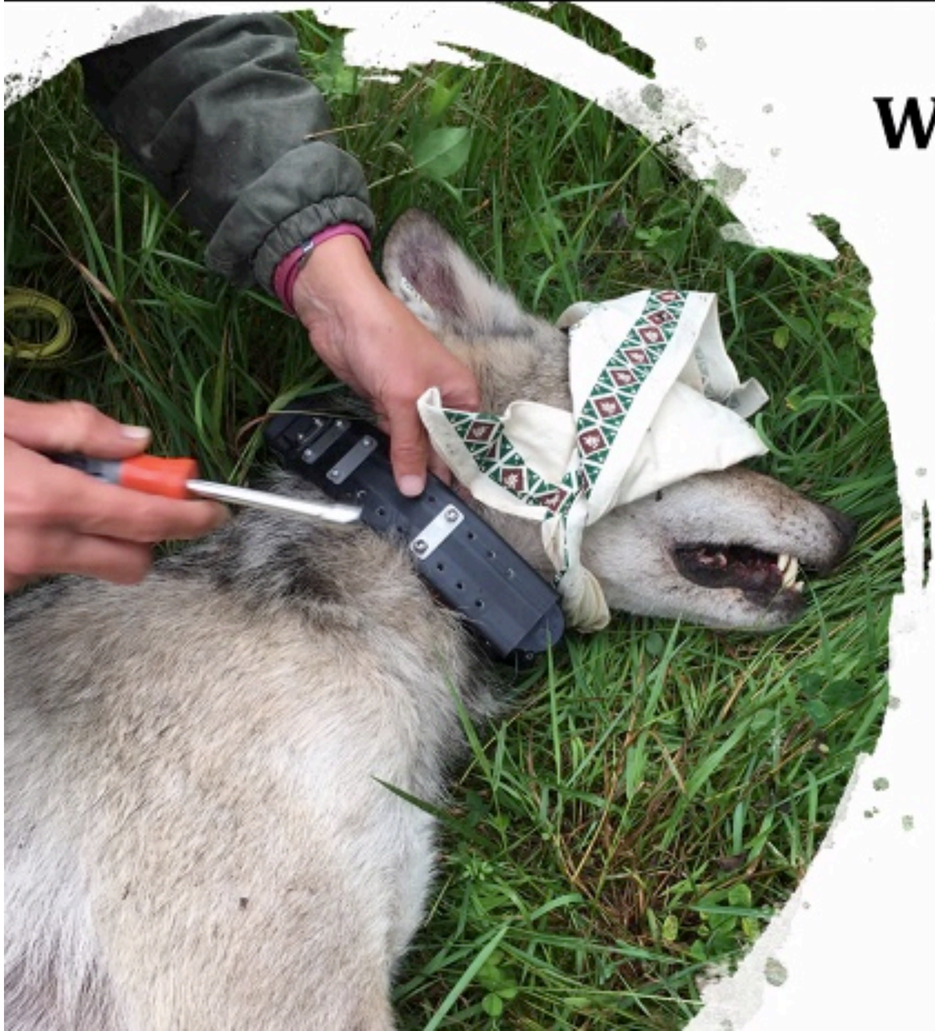

### Wolf Processing

Information Collected

- Measurements
  - Weight and Length (nose to tip of tail)
  - Neck and head diameter
  - For collar size and cable restraint size purposes
  - Toe pads & paws, shoulder to pad, eye to nose

**Currently collared wolves**

- **43** currently collared
  - 16 functioning correctly
  - 4 transmitting intermittently
  - 17 missing – collars stopped transmitting for unknown reasons
  - 6 are VHF collars
- 7 harvested during season
