## Supplementary material for "Uncertainty and precaution in hunting wolves twice in a year": Supp Info S2

Deaths uniform distribution between  0.38-0.56

| | | | | | | |
|---|---|---|---|---|---|---|
| **0.470** | 0.470 | 0.520 | 0.420 | 0.500 | 0.560 | 0.520 |
| **0.520** | 0.480 | 0.490 | 0.460 | 0.540 | 0.450 | 0.460 |
| **0.530** | 0.460 | 0.510 | 0.540 | 0.410 | 0.450 | 0.420 |
| **0.420** | 0.480 | 0.500 | 0.510 | 0.530 | 0.470 | 0.490 |
| **0.420** | 0.480 | 0.520 | 0.470 | 0.560 | 0.510 | 0.520 |
| **0.430** | 0.540 | 0.470 | 0.520 | 0.470 | 0.510 | 0.530 |
| **0.460** | 0.500 | 0.450 | 0.480 | 0.510 | 0.390 | 0.490 |
| **0.430** | 0.490 | 0.510 | 0.430 | 0.540 | 0.380 | 0.400 |
| **0.380** | 0.490 | 0.460 | 0.390 | 0.450 | 0.510 | 0.380 |
| **0.480** | 0.520 | 0.470 | 0.460 | 0.520 | 0.380 | 0.390 |
| **0.500** | 0.500 | 0.500 | 0.500 | 0.520 | 0.450 | 0.550 |
| **0.430** | 0.560 | 0.470 | 0.500 | 0.390 | 0.400 | 0.470 |
| **0.530** | 0.520 | 0.400 | 0.440 | 0.380 | 0.380 | 0.450 |
| **0.560** | 0.500 | 0.450 | 0.430 | 0.380 | 0.490 | 0.540 |
| **0.480** | 0.390 | 0.500 | 0.430 | 0.390 | 0.480 | 0.530 |
| **0.390** | 0.470 | 0.410 | 0.380 | 0.560 | 0.390 | 0.380 |
| **0.480** | 0.510 | 0.490 | 0.400 | 0.560 | 0.540 | 0.420 |
| **0.390** | 0.490 | 0.460 | 0.560 | 0.470 | 0.550 | 0.510 |
| **0.540** | 0.550 | 0.440 | 0.510 | 0.540 | 0.540 | 0.550 |
| **0.560** | 0.400 | 0.390 | 0.420 | 0.420 | 0.470 | 0.440 |
| **0.490** | 0.470 | 0.520 | 0.420 | 0.380 | 0.410 | 0.490 |
| **0.460** | 0.490 | 0.500 | 0.540 | 0.390 | 0.550 | 0.390 |
| **0.550** | 0.560 | 0.400 | 0.420 | 0.520 | 0.400 | 0.440 |
| **0.510** | 0.430 | 0.530 | 0.520 | 0.420 | 0.560 | 0.400 |
| **0.530** | 0.430 | 0.440 | 0.490 | 0.470 | 0.440 | 0.510 |
| **0.490** | 0.550 | 0.470 | 0.440 | 0.530 | 0.390 | 0.430 |
| **0.480** | 0.470 | 0.470 | 0.430 | 0.470 | 0.560 | 0.400 |
| **0.470** | 0.380 | 0.430 | 0.450 | 0.560 | 0.470 | 0.520 |
| **0.510** | 0.380 | 0.490 | 0.510 | 0.560 | 0.560 | 0.530 |
| **0.560** | 0.400 | 0.500 | 0.550 | 0.480 | 0.560 | 0.480 |
| **0.410** | 0.460 | 0.520 | 0.540 | 0.380 | 0.490 | 0.470 |
| **0.500** | 0.430 | 0.410 | 0.480 | 0.400 | 0.420 | 0.460 |
| **0.400** | 0.520 | 0.530 | 0.480 | 0.520 | 0.410 | 0.540 |
| **0.420** | 0.490 | 0.400 | 0.400 | 0.550 | 0.490 | 0.390 |
| **0.410** | 0.410 | 0.500 | 0.500 | 0.470 | 0.550 | 0.510 |
| **0.560** | 0.450 | 0.470 | 0.400 | 0.550 | 0.510 | 0.520 |
| **0.400** | 0.480 | 0.440 | 0.520 | 0.450 | 0.390 | 0.490 |
| **0.410** | 0.380 | 0.380 | 0.500 | 0.420 | 0.430 | 0.480 |
| **0.440** | 0.510 | 0.450 | 0.450 | 0.390 | 0.550 | 0.530 |
| **0.550** | 0.390 | 0.480 | 0.460 | 0.560 | 0.520 | 0.420 |
| **0.410** | 0.480 | 0.460 | 0.550 | 0.390 | 0.560 | 0.420 |
| **0.400** | 0.390 | 0.460 | 0.380 | 0.400 | 0.510 | 0.390 |
| **0.500** | 0.470 | 0.510 | 0.410 | 0.540 | 0.540 | 0.550 |
| **0.540** | 0.420 | 0.540 | 0.450 | 0.460 | 0.500 | 0.440 |
| **0.480** | 0.460 | 0.510 | 0.460 | 0.540 | 0.540 | 0.500 |
| **0.470** | 0.440 | 0.410 | 0.450 | 0.470 | 0.550 | 0.450 |

| | | | | | | |
|---|---|---|---|---|---|---|
| **0.410** | 0.540 | 0.520 | 0.500 | 0.550 | 0.530 | 0.520 |
| **0.390** | 0.400 | 0.530 | 0.460 | 0.500 | 0.540 | 0.560 |
| **0.450** | 0.420 | 0.480 | 0.480 | 0.520 | 0.560 | 0.540 |
| **0.530** | 0.510 | 0.500 | 0.420 | 0.510 | 0.530 | 0.460 |
| **0.450** | 0.470 | 0.510 | 0.460 | 0.550 | 0.410 | 0.560 |
| **0.380** | 0.390 | 0.510 | 0.540 | 0.430 | 0.500 | 0.460 |
| **0.550** | 0.460 | 0.410 | 0.440 | 0.390 | 0.480 | 0.450 |
| **0.450** | 0.430 | 0.380 | 0.530 | 0.480 | 0.390 | 0.470 |
| **0.390** | 0.470 | 0.420 | 0.430 | 0.540 | 0.410 | 0.470 |
| **0.420** | 0.500 | 0.410 | 0.450 | 0.560 | 0.450 | 0.490 |
| **0.540** | 0.410 | 0.440 | 0.470 | 0.440 | 0.410 | 0.540 |
| **0.410** | 0.450 | 0.550 | 0.390 | 0.460 | 0.540 | 0.540 |
| **0.400** | 0.440 | 0.490 | 0.410 | 0.470 | 0.380 | 0.540 |
| **0.420** | 0.500 | 0.500 | 0.540 | 0.420 | 0.460 | 0.420 |
| **0.480** | 0.380 | 0.400 | 0.440 | 0.380 | 0.440 | 0.430 |
| **0.540** | 0.430 | 0.470 | 0.540 | 0.530 | 0.390 | 0.430 |
| **0.440** | 0.390 | 0.390 | 0.500 | 0.480 | 0.560 | 0.540 |
| **0.480** | 0.510 | 0.550 | 0.530 | 0.480 | 0.540 | 0.500 |
| **0.470** | 0.490 | 0.510 | 0.500 | 0.400 | 0.530 | 0.540 |
| **0.560** | 0.420 | 0.410 | 0.560 | 0.430 | 0.520 | 0.460 |
| **0.500** | 0.390 | 0.470 | 0.400 | 0.420 | 0.410 | 0.550 |
| **0.520** | 0.470 | 0.440 | 0.440 | 0.480 | 0.540 | 0.380 |
| **0.540** | 0.500 | 0.490 | 0.520 | 0.550 | 0.410 | 0.460 |
| **0.440** | 0.470 | 0.490 | 0.530 | 0.380 | 0.500 | 0.420 |
| **0.550** | 0.470 | 0.380 | 0.560 | 0.430 | 0.380 | 0.560 |
| **0.440** | 0.470 | 0.540 | 0.420 | 0.540 | 0.560 | 0.540 |
| **0.440** | 0.520 | 0.400 | 0.500 | 0.450 | 0.480 | 0.420 |
| **0.470** | 0.490 | 0.420 | 0.380 | 0.410 | 0.540 | 0.510 |
| **0.530** | 0.500 | 0.400 | 0.510 | 0.510 | 0.420 | 0.430 |
| **0.540** | 0.400 | 0.510 | 0.450 | 0.410 | 0.430 | 0.450 |
| **0.500** | 0.500 | 0.560 | 0.520 | 0.530 | 0.380 | 0.510 |
| **0.550** | 0.500 | 0.550 | 0.380 | 0.410 | 0.430 | 0.410 |
| **0.520** | 0.380 | 0.450 | 0.550 | 0.520 | 0.390 | 0.490 |
| **0.530** | 0.560 | 0.410 | 0.540 | 0.510 | 0.520 | 0.510 |
| **0.510** | 0.530 | 0.400 | 0.440 | 0.440 | 0.520 | 0.510 |
| **0.480** | 0.420 | 0.510 | 0.410 | 0.470 | 0.440 | 0.420 |
| **0.380** | 0.520 | 0.410 | 0.400 | 0.520 | 0.520 | 0.400 |
| **0.450** | 0.520 | 0.480 | 0.510 | 0.510 | 0.440 | 0.410 |
| **0.560** | 0.550 | 0.540 | 0.470 | 0.380 | 0.480 | 0.430 |
| **0.410** | 0.450 | 0.550 | 0.540 | 0.450 | 0.560 | 0.540 |
| **0.460** | 0.410 | 0.410 | 0.550 | 0.540 | 0.470 | 0.420 |
| **0.420** | 0.400 | 0.410 | 0.470 | 0.430 | 0.430 | 0.550 |
| **0.500** | 0.400 | 0.480 | 0.450 | 0.430 | 0.510 | 0.470 |
| **0.450** | 0.490 | 0.510 | 0.430 | 0.430 | 0.390 | 0.400 |
| **0.490** | 0.470 | 0.460 | 0.430 | 0.450 | 0.420 | 0.480 |
| **0.390** | 0.520 | 0.550 | 0.560 | 0.430 | 0.490 | 0.510 |
| **0.540** | 0.560 | 0.500 | 0.380 | 0.380 | 0.540 | 0.380 |

| | | | | | | |
|---|---|---|---|---|---|---|
| **0.510** | 0.410 | 0.390 | 0.500 | 0.440 | 0.460 | 0.510 |
| **0.410** | 0.510 | 0.540 | 0.550 | 0.400 | 0.520 | 0.490 |
| **0.510** | 0.500 | 0.450 | 0.500 | 0.440 | 0.490 | 0.410 |
| **0.380** | 0.460 | 0.400 | 0.540 | 0.430 | 0.440 | 0.500 |
| **0.480** | 0.510 | 0.440 | 0.450 | 0.530 | 0.410 | 0.410 |
| **0.440** | 0.530 | 0.520 | 0.550 | 0.450 | 0.400 | 0.530 |
| **0.530** | 0.510 | 0.490 | 0.480 | 0.400 | 0.390 | 0.510 |
| **0.480** | 0.510 | 0.380 | 0.480 | 0.500 | 0.510 | 0.430 |
| **0.390** | 0.390 | 0.430 | 0.500 | 0.530 | 0.500 | 0.540 |
| **0.400** | 0.480 | 0.410 | 0.400 | 0.380 | 0.560 | 0.450 |
| **0.380** | 0.420 | 0.480 | 0.490 | 0.540 | 0.390 | 0.480 |
| **0.520** | 0.550 | 0.400 | 0.440 | 0.470 | 0.380 | 0.410 |
| **0.410** | 0.510 | 0.420 | 0.390 | 0.450 | 0.540 | 0.510 |
| **0.400** | 0.470 | 0.480 | 0.500 | 0.380 | 0.390 | 0.470 |
| **0.530** | 0.520 | 0.430 | 0.420 | 0.490 | 0.550 | 0.420 |
| **0.510** | 0.530 | 0.470 | 0.390 | 0.520 | 0.450 | 0.430 |
| **0.560** | 0.380 | 0.470 | 0.450 | 0.380 | 0.430 | 0.530 |
| **0.490** | 0.390 | 0.540 | 0.520 | 0.450 | 0.490 | 0.470 |
| **0.440** | 0.480 | 0.420 | 0.410 | 0.400 | 0.380 | 0.480 |
| **0.400** | 0.460 | 0.560 | 0.510 | 0.400 | 0.520 | 0.530 |
| **0.550** | 0.490 | 0.500 | 0.440 | 0.470 | 0.560 | 0.390 |
| **0.460** | 0.420 | 0.480 | 0.460 | 0.510 | 0.450 | 0.390 |
| **0.400** | 0.470 | 0.540 | 0.430 | 0.480 | 0.400 | 0.460 |
| **0.380** | 0.440 | 0.520 | 0.410 | 0.500 | 0.460 | 0.390 |
| **0.420** | 0.490 | 0.400 | 0.450 | 0.480 | 0.500 | 0.430 |
| **0.470** | 0.390 | 0.440 | 0.550 | 0.420 | 0.410 | 0.410 |
| **0.450** | 0.460 | 0.480 | 0.450 | 0.520 | 0.500 | 0.380 |

| | | | |
|---|---|---|---|
| **0.470** | 0.500 | 0.400 | 0.540 |
| **0.520** | 0.380 | 0.490 | 0.550 |
| **0.530** | 0.460 | 0.440 | 0.430 |
| **0.420** | 0.410 | 0.420 | 0.480 |
| **0.420** | 0.490 | 0.560 | 0.500 |
| **0.430** | 0.530 | 0.480 | 0.490 |
| **0.460** | 0.480 | 0.400 | 0.480 |
| **0.430** | 0.390 | 0.380 | 0.500 |
| **0.380** | 0.450 | 0.440 | 0.390 |
| **0.480** | 0.550 | 0.480 | 0.400 |
| **0.500** | 0.420 | 0.560 | 0.550 |
| **0.430** | 0.550 | 0.440 | 0.410 |
| **0.530** | 0.430 | 0.420 | 0.480 |
| **0.560** | 0.480 | 0.450 | 0.400 |
| **0.480** | 0.420 | 0.420 | 0.400 |
| **0.390** | 0.430 | 0.550 | 0.540 |
| **0.480** | 0.400 | 0.520 | 0.540 |
| **0.390** | 0.470 | 0.470 | 0.460 |
| **0.540** | 0.450 | 0.530 | 0.470 |
| **0.560** | 0.510 | 0.460 | 0.540 |
| **0.490** | 0.410 | 0.550 | 0.410 |
| **0.460** | 0.480 | 0.530 | 0.520 |
| **0.550** | 0.450 | 0.400 | 0.450 |
| **0.510** | 0.480 | 0.430 | 0.510 |
| **0.530** | 0.550 | 0.420 | 0.510 |
| **0.490** | 0.450 | 0.450 | 0.540 |
| **0.480** | 0.430 | 0.440 | 0.500 |
| **0.470** | 0.440 | 0.460 | 0.530 |
| **0.510** | 0.480 | 0.450 | 0.530 |
| **0.560** | 0.410 | 0.460 | 0.390 |
| **0.410** | 0.490 | 0.440 | 0.540 |
| **0.500** | 0.430 | 0.550 | 0.470 |
| **0.400** | 0.460 | 0.410 | 0.500 |
| **0.420** | 0.480 | 0.500 | 0.420 |
| **0.410** | 0.430 | 0.550 | 0.390 |
| **0.560** | 0.520 | 0.430 | 0.530 |
| **0.400** | 0.380 | 0.450 | 0.490 |
| **0.410** | 0.540 | 0.420 | 0.440 |
| **0.440** | 0.560 | 0.430 | 0.500 |
| **0.550** | 0.480 | 0.420 | 0.530 |
| **0.410** | 0.510 | 0.440 | 0.550 |
| **0.400** | 0.530 | 0.410 | 0.430 |
| **0.500** | 0.520 | 0.490 | 0.500 |
| **0.540** | 0.520 | 0.560 | 0.520 |
| **0.480** | 0.430 | 0.490 | 0.440 |
| **0.470** | 0.430 | 0.540 | 0.470 |

| | | | |
|---|---|---|---|
| **0.410** | 0.450 | 0.550 | 0.540 |
| **0.390** | 0.380 | 0.400 | 0.440 |
| **0.450** | 0.510 | 0.380 | 0.390 |
| **0.530** | 0.470 | 0.450 | 0.490 |
| **0.450** | 0.380 | 0.430 | 0.400 |
| **0.380** | 0.510 | 0.460 | 0.560 |
| **0.550** | 0.390 | 0.460 | 0.420 |
| **0.450** | 0.420 | 0.460 | 0.490 |
| **0.390** | 0.420 | 0.470 | 0.390 |
| **0.420** | 0.500 | 0.430 | 0.390 |
| **0.540** | 0.500 | 0.500 | 0.480 |
| **0.410** | 0.470 | 0.380 | 0.490 |
| **0.400** | 0.520 | 0.460 | 0.510 |
| **0.420** | 0.390 | 0.460 | 0.470 |
| **0.480** | 0.560 | 0.540 | 0.420 |
| **0.540** | 0.460 | 0.470 | 0.550 |
| **0.440** | 0.490 | 0.500 | 0.550 |
| **0.480** | 0.440 | 0.520 | 0.520 |
| **0.470** | 0.470 | 0.520 | 0.460 |
| **0.560** | 0.380 | 0.510 | 0.420 |
| **0.500** | 0.400 | 0.430 | 0.510 |
| **0.520** | 0.440 | 0.420 | 0.550 |
| **0.540** | 0.520 | 0.530 | 0.390 |
| **0.440** | 0.380 | 0.560 | 0.500 |
| **0.550** | 0.550 | 0.390 | 0.420 |
| **0.440** | 0.500 | 0.550 | 0.550 |
| **0.440** | 0.510 | 0.450 | 0.440 |
| **0.470** | 0.380 | 0.400 | 0.470 |
| **0.530** | 0.520 | 0.530 | 0.450 |
| **0.540** | 0.490 | 0.530 | 0.440 |
| **0.500** | 0.420 | 0.560 | 0.450 |
| **0.550** | 0.430 | 0.510 | 0.470 |
| **0.520** | 0.540 | 0.390 | 0.460 |
| **0.530** | 0.430 | 0.510 | 0.460 |
| **0.510** | 0.500 | 0.380 | 0.390 |
| **0.480** | 0.420 | 0.440 | 0.520 |
| **0.380** | 0.380 | 0.550 | 0.480 |
| **0.450** | 0.380 | 0.400 | 0.440 |
| **0.560** | 0.550 | 0.420 | 0.390 |
| **0.410** | 0.430 | 0.560 | 0.440 |
| **0.460** | 0.400 | 0.530 | 0.550 |
| **0.420** | 0.410 | 0.500 | 0.430 |
| **0.500** | 0.450 | 0.430 | 0.460 |
| **0.450** | 0.500 | 0.380 | 0.480 |
| **0.490** | 0.460 | 0.400 | 0.390 |
| **0.390** | 0.450 | 0.440 | 0.490 |
| **0.540** | 0.520 | 0.530 | 0.380 |

| | | | |
|---|---|---|---|
| **0.510** | 0.540 | 0.510 | 0.480 |
| **0.410** | 0.410 | 0.480 | 0.470 |
| **0.510** | 0.530 | 0.400 | 0.540 |
| **0.380** | 0.510 | 0.470 | 0.440 |
| **0.480** | 0.480 | 0.460 | 0.500 |
| **0.440** | 0.520 | 0.430 | 0.510 |
| **0.530** | 0.440 | 0.400 | 0.440 |
| **0.480** | 0.390 | 0.540 | 0.510 |
| **0.390** | 0.430 | 0.440 | 0.430 |
| **0.400** | 0.400 | 0.510 | 0.410 |
| **0.380** | 0.550 | 0.470 | 0.410 |
| **0.520** | 0.400 | 0.490 | 0.490 |
| **0.410** | 0.440 | 0.410 | 0.550 |
| **0.400** | 0.520 | 0.530 | 0.440 |
| **0.530** | 0.480 | 0.450 | 0.530 |
| **0.510** | 0.550 | 0.500 | 0.480 |
| **0.560** | 0.520 | 0.430 | 0.560 |
| **0.490** | 0.540 | 0.410 | 0.530 |
| **0.440** | 0.520 | 0.480 | 0.480 |
| **0.400** | 0.440 | 0.500 | 0.490 |
| **0.550** | 0.450 | 0.520 | 0.390 |
| **0.460** | 0.440 | 0.480 | 0.460 |
| **0.400** | 0.470 | 0.420 | 0.520 |
| **0.380** | 0.480 | 0.450 | 0.420 |
| **0.420** | 0.510 | 0.380 | 0.430 |
| **0.470** | 0.410 | 0.500 | 0.500 |
| **0.450** | 0.510 | 0.430 | 0.430 |

Packs with pups normal centered 0.72 (range 0.55-0.89)

| **0.72** | **0.68** | **0.68** | **0.56** | **0.65** | **0.60** | **0.75** | **0.62** | **0.59** | **0.75** |
|---|---|---|---|---|---|---|---|---|---|
| **0.72** | 0.53 | 0.69 | 0.66 | 0.58 | 0.66 | 0.72 | 0.79 | 0.60 | 0.67 |
| **0.65** | 0.67 | 0.66 | 0.58 | 0.65 | 0.62 | 0.60 | 0.63 | 0.70 | 0.63 |
| **0.62** | 0.72 | 0.82 | 0.63 | 0.69 | 0.68 | 0.79 | 0.65 | 0.60 | 0.72 |
| **0.65** | 0.69 | 0.81 | 0.68 | 0.75 | 0.67 | 0.65 | 0.73 | 0.69 | 0.65 |
| **0.74** | 0.67 | 0.77 | 0.68 | 0.81 | 0.65 | 0.69 | 0.62 | 0.80 | 0.75 |
| **0.67** | 0.80 | 0.79 | 0.75 | 0.65 | 0.60 | 0.71 | 0.54 | 0.72 | 0.64 |
| **0.80** | 0.67 | 0.76 | 0.62 | 0.74 | 0.68 | 0.66 | 0.72 | 0.72 | 0.83 |
| **0.58** | 0.64 | 0.71 | 0.70 | 0.76 | 0.77 | 0.72 | 0.77 | 0.53 | 0.83 |
| **0.60** | 0.71 | 0.74 | 0.67 | 0.75 | 0.71 | 0.58 | 0.76 | 0.78 | 0.72 |
| **0.72** | 0.77 | 0.61 | 0.68 | 0.66 | 0.62 | 0.65 | 0.74 | 0.57 | 0.69 |
| **0.72** | 0.62 | 0.66 | 0.70 | 0.53 | 0.60 | 0.59 | 0.60 | 0.65 | 0.68 |
| **0.55** | 0.81 | 0.71 | 0.79 | 0.70 | 0.66 | 0.73 | 0.69 | 0.73 | 0.80 |
| **0.79** | 0.61 | 0.75 | 0.63 | 0.61 | 0.71 | 0.72 | 0.75 | 0.70 | 0.67 |
| **0.66** | 0.74 | 0.62 | 0.66 | 0.65 | 0.74 | 0.60 | 0.75 | 0.53 | 0.74 |
| **0.67** | 0.73 | 0.71 | 0.64 | 0.68 | 0.71 | 0.59 | 0.75 | 0.57 | 0.57 |
| **0.57** | 0.58 | 0.83 | 0.63 | 0.70 | 0.68 | 0.62 | 0.67 | 0.64 | 0.70 |
| **0.65** | 0.69 | 0.67 | 0.74 | 0.59 | 0.70 | 0.62 | 0.73 | 0.77 | 0.69 |
| **0.75** | 0.81 | 0.69 | 0.60 | 0.59 | 0.72 | 0.69 | 0.70 | 0.64 | 0.76 |
| **0.67** | 0.56 | 0.79 | 0.76 | 0.64 | 0.72 | 0.69 | 0.66 | 0.75 | 0.81 |
| **0.66** | 0.60 | 0.80 | 0.56 | 0.75 | 0.68 | 0.69 | 0.74 | 0.60 | 0.67 |
| **0.67** | 0.64 | 0.68 | 0.66 | 0.73 | 0.81 | 0.77 | 0.55 | 0.60 | 0.71 |
| **0.69** | 0.66 | 0.68 | 0.59 | 0.69 | 0.66 | 0.73 | 0.71 | 0.64 | 0.72 |
| **0.74** | 0.79 | 0.67 | 0.68 | 0.65 | 0.72 | 0.79 | 0.60 | 0.64 | 0.68 |
| **0.59** | 0.65 | 0.83 | 0.81 | 0.62 | 0.79 | 0.66 | 0.67 | 0.68 | 0.72 |
| **0.72** | 0.71 | 0.58 | 0.60 | 0.62 | 0.66 | 0.69 | 0.77 | 0.71 | 0.64 |
| **0.73** | 0.68 | 0.53 | 0.78 | 0.62 | 0.55 | 0.64 | 0.68 | 0.70 | 0.64 |
| **0.68** | 0.72 | 0.62 | 0.74 | 0.69 | 0.71 | 0.65 | 0.64 | 0.59 | 0.67 |
| **0.69** | 0.67 | 0.59 | 0.64 | 0.66 | 0.79 | 0.74 | 0.59 | 0.66 | 0.68 |
| **0.70** | 0.66 | 0.74 | 0.69 | 0.68 | 0.71 | 0.76 | 0.55 | 0.67 | 0.77 |
| **0.66** | 0.61 | 0.60 | 0.54 | 0.70 | 0.68 | 0.60 | 0.63 | 0.83 | 0.67 |
| **0.71** | 0.63 | 0.76 | 0.71 | 0.70 | 0.61 | 0.66 | 0.74 | 0.65 | 0.67 |
| **0.68** | 0.66 | 0.68 | 0.82 | 0.76 | 0.67 | 0.76 | 0.64 | 0.72 | 0.80 |
| **0.70** | 0.67 | 0.59 | 0.62 | 0.76 | 0.66 | 0.64 | 0.68 | 0.67 | 0.64 |
| **0.63** | 0.69 | 0.62 | 0.73 | 0.77 | 0.74 | 0.62 | 0.68 | 0.70 | 0.65 |
| **0.82** | 0.60 | 0.53 | 0.81 | 0.74 | 0.63 | 0.76 | 0.77 | 0.69 | 0.68 |
| **0.72** | 0.61 | 0.70 | 0.77 | 0.61 | 0.81 | 0.54 | 0.74 | 0.73 | 0.74 |
| **0.67** | 0.71 | 0.70 | 0.69 | 0.72 | 0.62 | 0.66 | 0.66 | 0.78 | 0.61 |
| **0.73** | 0.71 | 0.79 | 0.58 | 0.60 | 0.63 | 0.68 | 0.77 | 0.70 | 0.67 |
| **0.76** | 0.68 | 0.55 | 0.67 | 0.57 | 0.67 | 0.71 | 0.66 | 0.59 | 0.65 |
| **0.59** | 0.72 | 0.66 | 0.74 | 0.70 | 0.63 | 0.63 | 0.67 | 0.80 | 0.68 |
| **0.62** | 0.70 | 0.59 | 0.63 | 0.74 | 0.74 | 0.65 | 0.75 | 0.65 | 0.65 |
| **0.58** | 0.66 | 0.75 | 0.66 | 0.63 | 0.80 | 0.76 | 0.72 | 0.72 | 0.59 |
| **0.59** | 0.80 | 0.75 | 0.70 | 0.60 | 0.65 | 0.68 | 0.73 | 0.54 | 0.74 |
| **0.71** | 0.63 | 0.83 | 0.63 | 0.59 | 0.64 | 0.74 | 0.58 | 0.77 | 0.72 |
| **0.78** | 0.81 | 0.67 | 0.57 | 0.70 | 0.76 | 0.69 | 0.56 | 0.72 | 0.71 |
| **0.70** | 0.66 | 0.79 | 0.79 | 0.71 | 0.62 | 0.70 | 0.58 | 0.76 | 0.78 |
| **0.65** | 0.71 | 0.63 | 0.81 | 0.67 | 0.64 | 0.74 | 0.61 | 0.61 | 0.70 |
| **0.77** | 0.61 | 0.75 | 0.78 | 0.77 | 0.78 | 0.78 | 0.61 | 0.62 | 0.62 |
| **0.69** | 0.67 | 0.61 | 0.66 | 0.78 | 0.79 | 0.67 | 0.67 | 0.72 | 0.59 |
| **0.66** | 0.68 | 0.62 | 0.70 | 0.60 | 0.63 | 0.55 | 0.72 | 0.58 | 0.55 |
| **0.58** | 0.65 | 0.57 | 0.60 | 0.54 | 0.67 | 0.73 | 0.70 | 0.59 | 0.62 |
| **0.66** | 0.64 | 0.82 | 0.69 | 0.66 | 0.65 | 0.78 | 0.59 | 0.75 | 0.56 |
| **0.57** | 0.73 | 0.78 | 0.73 | 0.68 | 0.66 | 0.64 | 0.55 | 0.68 | 0.77 |
| **0.60** | 0.71 | 0.60 | 0.59 | 0.64 | 0.70 | 0.73 | 0.69 | 0.68 | 0.67 |
| **0.57** | 0.62 | 0.62 | 0.61 | 0.72 | 0.72 | 0.70 | 0.61 | 0.73 | 0.58 |
| **0.59** | 0.61 | 0.61 | 0.77 | 0.71 | 0.70 | 0.70 | 0.66 | 0.79 | 0.68 |
| **0.66** | 0.54 | 0.63 | 0.69 | 0.79 | 0.72 | 0.79 | 0.76 | 0.60 | 0.68 |
| **0.68** | 0.69 | 0.77 | 0.68 | 0.71 | 0.72 | 0.77 | 0.69 | 0.76 | 0.54 |
| **0.73** | 0.71 | 0.69 | 0.60 | 0.64 | 0.76 | 0.68 | 0.74 | 0.68 | 0.75 |
| **0.60** | 0.76 | 0.76 | 0.75 | 0.66 | 0.72 | 0.65 | 0.69 | 0.74 | 0.63 |
| **0.65** | 0.73 | 0.68 | 0.66 | 0.65 | 0.78 | 0.69 | 0.70 | 0.73 | 0.67 |
| **0.74** | 0.60 | 0.79 | 0.80 | 0.59 | 0.78 | 0.72 | 0.74 | 0.60 | 0.61 |
| **0.68** | 0.61 | 0.77 | 0.79 | 0.63 | 0.71 | 0.75 | 0.77 | 0.71 | 0.74 |
| **0.73** | 0.66 | 0.60 | 0.64 | 0.64 | 0.68 | 0.67 | 0.60 | 0.58 | 0.64 |
| **0.59** | 0.59 | 0.76 | 0.66 | 0.60 | 0.68 | 0.68 | 0.75 | 0.77 | 0.69 |
| **0.60** | 0.56 | 0.68 | 0.54 | 0.75 | 0.76 | 0.69 | 0.82 | 0.70 | 0.77 |

| **0.72** | **0.68** | **0.68** | **0.56** | **0.65** | **0.60** | **0.75** | **0.62** | **0.59** | **0.75** |
|---|---|---|---|---|---|---|---|---|---|
| **0.71** | 0.60 | 0.62 | 0.74 | 0.73 | 0.77 | 0.65 | 0.70 | 0.66 | 0.64 |
| **0.66** | 0.64 | 0.66 | 0.64 | 0.68 | 0.60 | 0.69 | 0.59 | 0.82 | 0.71 |
| **0.59** | 0.68 | 0.73 | 0.69 | 0.69 | 0.69 | 0.57 | 0.68 | 0.66 | 0.55 |
| **0.65** | 0.63 | 0.67 | 0.62 | 0.69 | 0.65 | 0.75 | 0.80 | 0.79 | 0.78 |
| **0.74** | 0.58 | 0.73 | 0.63 | 0.69 | 0.66 | 0.63 | 0.71 | 0.57 | 0.74 |
| **0.67** | 0.70 | 0.66 | 0.75 | 0.71 | 0.67 | 0.63 | 0.65 | 0.67 | 0.77 |
| **0.70** | 0.72 | 0.67 | 0.65 | 0.78 | 0.70 | 0.74 | 0.62 | 0.64 | 0.59 |
| **0.78** | 0.70 | 0.74 | 0.60 | 0.67 | 0.70 | 0.67 | 0.64 | 0.64 | 0.76 |
| **0.67** | 0.66 | 0.63 | 0.58 | 0.58 | 0.59 | 0.73 | 0.72 | 0.64 | 0.61 |
| **0.68** | 0.62 | 0.60 | 0.72 | 0.82 | 0.64 | 0.60 | 0.54 | 0.79 | 0.66 |
| **0.61** | 0.59 | 0.73 | 0.72 | 0.75 | 0.75 | 0.73 | 0.72 | 0.76 | 0.57 |
| **0.57** | 0.62 | 0.71 | 0.69 | 0.69 | 0.64 | 0.63 | 0.80 | 0.67 | 0.67 |
| **0.82** | 0.68 | 0.56 | 0.68 | 0.73 | 0.71 | 0.70 | 0.64 | 0.68 | 0.69 |
| **0.76** | 0.70 | 0.57 | 0.81 | 0.79 | 0.59 | 0.59 | 0.63 | 0.75 | 0.57 |
| **0.83** | 0.68 | 0.56 | 0.69 | 0.56 | 0.55 | 0.65 | 0.79 | 0.68 | 0.67 |
| **0.60** | 0.68 | 0.73 | 0.66 | 0.71 | 0.69 | 0.78 | 0.78 | 0.63 | 0.69 |
| **0.79** | 0.73 | 0.63 | 0.65 | 0.67 | 0.70 | 0.64 | 0.79 | 0.74 | 0.71 |
| **0.73** | 0.80 | 0.66 | 0.76 | 0.71 | 0.67 | 0.71 | 0.73 | 0.74 | 0.69 |
| **0.65** | 0.61 | 0.64 | 0.62 | 0.65 | 0.66 | 0.58 | 0.71 | 0.68 | 0.71 |
| **0.70** | 0.65 | 0.70 | 0.57 | 0.71 | 0.69 | 0.56 | 0.55 | 0.69 | 0.69 |
| **0.62** | 0.66 | 0.69 | 0.66 | 0.69 | 0.67 | 0.67 | 0.72 | 0.62 | 0.56 |
| **0.77** | 0.64 | 0.53 | 0.65 | 0.74 | 0.69 | 0.73 | 0.59 | 0.60 | 0.74 |
| **0.66** | 0.64 | 0.55 | 0.63 | 0.77 | 0.78 | 0.64 | 0.71 | 0.77 | 0.60 |
| **0.63** | 0.66 | 0.64 | 0.78 | 0.57 | 0.76 | 0.64 | 0.78 | 0.66 | 0.64 |
| **0.68** | 0.73 | 0.56 | 0.81 | 0.70 | 0.57 | 0.60 | 0.64 | 0.52 | 0.60 |
| **0.69** | 0.70 | 0.68 | 0.65 | 0.66 | 0.55 | 0.67 | 0.70 | 0.65 | 0.60 |
| **0.74** | 0.76 | 0.63 | 0.68 | 0.76 | 0.70 | 0.61 | 0.68 | 0.69 | 0.66 |
| **0.76** | 0.57 | 0.71 | 0.71 | 0.63 | 0.76 | 0.82 | 0.55 | 0.70 | 0.78 |
| **0.78** | 0.72 | 0.63 | 0.68 | 0.67 | 0.83 | 0.80 | 0.69 | 0.65 | 0.60 |
| **0.63** | 0.78 | 0.72 | 0.76 | 0.70 | 0.74 | 0.70 | 0.65 | 0.71 | 0.81 |
| **0.71** | 0.65 | 0.62 | 0.72 | 0.54 | 0.55 | 0.60 | 0.66 | 0.68 | 0.68 |
| **0.66** | 0.75 | 0.66 | 0.60 | 0.70 | 0.76 | 0.76 | 0.73 | 0.65 | 0.66 |
| **0.78** | 0.69 | 0.65 | 0.71 | 0.66 | 0.80 | 0.82 | 0.75 | 0.61 | 0.76 |
| **0.71** | 0.65 | 0.60 | 0.75 | 0.71 | 0.78 | 0.82 | 0.69 | 0.65 | 0.68 |
| **0.55** | 0.62 | 0.71 | 0.61 | 0.63 | 0.80 | 0.75 | 0.67 | 0.69 | 0.53 |
| **0.71** | 0.78 | 0.74 | 0.67 | 0.59 | 0.76 | 0.77 | 0.77 | 0.67 | 0.57 |
| **0.73** | 0.71 | 0.56 | 0.59 | 0.61 | 0.73 | 0.70 | 0.64 | 0.68 | 0.72 |
| **0.59** | 0.64 | 0.65 | 0.67 | 0.57 | 0.70 | 0.65 | 0.64 | 0.78 | 0.65 |
| **0.69** | 0.64 | 0.76 | 0.56 | 0.82 | 0.64 | 0.70 | 0.72 | 0.68 | 0.64 |
| **0.59** | 0.60 | 0.64 | 0.75 | 0.58 | 0.74 | 0.78 | 0.61 | 0.68 | 0.62 |
| **0.57** | 0.66 | 0.63 | 0.64 | 0.68 | 0.63 | 0.79 | 0.74 | 0.69 | 0.78 |
| **0.71** | 0.65 | 0.71 | 0.54 | 0.77 | 0.73 | 0.73 | 0.60 | 0.73 | 0.74 |
| **0.81** | 0.57 | 0.58 | 0.64 | 0.71 | 0.62 | 0.62 | 0.67 | 0.57 | 0.64 |
| **0.63** | 0.69 | 0.67 | 0.71 | 0.63 | 0.63 | 0.64 | 0.64 | 0.65 | 0.72 |
| **0.69** | 0.67 | 0.63 | 0.61 | 0.74 | 0.63 | 0.60 | 0.66 | 0.64 | 0.58 |
| **0.68** | 0.67 | 0.58 | 0.63 | 0.65 | 0.71 | 0.57 | 0.72 | 0.62 | 0.64 |
| **0.58** | 0.63 | 0.71 | 0.75 | 0.56 | 0.77 | 0.69 | 0.62 | 0.62 | 0.59 |
| **0.62** | 0.63 | 0.72 | 0.60 | 0.62 | 0.69 | 0.62 | 0.71 | 0.66 | 0.77 |
| **0.66** | 0.63 | 0.80 | 0.59 | 0.69 | 0.64 | 0.75 | 0.65 | 0.69 | 0.64 |
| **0.79** | 0.65 | 0.61 | 0.74 | 0.80 | 0.68 | 0.64 | 0.71 | 0.66 | 0.76 |
| **0.59** | 0.68 | 0.59 | 0.66 | 0.62 | 0.74 | 0.69 | 0.69 | 0.72 | 0.67 |
| **0.74** | 0.79 | 0.74 | 0.71 | 0.66 | 0.80 | 0.79 | 0.58 | 0.69 | 0.74 |
| **0.69** | 0.70 | 0.75 | 0.56 | 0.74 | 0.54 | 0.55 | 0.57 | 0.66 | 0.61 |

Litter size 4.8 (3-6) normal centered on 4.8

| 4 | 5 | 4 | 4 | 3 | 4 | 4 | 4 | 6 | 6 |
|---|---|---|---|---|---|---|---|---|---|
| 3 | 4 | 4 | 4 | 5 | 4 | 3 | 5 | 3 | 5 |
| 4 | 3 | 5 | 4 | 3 | 5 | 5 | 3 | 4 | 4 |
| 5 | 5 | 5 | 4 | 4 | 3 | 4 | 4 | 5 | 4 |
| 5 | 4 | 4 | 4 | 4 | 5 | 3 | 3 | 4 | 5 |
| 4 | 4 | 5 | 4 | 4 | 5 | 6 | 5 | 5 | 4 |
| 3 | 4 | 4 | 4 | 4 | 5 | 5 | 5 | 4 | 5 |
| 5 | 4 | 3 | 4 | 4 | 4 | 4 | 4 | 4 | 4 |
| 5 | 5 | 5 | 4 | 4 | 5 | 3 | 4 | 4 | 5 |
| 4 | 4 | 4 | 4 | 5 | 4 | 3 | 4 | 5 | 4 |
| 4 | 4 | 4 | 5 | 5 | 5 | 3 | 5 | 5 | 5 |
| 5 | 4 | 5 | 4 | 5 | 5 | 3 | 5 | 5 | 4 |
| 5 | 3 | 4 | 5 | 4 | 5 | 5 | 6 | 5 | 3 |
| 4 | 3 | 5 | 3 | 4 | 4 | 3 | 4 | 3 | 5 |
| 4 | 5 | 4 | 6 | 4 | 4 | 5 | 5 | 4 | 4 |
| 5 | 3 | 3 | 4 | 4 | 4 | 5 | 5 | 6 | 4 |
| 4 | 4 | 4 | 4 | 5 | 4 | 4 | 4 | 4 | 4 |
| 4 | 3 | 6 | 6 | 6 | 3 | 5 | 3 | 4 | 3 |
| 4 | 5 | 4 | 4 | 6 | 4 | 4 | 4 | 3 | 6 |
| 5 | 4 | 4 | 4 | 5 | 5 | 3 | 4 | 3 | 5 |
| 4 | 5 | 5 | 3 | 5 | 3 | 5 | 5 | 5 | 5 |
| 5 | 4 | 5 | 4 | 5 | 5 | 4 | 4 | 4 | 5 |
| 4 | 3 | 4 | 5 | 4 | 4 | 4 | 4 | 3 | 5 |
| 4 | 5 | 5 | 4 | 6 | 4 | 4 | 3 | 4 | 4 |
| 5 | 4 | 4 | 3 | 6 | 3 | 4 | 4 | 5 | 4 |
| 3 | 5 | 4 | 4 | 4 | 3 | 4 | 3 | 5 | 4 |
| 3 | 3 | 5 | 5 | 6 | 4 | 4 | 6 | 3 | 4 |
| 5 | 3 | 4 | 4 | 3 | 4 | 4 | 4 | 3 | 4 |
| 4 | 3 | 3 | 5 | 5 | 4 | 4 | 5 | 3 | 5 |
| 3 | 5 | 5 | 5 | 3 | 3 | 3 | 6 | 3 | 4 |
| 4 | 5 | 5 | 4 | 6 | 4 | 4 | 5 | 4 | 4 |
| 4 | 4 | 4 | 5 | 4 | 3 | 3 | 4 | 5 | 4 |
| 3 | 4 | 5 | 4 | 4 | 3 | 5 | 5 | 5 | 3 |
| 4 | 4 | 5 | 4 | 4 | 4 | 4 | 6 | 4 | 3 |
| 4 | 5 | 3 | 3 | 4 | 3 | 3 | 3 | 5 | 4 |
| 6 | 5 | 4 | 3 | 5 | 4 | 6 | 5 | 5 | 4 |
| 4 | 5 | 4 | 5 | 5 | 5 | 5 | 4 | 5 | 3 |
| 5 | 4 | 4 | 4 | 5 | 4 | 4 | 4 | 6 | 4 |
| 3 | 4 | 4 | 4 | 4 | 5 | 4 | 5 | 4 | 4 |
| 5 | 5 | 3 | 4 | 3 | 4 | 4 | 4 | 4 | 4 |
| 4 | 3 | 4 | 4 | 3 | 4 | 6 | 6 | 5 | 3 |
| 3 | 4 | 4 | 4 | 5 | 5 | 3 | 4 | 5 | 5 |
| 5 | 5 | 4 | 4 | 5 | 4 | 5 | 4 | 3 | 4 |
| 4 | 4 | 5 | 6 | 4 | 5 | 4 | 3 | 3 | 5 |
| 3 | 4 | 5 | 3 | 4 | 4 | 5 | 5 | 6 | 4 |
| 4 | 4 | 3 | 4 | 3 | 4 | 3 | 3 | 3 | 5 |
| 4 | 3 | 5 | 4 | 3 | 5 | 3 | 4 | 5 | 4 |
| 5 | 5 | 5 | 4 | 3 | 4 | 4 | 4 | 6 | 4 |
| 3 | 5 | 4 | 4 | 5 | 5 | 4 | 3 | 5 | 4 |
| 3 | 5 | 3 | 5 | 4 | 5 | 4 | 4 | 4 | 5 |
| 5 | 4 | 6 | 4 | 4 | 4 | 4 | 3 | 3 | 4 |
| 5 | 5 | 4 | 4 | 4 | 4 | 4 | 4 | 5 | 5 |
| 4 | 4 | 5 | 3 | 4 | 5 | 3 | 4 | 4 | 5 |
| 4 | 5 | 4 | 3 | 4 | 3 | 4 | 4 | 5 | 4 |
| 4 | 4 | 5 | 4 | 3 | 6 | 5 | 6 | 4 | 4 |
| 3 | 5 | 5 | 4 | 3 | 3 | 5 | 3 | 4 | 5 |
| 4 | 4 | 6 | 5 | 4 | 3 | 5 | 4 | 4 | 5 |
| 4 | 4 | 4 | 5 | 3 | 4 | 4 | 4 | 5 | 4 |
| 5 | 4 | 5 | 6 | 3 | 4 | 4 | 4 | 3 | 3 |
| 5 | 4 | 3 | 3 | 5 | 5 | 3 | 4 | 5 | 4 |
| 5 | 4 | 5 | 5 | 5 | 4 | 5 | 3 | 5 | 5 |
| 5 | 3 | 5 | 4 | 6 | 3 | 4 | 4 | 4 | 5 |
| 6 | 3 | 4 | 5 | 5 | 3 | 6 | 4 | 3 | 5 |
| 4 | 3 | 6 | 4 | 3 | 4 | 4 | 4 | 3 | 3 |
| 4 | 4 | 4 | 6 | 5 | 5 | 5 | 3 | 5 | 5 |
| 3 | 4 | 4 | 4 | 4 | 4 | 4 | 5 | 4 | 5 |
| 5 | 4 | 3 | 4 | 4 | 6 | 4 | 6 | 4 | 5 |

| | | | | | | | | | |
|---|---|---|---|---|---|---|---|---|---|
| **4** | **5** | **4** | **4** | **3** | **4** | **4** | **4** | **6** | **6** |
| **5** | 5 | 3 | 3 | 4 | 3 | 3 | 4 | 4 | 4 |
| **5** | 6 | 4 | 4 | 4 | 5 | 4 | 5 | 4 | 6 |
| **4** | 4 | 5 | 4 | 4 | 4 | 4 | 6 | 4 | 4 |
| **4** | 5 | 4 | 3 | 6 | 5 | 4 | 5 | 5 | 4 |
| **5** | 4 | 4 | 6 | 3 | 3 | 3 | 4 | 3 | 4 |
| **3** | 6 | 4 | 3 | 5 | 4 | 5 | 3 | 4 | 3 |
| **4** | 5 | 6 | 5 | 5 | 4 | 5 | 4 | 4 | 3 |
| **4** | 5 | 4 | 5 | 4 | 5 | 3 | 5 | 4 | 5 |
| **4** | 4 | 5 | 4 | 6 | 4 | 4 | 5 | 4 | 4 |
| **4** | 3 | 4 | 5 | 4 | 4 | 4 | 5 | 3 | 5 |
| **3** | 4 | 4 | 5 | 3 | 5 | 4 | 5 | 5 | 6 |
| **4** | 4 | 4 | 3 | 3 | 3 | 4 | 5 | 5 | 5 |
| **3** | 5 | 5 | 6 | 4 | 4 | 3 | 5 | 3 | 4 |
| **4** | 6 | 3 | 4 | 5 | 3 | 5 | 5 | 4 | 6 |
| **3** | 5 | 6 | 3 | 4 | 3 | 5 | 3 | 4 | 5 |
| **3** | 3 | 5 | 6 | 3 | 4 | 6 | 4 | 5 | 5 |
| **5** | 4 | 3 | 5 | 4 | 6 | 4 | 3 | 4 | 4 |
| **3** | 4 | 5 | 5 | 4 | 4 | 5 | 4 | 4 | 3 |
| **4** | 4 | 5 | 4 | 3 | 3 | 4 | 5 | 6 | 4 |
| **4** | 4 | 3 | 4 | 5 | 4 | 5 | 3 | 3 | 4 |
| **3** | 4 | 4 | 3 | 5 | 4 | 5 | 4 | 3 | 5 |
| **4** | 5 | 4 | 4 | 3 | 4 | 4 | 4 | 4 | 4 |
| **4** | 5 | 3 | 3 | 4 | 4 | 4 | 5 | 4 | 3 |
| **5** | 4 | 5 | 4 | 3 | 4 | 4 | 4 | 4 | 4 |
| **5** | 4 | 3 | 4 | 4 | 4 | 4 | 4 | 4 | 4 |
| **5** | 3 | 5 | 3 | 4 | 3 | 4 | 4 | 3 | 3 |
| **4** | 6 | 4 | 4 | 3 | 4 | 5 | 5 | 3 | 5 |
| **3** | 4 | 3 | 4 | 3 | 3 | 4 | 3 | 5 | 5 |
| **5** | 4 | 3 | 3 | 4 | 3 | 4 | 4 | 4 | 5 |
| **5** | 4 | 3 | 3 | 5 | 3 | 4 | 5 | 5 | 5 |
| **4** | 5 | 4 | 4 | 5 | 3 | 4 | 5 | 4 | 4 |
| **5** | 5 | 3 | 4 | 5 | 4 | 4 | 3 | 4 | 3 |
| **5** | 3 | 5 | 3 | 5 | 4 | 6 | 5 | 4 | 5 |
| **5** | 3 | 3 | 4 | 4 | 5 | 3 | 3 | 4 | 3 |
| **3** | 5 | 4 | 5 | 3 | 4 | 4 | 4 | 6 | 5 |
| **4** | 5 | 4 | 4 | 5 | 5 | 5 | 3 | 4 | 4 |
| **4** | 4 | 4 | 5 | 5 | 5 | 5 | 4 | 5 | 5 |
| **5** | 5 | 4 | 5 | 4 | 5 | 4 | 5 | 4 | 4 |
| **3** | 4 | 5 | 5 | 4 | 4 | 4 | 4 | 3 | 4 |
| **5** | 5 | 5 | 5 | 3 | 4 | 5 | 4 | 4 | 5 |
| **4** | 5 | 4 | 4 | 4 | 4 | 5 | 3 | 4 | 4 |
| **4** | 4 | 4 | 3 | 3 | 4 | 4 | 4 | 3 | 4 |
| **4** | 3 | 3 | 4 | 3 | 4 | 4 | 4 | 5 | 4 |
| **5** | 5 | 4 | 5 | 5 | 5 | 6 | 4 | 4 | 4 |
| **4** | 5 | 5 | 5 | 4 | 4 | 5 | 6 | 4 | 6 |
| **5** | 3 | 6 | 5 | 4 | 4 | 4 | 3 | 5 | 4 |
| **4** | 4 | 5 | 3 | 4 | 3 | 4 | 5 | 4 | 4 |
| **5** | 4 | 4 | 4 | 4 | 4 | 3 | 3 | 3 | 4 |
| **4** | 4 | 4 | 4 | 3 | 4 | 4 | 5 | 5 | 4 |
| **6** | 4 | 3 | 5 | 4 | 5 | 4 | 4 | 4 | 5 |
| **4** | 4 | 3 | 6 | 5 | 4 | 4 | 4 | 4 | 4 |
| **3** | 4 | 3 | 5 | 4 | 5 | 4 | 4 | 4 | 5 |
| **4** | 5 | 5 | 4 | 5 | 6 | 5 | 3 | 5 | 5 |

Pup survival to 3-9 months normal, long right tail, mean 0.2 (0.05-0.72)

| **0.22** | **0.26** | **0.26** | **0.11** | **0.45** | **0.15** | **0.21** | **0.16** | **0.12** | **0.26** |
|---|---|---|---|---|---|---|---|---|---|
| **0.20** | 0.27 | 0.13 | 0.11 | 0.14 | 0.23 | 0.27 | 0.52 | 0.29 | 0.10 |
| **0.06** | 0.03 | 0.38 | 0.09 | 0.09 | 0.25 | 0.34 | 0.03 | 0.06 | 0.16 |
| **0.57** | 0.21 | 0.38 | 0.12 | 0.13 | 0.41 | 0.05 | 0.38 | 0.59 | 0.12 |
| **0.10** | 0.20 | 0.12 | 0.19 | 0.14 | 0.24 | 0.14 | 0.29 | 0.26 | 0.13 |
| **0.15** | 0.09 | 0.24 | 0.17 | 0.18 | 0.09 | 0.23 | 0.11 | 0.18 | 0.39 |
| **0.12** | 0.41 | 0.08 | 0.24 | 0.15 | 0.12 | 0.28 | 0.27 | 0.48 | 0.19 |
| **0.16** | 0.24 | 0.30 | 0.14 | 0.20 | 0.17 | 0.18 | 0.18 | 0.06 | 0.13 |
| **0.27** | 0.16 | 0.18 | 0.29 | 0.13 | 0.30 | 0.21 | 0.24 | 0.15 | 0.11 |
| **0.19** | 0.08 | 0.56 | 0.15 | 0.15 | 0.26 | 0.24 | 0.18 | 0.39 | 0.23 |
| **0.15** | 0.14 | 0.17 | 0.17 | 0.16 | 0.53 | 0.43 | 0.31 | 0.17 | 0.18 |
| **0.02** | 0.56 | 0.13 | 0.16 | 0.26 | 0.10 | 0.20 | 0.15 | 0.20 | 0.53 |
| **0.35** | 0.53 | 0.12 | 0.23 | 0.19 | 0.01 | 0.39 | 0.64 | 0.14 | 0.23 |
| **0.16** | 0.33 | 0.25 | 0.24 | 0.06 | 0.19 | 0.15 | 0.50 | 0.31 | 0.34 |
| **0.07** | 0.06 | 0.11 | 0.21 | 0.06 | 0.11 | 0.55 | 0.39 | 0.59 | 0.05 |
| **0.14** | 0.02 | 0.15 | 0.15 | 0.23 | 0.06 | 0.23 | 0.13 | 0.23 | 0.32 |
| **0.18** | 0.08 | 0.55 | 0.18 | 0.30 | 0.16 | 0.29 | 0.03 | 0.13 | 0.16 |
| **0.17** | 0.25 | 0.11 | 0.18 | 0.16 | 0.65 | 0.11 | 0.24 | 0.09 | 0.23 |
| **0.16** | 0.11 | 0.22 | 0.10 | 0.56 | 0.27 | 0.03 | 0.20 | 0.18 | 0.11 |
| **0.20** | 0.19 | 0.23 | 0.26 | 0.04 | 0.11 | 0.18 | 0.53 | 0.16 | 0.23 |
| **0.52** | 0.23 | 0.15 | 0.27 | 0.22 | 0.19 | 0.25 | 0.14 | 0.25 | 0.18 |
| **0.22** | 0.20 | 0.25 | 0.17 | 0.13 | 0.05 | 0.13 | 0.09 | 0.23 | 0.16 |
| **0.16** | 0.51 | 0.22 | 0.18 | 0.52 | 0.16 | 0.12 | 0.18 | 0.16 | 0.13 |
| **0.13** | 0.25 | 0.24 | 0.12 | 0.22 | 0.47 | 0.30 | 0.54 | 0.20 | 0.13 |
| **0.17** | 0.03 | 0.54 | 0.12 | 0.16 | 0.11 | 0.10 | 0.26 | 0.24 | 0.20 |
| **0.13** | 0.18 | 0.14 | 0.16 | 0.13 | 0.17 | 0.29 | 0.05 | 0.52 | 0.11 |
| **0.06** | 0.06 | 0.16 | 0.18 | 0.19 | 0.14 | 0.13 | 0.16 | 0.37 | 0.17 |
| **0.33** | 0.23 | 0.04 | 0.09 | 0.17 | 0.33 | 0.12 | 0.23 | 0.28 | 0.32 |
| **0.44** | 0.21 | 0.25 | 0.15 | 0.42 | 0.17 | 0.49 | 0.23 | 0.27 | 0.33 |
| **0.30** | 0.16 | 0.17 | 0.55 | 0.15 | 0.13 | 0.19 | 0.07 | 0.34 | 0.13 |
| **0.19** | 0.32 | 0.26 | 0.15 | 0.16 | 0.12 | 0.21 | 0.12 | 0.17 | 0.15 |
| **0.23** | 0.32 | 0.24 | 0.28 | 0.26 | 0.27 | 0.07 | 0.11 | 0.19 | 0.17 |
| **0.14** | 0.54 | 0.50 | 0.33 | 0.22 | 0.20 | 0.22 | 0.24 | 0.58 | 0.23 |
| **0.11** | 0.17 | 0.17 | 0.06 | 0.14 | 0.13 | 0.17 | 0.17 | 0.01 | 0.27 |
| **0.01** | 0.15 | 0.13 | 0.08 | 0.13 | 0.07 | 0.02 | 0.17 | 0.03 | 0.12 |
| **0.28** | 0.44 | 0.48 | 0.06 | 0.26 | 0.18 | 0.13 | 0.18 | 0.06 | 0.21 |
| **0.02** | 0.18 | 0.49 | 0.18 | 0.25 | 0.21 | 0.31 | 0.27 | 0.10 | 0.05 |
| **0.24** | 0.37 | 0.05 | 0.12 | 0.12 | 0.58 | 0.25 | 0.42 | 0.28 | 0.19 |
| **0.25** | 0.47 | 0.20 | 0.25 | 0.17 | 0.19 | 0.22 | 0.24 | 0.02 | 0.45 |
| **0.13** | 0.27 | 0.16 | 0.56 | 0.13 | 0.20 | 0.21 | 0.19 | 0.22 | 0.26 |
| **0.15** | 0.31 | 0.28 | 0.15 | 0.11 | 0.13 | 0.13 | 0.10 | 0.14 | 0.51 |
| **0.10** | 0.49 | 0.25 | 0.04 | 0.46 | 0.25 | 0.12 | 0.14 | 0.01 | 0.41 |
| **0.17** | 0.31 | 0.20 | 0.26 | 0.23 | 0.28 | 0.15 | 0.15 | 0.41 | 0.13 |
| **0.40** | 0.23 | 0.04 | 0.12 | 0.10 | 0.17 | 0.17 | 0.21 | 0.31 | 0.67 |
| **0.11** | 0.15 | 0.10 | 0.47 | 0.26 | 0.04 | 0.51 | 0.05 | 0.12 | 0.56 |
| **0.22** | 0.62 | 0.26 | 0.16 | 0.09 | 0.12 | 0.16 | 0.19 | 0.21 | 0.11 |
| **0.12** | 0.13 | 0.16 | 0.23 | 0.05 | 0.08 | 0.15 | 0.16 | 0.27 | 0.15 |
| **0.01** | 0.21 | 0.05 | 0.11 | 0.18 | 0.27 | 0.14 | 0.43 | 0.10 | 0.21 |
| **0.23** | 0.24 | 0.19 | 0.21 | 0.23 | 0.14 | 0.39 | 0.52 | 0.18 | 0.35 |
| **0.43** | 0.13 | 0.22 | 0.00 | 0.21 | 0.32 | 0.10 | 0.25 | 0.13 | 0.24 |
| **0.27** | 0.11 | 0.30 | 0.23 | 0.06 | 0.22 | 0.13 | 0.01 | 0.24 | 0.55 |
| **0.13** | 0.58 | 0.12 | 0.13 | 0.38 | 0.17 | 0.34 | 0.29 | 0.29 | 0.60 |
| **0.59** | 0.15 | 0.39 | 0.30 | 0.45 | 0.17 | 0.34 | 0.03 | 0.23 | 0.15 |
| **0.13** | 0.23 | 0.17 | 0.18 | 0.06 | 0.10 | 0.16 | 0.26 | 0.27 | 0.28 |
| **0.35** | 0.01 | 0.03 | 0.09 | 0.18 | 0.17 | 0.28 | 0.13 | 0.04 | 0.21 |
| **0.30** | 0.25 | 0.18 | 0.16 | 0.12 | 0.18 | 0.24 | 0.27 | 0.60 | 0.38 |
| **0.63** | 0.20 | 0.24 | 0.24 | 0.13 | 0.09 | 0.64 | 0.22 | 0.24 | 0.19 |
| **0.13** | 0.14 | 0.47 | 0.15 | 0.27 | 0.10 | 0.08 | 0.43 | 0.17 | 0.17 |
| **0.16** | 0.21 | 0.16 | 0.08 | 0.10 | 0.48 | 0.28 | 0.59 | 0.10 | 0.09 |
| **0.26** | 0.25 | 0.48 | 0.16 | 0.22 | 0.14 | 0.17 | 0.25 | 0.15 | 0.16 |
| **0.13** | 0.08 | 0.22 | 0.11 | 0.13 | 0.06 | 0.14 | 0.20 | 0.36 | 0.18 |
| **0.08** | 0.16 | 0.59 | 0.35 | 0.15 | 0.26 | 0.08 | 0.18 | 0.24 | 0.16 |
| **0.16** | 0.28 | 0.19 | 0.20 | 0.03 | 0.21 | 0.14 | 0.15 | 0.17 | 0.16 |
| **0.15** | 0.02 | 0.19 | 0.22 | 0.15 | 0.17 | 0.05 | 0.26 | 0.13 | 0.12 |
| **0.42** | 0.17 | 0.04 | 0.28 | 0.12 | 0.12 | 0.20 | 0.14 | 0.14 | 0.18 |
| **0.09** | 0.08 | 0.42 | 0.25 | 0.06 | 0.15 | 0.24 | 0.62 | 0.15 | 0.15 |
| **0.17** | 0.25 | 0.11 | 0.22 | 0.08 | 0.15 | 0.44 | 0.12 | 0.04 | 0.17 |

| **0.22** | **0.26** | **0.26** | **0.11** | **0.45** | **0.15** | **0.21** | **0.16** | **0.12** | **0.26** |
|---|---|---|---|---|---|---|---|---|---|
| **0.07** | 0.22 | 0.22 | 0.57 | 0.12 | 0.63 | 0.12 | 0.06 | 0.12 | 0.21 |
| **0.15** | 0.28 | 0.07 | 0.06 | 0.56 | 0.54 | 0.24 | 0.17 | 0.18 | 0.29 |
| **0.26** | 0.52 | 0.24 | 0.28 | 0.20 | 0.03 | 0.26 | 0.18 | 0.03 | 0.30 |
| **0.28** | 0.29 | 0.31 | 0.31 | 0.16 | 0.22 | 0.10 | 0.07 | 0.25 | 0.23 |
| **0.19** | 0.08 | 0.09 | 0.13 | 0.18 | 0.25 | 0.22 | 0.25 | 0.37 | 0.17 |
| **0.23** | 0.20 | 0.16 | 0.28 | 0.20 | 0.08 | 0.14 | 0.25 | 0.21 | 0.20 |
| **0.09** | 0.25 | 0.28 | 0.10 | 0.30 | 0.57 | 0.27 | 0.27 | 0.16 | 0.22 |
| **0.10** | 0.21 | 0.22 | 0.13 | 0.40 | 0.10 | 0.09 | 0.53 | 0.16 | 0.13 |
| **0.51** | 0.20 | 0.22 | 0.33 | 0.09 | 0.49 | 0.06 | 0.19 | 0.06 | 0.27 |
| **0.25** | 0.58 | 0.16 | 0.09 | 0.06 | 0.20 | 0.01 | 0.19 | 0.10 | 0.10 |
| **0.19** | 0.18 | 0.17 | 0.09 | 0.49 | 0.23 | 0.22 | 0.16 | 0.30 | 0.49 |
| **0.41** | 0.27 | 0.22 | 0.17 | 0.19 | 0.31 | 0.53 | 0.19 | 0.09 | 0.32 |
| **0.13** | 0.49 | 0.18 | 0.35 | 0.11 | 0.19 | 0.15 | 0.09 | 0.26 | 0.11 |
| **0.15** | 0.62 | 0.13 | 0.37 | 0.04 | 0.22 | 0.23 | 0.17 | 0.24 | 0.16 |
| **0.30** | 0.43 | 0.12 | 0.03 | 0.12 | 0.40 | 0.02 | 0.35 | 0.19 | 0.14 |
| **0.53** | 0.29 | 0.23 | 0.05 | 0.23 | 0.01 | 0.24 | 0.19 | 0.11 | 0.43 |
| **0.12** | 0.11 | 0.19 | 0.18 | 0.22 | 0.16 | 0.23 | 0.47 | 0.01 | 0.23 |
| **0.15** | 0.25 | 0.29 | 0.16 | 0.01 | 0.21 | 0.21 | 0.17 | 0.17 | 0.10 |
| **0.43** | 0.40 | 0.18 | 0.22 | 0.27 | 0.62 | 0.16 | 0.19 | 0.16 | 0.23 |
| **0.11** | 0.18 | 0.39 | 0.18 | 0.14 | 0.44 | 0.36 | 0.17 | 0.44 | 0.25 |
| **0.32** | 0.17 | 0.53 | 0.12 | 0.23 | 0.20 | 0.17 | 0.20 | 0.22 | 0.15 |
| **0.26** | 0.10 | 0.19 | 0.44 | 0.16 | 0.20 | 0.05 | 0.20 | 0.22 | 0.51 |
| **0.13** | 0.50 | 0.29 | 0.16 | 0.41 | 0.19 | 0.23 | 0.27 | 0.17 | 0.19 |
| **0.27** | 0.23 | 0.18 | 0.15 | 0.12 | 0.27 | 0.19 | 0.28 | 0.10 | 0.29 |
| **0.18** | 0.10 | 0.15 | 0.14 | 0.40 | 0.09 | 0.05 | 0.17 | 0.32 | 0.14 |
| **0.13** | 0.08 | 0.25 | 0.66 | 0.21 | 0.26 | 0.05 | 0.24 | 0.09 | 0.45 |
| **0.14** | 0.01 | 0.30 | 0.05 | 0.23 | 0.24 | 0.04 | 0.26 | 0.19 | 0.16 |
| **0.07** | 0.11 | 0.23 | 0.14 | 0.17 | 0.23 | 0.23 | 0.24 | 0.14 | 0.44 |
| **0.22** | 0.48 | 0.34 | 0.11 | 0.20 | 0.18 | 0.23 | 0.24 | 0.13 | 0.18 |
| **0.21** | 0.13 | 0.13 | 0.43 | 0.18 | 0.14 | 0.47 | 0.26 | 0.16 | 0.28 |
| **0.12** | 0.18 | 0.13 | 0.15 | 0.23 | 0.16 | 0.15 | 0.25 | 0.08 | 0.37 |
| **0.35** | 0.15 | 0.01 | 0.18 | 0.16 | 0.16 | 0.54 | 0.08 | 0.04 | 0.16 |
| **0.11** | 0.28 | 0.50 | 0.06 | 0.12 | 0.23 | 0.05 | 0.13 | 0.26 | 0.57 |
| **0.21** | 0.47 | 0.01 | 0.11 | 0.22 | 0.14 | 0.24 | 0.16 | 0.31 | 0.03 |
| **0.10** | 0.15 | 0.17 | 0.10 | 0.16 | 0.31 | 0.25 | 0.06 | 0.05 | 0.04 |
| **0.21** | 0.11 | 0.15 | 0.42 | 0.22 | 0.25 | 0.23 | 0.13 | 0.13 | 0.12 |
| **0.11** | 0.06 | 0.16 | 0.42 | 0.04 | 0.26 | 0.14 | 0.26 | 0.21 | 0.11 |
| **0.16** | 0.26 | 0.12 | 0.39 | 0.12 | 0.40 | 0.52 | 0.35 | 0.13 | 0.42 |
| **0.57** | 0.53 | 0.11 | 0.48 | 0.07 | 0.20 | 0.54 | 0.16 | 0.16 | 0.35 |
| **0.16** | 0.17 | 0.52 | 0.34 | 0.23 | 0.10 | 0.27 | 0.08 | 0.11 | 0.18 |
| **0.25** | 0.09 | 0.08 | 0.09 | 0.07 | 0.21 | 0.02 | 0.18 | 0.07 | 0.30 |
| **0.46** | 0.24 | 0.54 | 0.22 | 0.51 | 0.42 | 0.24 | 0.30 | 0.31 | 0.21 |
| **0.10** | 0.07 | 0.27 | 0.30 | 0.09 | 0.11 | 0.08 | 0.30 | 0.45 | 0.21 |
| **0.13** | 0.28 | 0.14 | 0.16 | 0.32 | 0.13 | 0.23 | 0.13 | 0.11 | 0.07 |
| **0.15** | 0.06 | 0.25 | 0.12 | 0.31 | 0.12 | 0.39 | 0.17 | 0.09 | 0.31 |
| **0.08** | 0.09 | 0.21 | 0.16 | 0.11 | 0.17 | 0.23 | 0.29 | 0.16 | 0.06 |
| **0.17** | 0.19 | 0.12 | 0.18 | 0.10 | 0.45 | 0.11 | 0.39 | 0.02 | 0.23 |
| **0.22** | 0.58 | 0.05 | 0.19 | 0.11 | 0.16 | 0.20 | 0.16 | 0.06 | 0.18 |
| **0.32** | 0.16 | 0.15 | 0.40 | 0.11 | 0.20 | 0.28 | 0.14 | 0.05 | 0.16 |
| **0.34** | 0.20 | 0.17 | 0.16 | 0.25 | 0.22 | 0.28 | 0.18 | 0.17 | 0.51 |
| **0.24** | 0.50 | 0.23 | 0.23 | 0.20 | 0.25 | 0.25 | 0.43 | 0.54 | 0.52 |
| **0.17** | 0.18 | 0.17 | 0.17 | 0.07 | 0.20 | 0.18 | 0.13 | 0.16 | 0.12 |
| **0.32** | 0.03 | 0.11 | 0.20 | 0.23 | 0.18 | 0.26 | 0.15 | 0.30 | 0.19 |

Number of breeding packs 74-167 uniform random

| | | | | | | | | | |
|---|---|---|---|---|---|---|---|---|---|
| **125** | **95** | **109** | **167** | **157** | **138** | **75** | **114** | **100** | **99** |
| **155** | 121 | 159 | 154 | 141 | 120 | 88 | 104 | 122 | 100 |
| **165** | 121 | 144 | 122 | 152 | 157 | 75 | 77 | 79 | 132 |
| **119** | 76 | 134 | 80 | 79 | 86 | 166 | 125 | 162 | 98 |
| **74** | 163 | 148 | 149 | 114 | 136 | 78 | 105 | 86 | 144 |
| **132** | 134 | 134 | 136 | 160 | 130 | 135 | 80 | 87 | 101 |
| **127** | 126 | 107 | 152 | 157 | 139 | 101 | 141 | 140 | 90 |
| **153** | 100 | 160 | 139 | 100 | 116 | 155 | 124 | 139 | 107 |
| **108** | 166 | 114 | 165 | 165 | 149 | 79 | 105 | 134 | 109 |
| **163** | 113 | 102 | 135 | 89 | 129 | 152 | 148 | 149 | 145 |
| **94** | 101 | 82 | 164 | 91 | 115 | 108 | 82 | 82 | 80 |
| **138** | 147 | 158 | 75 | 105 | 87 | 131 | 116 | 144 | 143 |
| **120** | 160 | 149 | 109 | 130 | 166 | 93 | 92 | 119 | 160 |
| **129** | 85 | 86 | 133 | 131 | 128 | 165 | 93 | 144 | 136 |
| **126** | 138 | 130 | 156 | 160 | 143 | 131 | 155 | 149 | 145 |
| **121** | 125 | 91 | 111 | 133 | 152 | 140 | 79 | 152 | 143 |
| **143** | 77 | 85 | 141 | 132 | 116 | 96 | 129 | 92 | 162 |
| **137** | 90 | 153 | 111 | 88 | 149 | 140 | 119 | 137 | 111 |
| **151** | 137 | 115 | 166 | 120 | 109 | 84 | 78 | 95 | 148 |
| **108** | 155 | 86 | 91 | 101 | 128 | 87 | 106 | 115 | 120 |
| **101** | 156 | 143 | 157 | 75 | 82 | 114 | 143 | 123 | 165 |
| **89** | 117 | 160 | 153 | 140 | 89 | 140 | 97 | 115 | 119 |
| **120** | 111 | 139 | 101 | 79 | 107 | 108 | 145 | 83 | 109 |
| **119** | 141 | 166 | 75 | 145 | 153 | 83 | 162 | 127 | 151 |
| **112** | 86 | 105 | 96 | 79 | 76 | 140 | 129 | 129 | 132 |
| **121** | 96 | 108 | 97 | 95 | 74 | 161 | 85 | 81 | 125 |
| **114** | 135 | 91 | 138 | 137 | 106 | 78 | 116 | 107 | 86 |
| **81** | 83 | 80 | 150 | 142 | 158 | 82 | 139 | 124 | 167 |
| **162** | 108 | 80 | 153 | 155 | 128 | 138 | 149 | 126 | 135 |
| **165** | 152 | 155 | 130 | 120 | 167 | 131 | 138 | 128 | 107 |
| **85** | 161 | 166 | 115 | 108 | 110 | 93 | 101 | 132 | 153 |
| **146** | 77 | 101 | 143 | 116 | 83 | 136 | 152 | 102 | 97 |
| **143** | 143 | 151 | 148 | 80 | 96 | 132 | 100 | 119 | 118 |
| **130** | 117 | 91 | 83 | 120 | 99 | 131 | 109 | 118 | 125 |
| **134** | 117 | 105 | 134 | 85 | 133 | 113 | 142 | 167 | 134 |
| **153** | 115 | 120 | 86 | 135 | 107 | 135 | 115 | 122 | 157 |
| **117** | 156 | 126 | 89 | 89 | 161 | 159 | 82 | 144 | 74 |
| **107** | 112 | 123 | 140 | 92 | 80 | 143 | 104 | 145 | 105 |
| **80** | 81 | 159 | 167 | 82 | 151 | 150 | 116 | 76 | 139 |
| **103** | 154 | 76 | 83 | 77 | 166 | 77 | 95 | 163 | 155 |
| **111** | 130 | 151 | 148 | 82 | 83 | 125 | 78 | 79 | 142 |
| **118** | 101 | 127 | 135 | 93 | 133 | 167 | 115 | 122 | 164 |
| **163** | 121 | 163 | 145 | 111 | 166 | 126 | 135 | 113 | 127 |
| **105** | 157 | 83 | 153 | 120 | 155 | 167 | 106 | 109 | 74 |
| **156** | 131 | 166 | 88 | 77 | 89 | 98 | 158 | 121 | 96 |
| **127** | 112 | 86 | 78 | 127 | 111 | 133 | 164 | 101 | 110 |
| **94** | 107 | 145 | 89 | 130 | 146 | 152 | 131 | 76 | 116 |
| **137** | 120 | 74 | 117 | 90 | 128 | 84 | 93 | 151 | 140 |
| **107** | 152 | 83 | 127 | 97 | 124 | 87 | 142 | 148 | 121 |
| **155** | 87 | 119 | 119 | 159 | 142 | 135 | 76 | 92 | 86 |
| **129** | 122 | 111 | 75 | 80 | 90 | 91 | 148 | 99 | 148 |
| **74** | 147 | 146 | 127 | 85 | 115 | 136 | 113 | 167 | 145 |
| **105** | 124 | 81 | 152 | 85 | 110 | 82 | 130 | 138 | 95 |
| **162** | 105 | 156 | 128 | 127 | 134 | 132 | 124 | 133 | 152 |
| **117** | 156 | 82 | 164 | 158 | 150 | 74 | 152 | 83 | 112 |
| **124** | 159 | 124 | 151 | 146 | 159 | 78 | 85 | 76 | 130 |
| **109** | 99 | 93 | 130 | 120 | 86 | 102 | 120 | 97 | 108 |
| **125** | 91 | 98 | 107 | 158 | 113 | 82 | 159 | 110 | 110 |
| **145** | 98 | 117 | 129 | 92 | 161 | 89 | 119 | 158 | 147 |
| **125** | 134 | 114 | 139 | 118 | 111 | 107 | 78 | 81 | 120 |
| **107** | 109 | 129 | 148 | 121 | 90 | 166 | 89 | 142 | 104 |
| **107** | 145 | 112 | 159 | 148 | 91 | 93 | 165 | 101 | 151 |
| **103** | 146 | 80 | 86 | 150 | 123 | 141 | 114 | 83 | 81 |
| **143** | 125 | 145 | 98 | 111 | 92 | 77 | 136 | 135 | 120 |
| **106** | 79 | 127 | 167 | 114 | 123 | 118 | 100 | 156 | 92 |
| **109** | 125 | 151 | 167 | 115 | 131 | 77 | 113 | 97 | 111 |
| **133** | 87 | 151 | 124 | 120 | 149 | 93 | 135 | 80 | 154 |

| | | | | | | | | | |
|---|---|---|---|---|---|---|---|---|---|
| **125** | **95** | **109** | **167** | **157** | **138** | **75** | **114** | **100** | **99** |
| **123** | 139 | 119 | 87 | 105 | 141 | 77 | 82 | 141 | 106 |
| **91** | 155 | 104 | 112 | 94 | 92 | 80 | 88 | 89 | 131 |
| **125** | 162 | 80 | 151 | 144 | 74 | 101 | 122 | 109 | 86 |
| **77** | 163 | 79 | 122 | 113 | 162 | 136 | 120 | 131 | 135 |
| **133** | 80 | 112 | 124 | 132 | 135 | 142 | 112 | 75 | 136 |
| **87** | 95 | 121 | 128 | 145 | 94 | 144 | 121 | 151 | 158 |
| **133** | 75 | 155 | 100 | 88 | 165 | 130 | 152 | 122 | 131 |
| **91** | 149 | 120 | 120 | 108 | 127 | 166 | 77 | 139 | 160 |
| **134** | 153 | 132 | 129 | 87 | 115 | 96 | 108 | 157 | 84 |
| **129** | 159 | 164 | 92 | 77 | 157 | 131 | 91 | 146 | 75 |
| **107** | 86 | 140 | 148 | 108 | 109 | 148 | 162 | 153 | 98 |
| **127** | 87 | 104 | 106 | 101 | 133 | 78 | 136 | 116 | 119 |
| **109** | 143 | 144 | 76 | 131 | 135 | 99 | 112 | 150 | 101 |
| **120** | 153 | 132 | 161 | 99 | 85 | 117 | 155 | 155 | 108 |
| **102** | 90 | 115 | 92 | 79 | 152 | 157 | 139 | 81 | 125 |
| **86** | 80 | 77 | 140 | 141 | 76 | 153 | 101 | 155 | 93 |
| **131** | 163 | 98 | 151 | 153 | 115 | 127 | 132 | 147 | 101 |
| **101** | 166 | 118 | 148 | 122 | 92 | 105 | 81 | 107 | 136 |
| **161** | 144 | 80 | 90 | 108 | 165 | 135 | 136 | 96 | 144 |
| **78** | 74 | 150 | 153 | 83 | 155 | 79 | 136 | 161 | 126 |
| **82** | 77 | 135 | 147 | 128 | 117 | 159 | 82 | 131 | 145 |
| **116** | 121 | 134 | 136 | 144 | 83 | 143 | 78 | 117 | 120 |
| **77** | 77 | 143 | 106 | 153 | 114 | 161 | 107 | 77 | 127 |
| **129** | 109 | 79 | 120 | 162 | 140 | 161 | 160 | 113 | 80 |
| **139** | 121 | 146 | 122 | 127 | 100 | 114 | 146 | 151 | 105 |
| **109** | 84 | 92 | 109 | 139 | 139 | 111 | 82 | 141 | 149 |
| **114** | 156 | 79 | 128 | 91 | 93 | 118 | 85 | 105 | 116 |
| **141** | 164 | 156 | 75 | 115 | 167 | 158 | 124 | 135 | 157 |
| **119** | 136 | 116 | 157 | 150 | 77 | 86 | 148 | 147 | 132 |
| **97** | 161 | 165 | 111 | 81 | 147 | 152 | 98 | 167 | 98 |
| **165** | 81 | 91 | 132 | 93 | 128 | 159 | 151 | 91 | 126 |
| **75** | 93 | 86 | 110 | 134 | 95 | 163 | 148 | 77 | 114 |
| **152** | 161 | 166 | 119 | 102 | 134 | 140 | 152 | 81 | 81 |
| **144** | 149 | 142 | 143 | 78 | 133 | 88 | 76 | 99 | 78 |
| **159** | 141 | 101 | 80 | 105 | 123 | 151 | 124 | 136 | 111 |
| **156** | 123 | 116 | 126 | 131 | 117 | 95 | 159 | 79 | 109 |
| **121** | 162 | 80 | 79 | 85 | 152 | 129 | 94 | 163 | 130 |
| **163** | 105 | 159 | 135 | 166 | 98 | 126 | 112 | 87 | 140 |
| **105** | 166 | 156 | 112 | 165 | 127 | 98 | 153 | 99 | 150 |
| **75** | 134 | 91 | 127 | 132 | 129 | 122 | 125 | 147 | 83 |
| **119** | 135 | 107 | 151 | 76 | 110 | 125 | 139 | 117 | 118 |
| **147** | 167 | 78 | 82 | 80 | 83 | 93 | 164 | 117 | 114 |
| **75** | 120 | 139 | 158 | 161 | 158 | 114 | 122 | 123 | 126 |
| **162** | 111 | 154 | 117 | 148 | 100 | 158 | 165 | 150 | 131 |
| **120** | 138 | 119 | 146 | 79 | 78 | 105 | 139 | 126 | 127 |
| **140** | 149 | 104 | 123 | 110 | 118 | 114 | 128 | 150 | 166 |
| **162** | 137 | 98 | 84 | 89 | 89 | 118 | 126 | 106 | 103 |
| **103** | 140 | 124 | 140 | 127 | 102 | 127 | 125 | 151 | 97 |
| **146** | 157 | 103 | 167 | 153 | 111 | 92 | 146 | 137 | 156 |
| **138** | 104 | 114 | 90 | 98 | 90 | 144 | 77 | 107 | 104 |
| **95** | 115 | 116 | 101 | 118 | 141 | 125 | 108 | 128 | 98 |
| **103** | 150 | 158 | 163 | 78 | 119 | 153 | 130 | 137 | 79 |
| **116** | 143 | 147 | 96 | 123 | 146 | 96 | 98 | 105 | 99 |

N2021 traditional =695-751 uniform

| 704 | 723 | 731 | 721 | 750 | 743 | 723 | 738 | 741 | 731 |
|---|---|---|---|---|---|---|---|---|---|
| 697 | 701 | 728 | 728 | 748 | 719 | 719 | 708 | 738 | 740 |
| 715 | 700 | 710 | 715 | 748 | 718 | 750 | 742 | 743 | 747 |
| 738 | 746 | 718 | 710 | 706 | 751 | 713 | 720 | 701 | 735 |
| 741 | 747 | 708 | 737 | 742 | 703 | 746 | 695 | 702 | 725 |
| 734 | 741 | 709 | 721 | 724 | 741 | 703 | 721 | 711 | 721 |
| 751 | 742 | 705 | 735 | 714 | 740 | 716 | 698 | 729 | 728 |
| 699 | 737 | 730 | 704 | 728 | 749 | 734 | 712 | 742 | 745 |
| 749 | 696 | 720 | 720 | 721 | 702 | 725 | 702 | 731 | 741 |
| 729 | 743 | 751 | 708 | 740 | 726 | 699 | 729 | 704 | 741 |
| 735 | 723 | 708 | 745 | 721 | 720 | 719 | 711 | 728 | 701 |
| 704 | 715 | 747 | 732 | 734 | 740 | 751 | 730 | 730 | 732 |
| 742 | 731 | 713 | 702 | 701 | 703 | 748 | 751 | 718 | 741 |
| 714 | 719 | 723 | 737 | 724 | 726 | 729 | 715 | 712 | 721 |
| 708 | 732 | 711 | 705 | 699 | 724 | 744 | 706 | 695 | 722 |
| 706 | 728 | 720 | 738 | 745 | 749 | 713 | 721 | 723 | 748 |
| 727 | 709 | 747 | 722 | 719 | 729 | 703 | 714 | 750 | 730 |
| 729 | 701 | 738 | 747 | 727 | 701 | 704 | 717 | 734 | 732 |
| 723 | 719 | 741 | 751 | 751 | 697 | 738 | 744 | 715 | 749 |
| 731 | 724 | 730 | 743 | 751 | 701 | 703 | 730 | 698 | 702 |
| 743 | 719 | 727 | 715 | 715 | 715 | 720 | 732 | 704 | 747 |
| 709 | 723 | 735 | 734 | 749 | 749 | 698 | 717 | 711 | 727 |
| 743 | 729 | 741 | 711 | 702 | 743 | 706 | 730 | 711 | 728 |
| 710 | 710 | 743 | 695 | 732 | 704 | 738 | 738 | 741 | 725 |
| 728 | 742 | 734 | 739 | 706 | 747 | 712 | 732 | 708 | 750 |
| 737 | 719 | 721 | 740 | 702 | 738 | 750 | 727 | 706 | 738 |
| 723 | 751 | 707 | 736 | 749 | 725 | 749 | 738 | 744 | 728 |
| 725 | 699 | 701 | 711 | 748 | 749 | 749 | 733 | 750 | 703 |
| 741 | 698 | 738 | 738 | 695 | 723 | 743 | 713 | 716 | 740 |
| 710 | 744 | 725 | 737 | 747 | 724 | 745 | 733 | 742 | 751 |
| 732 | 717 | 696 | 703 | 695 | 721 | 734 | 717 | 739 | 743 |
| 721 | 730 | 729 | 726 | 746 | 738 | 715 | 728 | 707 | 700 |
| 749 | 705 | 710 | 749 | 726 | 715 | 699 | 742 | 720 | 726 |

| 704 | 723 | 731 | 721 | 750 | 743 | 723 | 738 | 741 | 731 |
|---|---|---|---|---|---|---|---|---|---|
| **739** | 721 | 697 | 698 | 719 | 746 | 710 | 741 | 696 | 724 |
| **747** | 726 | 747 | 714 | 745 | 726 | 722 | 699 | 740 | 751 |
| **727** | 697 | 699 | 721 | 742 | 732 | 706 | 738 | 743 | 726 |
| **723** | 751 | 695 | 739 | 711 | 751 | 703 | 704 | 710 | 705 |
| **719** | 742 | 727 | 720 | 723 | 737 | 698 | 698 | 702 | 703 |
| **726** | 695 | 717 | 712 | 701 | 726 | 704 | 701 | 748 | 741 |
| **730** | 703 | 728 | 730 | 699 | 732 | 727 | 720 | 709 | 739 |
| **729** | 748 | 713 | 695 | 735 | 751 | 730 | 742 | 740 | 739 |
| **749** | 723 | 706 | 715 | 750 | 711 | 721 | 710 | 712 | 735 |
| **729** | 709 | 740 | 697 | 712 | 724 | 725 | 729 | 745 | 712 |
| **704** | 731 | 742 | 729 | 706 | 749 | 709 | 728 | 716 | 750 |
| **741** | 702 | 701 | 709 | 739 | 745 | 750 | 735 | 719 | 748 |
| **725** | 724 | 722 | 717 | 732 | 743 | 737 | 696 | 705 | 716 |
| **701** | 727 | 726 | 710 | 698 | 720 | 738 | 706 | 709 | 729 |
| **711** | 749 | 696 | 727 | 713 | 696 | 703 | 709 | 709 | 738 |
| **734** | 720 | 725 | 711 | 714 | 746 | 703 | 724 | 702 | 734 |
| **700** | 695 | 706 | 716 | 730 | 731 | 739 | 712 | 717 | 737 |
| **717** | 714 | 728 | 741 | 695 | 726 | 731 | 737 | 735 | 738 |
| **715** | 731 | 743 | 709 | 695 | 743 | 731 | 718 | 732 | 728 |
| **723** | 736 | 736 | 737 | 711 | 729 | 727 | 705 | 720 | 699 |
| **742** | 711 | 744 | 718 | 705 | 720 | 709 | 717 | 740 | 708 |
| **743** | 724 | 739 | 722 | 741 | 746 | 716 | 711 | 709 | 737 |
| **701** | 729 | 745 | 727 | 751 | 736 | 717 | 746 | 739 | 723 |
| **719** | 708 | 743 | 714 | 735 | 710 | 719 | 723 | 749 | 746 |
| **727** | 743 | 721 | 733 | 731 | 743 | 746 | 743 | 717 | 712 |
| **743** | 696 | 696 | 708 | 714 | 699 | 748 | 735 | 697 | 738 |
| **707** | 735 | 703 | 745 | 713 | 743 | 715 | 717 | 705 | 701 |
| **721** | 698 | 708 | 721 | 707 | 700 | 697 | 729 | 702 | 735 |
| **728** | 751 | 725 | 709 | 732 | 702 | 696 | 726 | 718 | 702 |
| **699** | 745 | 732 | 732 | 703 | 699 | 710 | 722 | 723 | 737 |
| **702** | 733 | 695 | 733 | 735 | 735 | 700 | 715 | 705 | 716 |
| **722** | 701 | 725 | 723 | 744 | 727 | 695 | 710 | 716 | 695 |
| **703** | 737 | 743 | 725 | 715 | 733 | 721 | 695 | 731 | 731 |

| 704 | 723 | 731 | 721 | 750 | 743 | 723 | 738 | 741 | 731 |
|---|---|---|---|---|---|---|---|---|---|
| **711** | 742 | 747 | 748 | 698 | 748 | 738 | 700 | 713 | 746 |
| **711** | 735 | 703 | 738 | 745 | 736 | 730 | 701 | 726 | 722 |
| **740** | 728 | 729 | 743 | 725 | 735 | 723 | 707 | 703 | 713 |
| **701** | 725 | 739 | 700 | 733 | 707 | 719 | 728 | 710 | 723 |
| **727** | 734 | 705 | 735 | 742 | 750 | 750 | 718 | 698 | 713 |
| **735** | 723 | 731 | 713 | 735 | 714 | 726 | 700 | 717 | 697 |
| **743** | 703 | 729 | 702 | 695 | 731 | 737 | 734 | 718 | 715 |
| **743** | 717 | 747 | 751 | 696 | 698 | 714 | 719 | 707 | 720 |
| **735** | 748 | 711 | 747 | 750 | 734 | 750 | 746 | 703 | 711 |
| **725** | 729 | 721 | 709 | 728 | 745 | 706 | 736 | 750 | 718 |
| **710** | 712 | 705 | 750 | 746 | 701 | 726 | 722 | 703 | 702 |
| **713** | 738 | 703 | 735 | 708 | 697 | 741 | 728 | 734 | 711 |
| **706** | 707 | 719 | 700 | 750 | 696 | 731 | 717 | 733 | 713 |
| **742** | 740 | 711 | 727 | 740 | 719 | 719 | 700 | 709 | 723 |
| **735** | 745 | 710 | 731 | 730 | 736 | 731 | 716 | 713 | 705 |
| **749** | 741 | 715 | 697 | 720 | 746 | 743 | 747 | 710 | 712 |
| **699** | 709 | 708 | 717 | 719 | 702 | 700 | 708 | 723 | 739 |
| **707** | 701 | 733 | 745 | 745 | 737 | 732 | 724 | 714 | 718 |
| **748** | 716 | 751 | 702 | 725 | 720 | 723 | 729 | 733 | 700 |
| **718** | 734 | 711 | 734 | 707 | 702 | 718 | 720 | 738 | 744 |
| **704** | 702 | 713 | 740 | 718 | 698 | 716 | 704 | 698 | 711 |
| **709** | 702 | 740 | 719 | 746 | 720 | 713 | 732 | 700 | 704 |
| **726** | 734 | 713 | 748 | 722 | 741 | 697 | 714 | 707 | 709 |
| **716** | 739 | 732 | 751 | 749 | 728 | 722 | 702 | 697 | 745 |
| **732** | 714 | 712 | 723 | 696 | 740 | 737 | 715 | 730 | 742 |
| **750** | 723 | 746 | 719 | 733 | 716 | 713 | 701 | 744 | 722 |
| **726** | 720 | 707 | 711 | 744 | 732 | 706 | 722 | 738 | 718 |
| **707** | 739 | 695 | 701 | 743 | 747 | 696 | 731 | 717 | 701 |
| **696** | 748 | 735 | 716 | 711 | 727 | 738 | 742 | 713 | 731 |
| **739** | 744 | 706 | 745 | 720 | 699 | 715 | 700 | 740 | 719 |
| **736** | 716 | 725 | 707 | 706 | 730 | 723 | 750 | 702 | 746 |
| **696** | 714 | 709 | 737 | 712 | 707 | 718 | 699 | 736 | 735 |
| **750** | 718 | 723 | 728 | 739 | 751 | 696 | 734 | 735 | 748 |

| **704** | **723** | **731** | **721** | **750** | **743** | **723** | **738** | **741** | **731** |
|---|---|---|---|---|---|---|---|---|---|
| **710** | 696 | 738 | 730 | 699 | 717 | 740 | 748 | 699 | 726 |
| **731** | 708 | 712 | 737 | 732 | 718 | 727 | 736 | 742 | 727 |
| **728** | 732 | 751 | 736 | 718 | 741 | 710 | 720 | 748 | 697 |
| **732** | 745 | 748 | 729 | 730 | 737 | 720 | 697 | 707 | 704 |
| **709** | 726 | 707 | 705 | 732 | 746 | 705 | 733 | 718 | 751 |
| **735** | 732 | 726 | 746 | 696 | 704 | 742 | 701 | 746 | 739 |
| **723** | 695 | 750 | 739 | 733 | 733 | 729 | 724 | 746 | 709 |
| **727** | 732 | 704 | 701 | 704 | 697 | 715 | 716 | 721 | 714 |
| **743** | 743 | 695 | 718 | 740 | 697 | 744 | 712 | 718 | 751 |
| **702** | 724 | 737 | 735 | 734 | 750 | 702 | 710 | 722 | 698 |
| **743** | 711 | 725 | 733 | 705 | 738 | 717 | 722 | 715 | 722 |
| **735** | 709 | 696 | 742 | 750 | 718 | 696 | 707 | 718 | 733 |
| **696** | 726 | 733 | 745 | 733 | 702 | 703 | 724 | 716 | 742 |
| **703** | 724 | 751 | 750 | 706 | 740 | 737 | 737 | 717 | 717 |
| **751** | 699 | 718 | 730 | 728 | 728 | 743 | 697 | 695 | 715 |
| **695** | 746 | 723 | 735 | 719 | 749 | 748 | 698 | 742 | 725 |
| **750** | 713 | 737 | 710 | 720 | 749 | 730 | 709 | 721 | 716 |
| **732** | 712 | 748 | 738 | 749 | 736 | 729 | 739 | 728 | 723 |
| **750** | 708 | 715 | 728 | 713 | 704 | 702 | 740 | 720 | 735 |
| **705** | 723 | 722 | 700 | 702 | 736 | 696 | 734 | 708 | 747 |
| **700** | 721 | 743 | 726 | 710 | 705 | 698 | 731 | 751 | 696 |

N2021new =1195 (957-1573 normal) The 218 killed in Feb 2021 wolf-hunt are deducted in subsequent steps.

| 1605.5 | 1210.5 | 1178.5 | 1328 | 1234.5 | 951 | 1090.5 | 1687.5 | 1388 | 1504 |
|---|---|---|---|---|---|---|---|---|---|
| 933 | 1258.5 | 739 | 824.5 | 1671 | 1396.5 | 1550.5 | 1200.5 | 1230.5 | 1141 |
| 1462 | 1238.5 | 1223 | 1144 | 981.5 | 1417.5 | 830 | 1312.5 | 1169 | 1323.5 |
| 1106.5 | 1160 | 1334.5 | 1658 | 1276.5 | 1610 | 1370.5 | 1750.5 | 1719.5 | 1373.5 |
| 1160.5 | 1574 | 1547 | 1035.5 | 1152.5 | 1273.5 | 1042.5 | 1662.5 | 1282 | 1019.5 |
| 1189 | 1382.5 | 1509 | 1180 | 1596.5 | 1291.5 | 1485 | 1189 | 1386 | 1386 |
| 918.5 | 1373 | 1203 | 1171 | 942 | 879.5 | 1206 | 746 | 1156.5 | 961.5 |
| 1262.5 | 1421.5 | 970 | 1367 | 1293.5 | 917 | 1090.5 | 1041 | 1227 | 1281.5 |
| 1077.5 | 1752 | 1234 | 878 | 1389.5 | 948 | 1260.5 | 1328 | 1667 | 798 |
| 1169 | 809 | 1292.5 | 1009 | 1175 | 977 | 1387.5 | 868.5 | 1479.5 | 1443 |
| 1256 | 1056.5 | 1079.5 | 1603.5 | 1418.5 | 908 | 1178.5 | 1182 | 1368 | 1386 |
| 1681 | 1186.5 | 1069 | 1114 | 680.5 | 910.5 | 984 | 1225 | 1126 | 1409.5 |
| 1396 | 1623 | 1327 | 1148 | 1024 | 1326.5 | 1476.5 | 1365 | 834.5 | 1711.5 |
| 697.5 | 1237.5 | 1074.5 | 1594.5 | 995 | 1160.5 | 1478 | 1150 | 1000.5 | 1422 |
| 1218.5 | 1166 | 791.5 | 1281 | 1050.5 | 1100.5 | 1237 | 1243.5 | 1085.5 | 1575 |
| 1546 | 1340.5 | 1061.5 | 963 | 743 | 1604.5 | 855 | 968.5 | 1008 | 790.5 |
| 1303 | 1076.5 | 1778.5 | 1374 | 1332.5 | 1239 | 1373 | 1158 | 972.5 | 788.5 |
| 1465.5 | 1539.5 | 1452.5 | 1207 | 1169.5 | 1148 | 1060.5 | 1248 | 704.5 | 1139 |
| 599.5 | 1021 | 914 | 1303.5 | 1009.5 | 1135.5 | 1385 | 1461.5 | 1570.5 | 1081.5 |
| 1276 | 1114.5 | 638 | 1050.5 | 1638.5 | 890 | 1183.5 | 1289.5 | 1460.5 | 934.5 |
| 1473.5 | 1260.5 | 1002.5 | 1378 | 906 | 1763.5 | 1497 | 1284 | 1078 | 1493.5 |
| 1432.5 | 1654 | 725 | 951 | 1401.5 | 1709 | 1087.5 | 807 | 1218 | 1618.5 |
| 1209.5 | 640.5 | 1396 | 1062.5 | 961.5 | 1730 | 1236.5 | 1028 | 1228 | 896.5 |
| 1118 | 1625 | 893 | 1088.5 | 1355 | 864.5 | 1692 | 1137 | 798.5 | 897.5 |
| 1527 | 881.5 | 1625 | 1560 | 1134.5 | 1054.5 | 686 | 1408.5 | 1148 | 915 |
| 1536 | 843.5 | 1181.5 | 1581.5 | 1063.5 | 1295 | 649 | 845 | 1005 | 1151 |
| 782 | 1130.5 | 1449.5 | 1240 | 889.5 | 1337.5 | 1423.5 | 1195 | 881.5 | 1467 |
| 1160.5 | 822 | 1437.5 | 862 | 1128 | 1113.5 | 1297 | 1441 | 1206.5 | 1161.5 |
| 860.5 | 1570 | 1194.5 | 1190 | 1650.5 | 1554 | 1691 | 1210 | 1046.5 | 1007 |
| 1281.5 | 1075.5 | 1529.5 | 1439.5 | 1285 | 1113.5 | 1366.5 | 985 | 1222 | 1051.5 |
| 1385 | 1192.5 | 1336 | 1432.5 | 1045 | 1561.5 | 1100.5 | 946 | 1618 | 1566 |
| 1249 | 664 | 1716.5 | 1339 | 1174.5 | 1231 | 1377 | 870.5 | 1351.5 | 1440.5 |
| 1238.5 | 856.5 | 974 | 1224 | 1357 | 1009 | 1522.5 | 979 | 1503 | 933.5 |

| **1605.5** | **1210.5** | **1178.5** | **1328** | **1234.5** | **951** | **1090.5** | **1687.5** | **1388** | **1504** |
|---|---|---|---|---|---|---|---|---|---|
| **957** | 1351 | 1014 | 1408.5 | 671 | 1321 | 1206.5 | 1117 | 1115.5 | 623.5 |
| **1390** | 1159.5 | 1443.5 | 1327.5 | 853 | 898.5 | 1533 | 1350.5 | 1239 | 1493.5 |
| **953.5** | 1275.5 | 1393.5 | 1171.5 | 1007 | 1029.5 | 1399 | 1211.5 | 1328 | 1257 |
| **1346.5** | 1417 | 634.5 | 1572 | 1013 | 949.5 | 1365.5 | 1028.5 | 898 | 1176.5 |
| **1663.5** | 1226.5 | 1233 | 960.5 | 1111 | 778 | 1162.5 | 1153.5 | 1150.5 | 1379 |
| **1155.5** | 1063 | 1464.5 | 693 | 834.5 | 927.5 | 1120 | 1079.5 | 1510.5 | 1010.5 |
| **880** | 958.5 | 831.5 | 930.5 | 1064 | 881.5 | 1330.5 | 1355.5 | 1219 | 1604.5 |
| **1032** | 1571 | 1035.5 | 1507.5 | 1011 | 909.5 | 1374 | 1149.5 | 1119 | 1399 |
| **1291.5** | 1071 | 1432.5 | 1139.5 | 905 | 1356 | 1325 | 1073 | 1415.5 | 1129 |
| **1395.5** | 1443.5 | 1356 | 1071 | 1162 | 1365.5 | 926 | 1271 | 1106 | 1143 |
| **1002** | 1014 | 1294 | 1608 | 1415.5 | 1168.5 | 1228.5 | 1558.5 | 1038.5 | 1385 |
| **1228.5** | 1753 | 1565.5 | 1205 | 830.5 | 812 | 920 | 1074.5 | 1348.5 | 1031.5 |
| **1602.5** | 1452 | 1602.5 | 1217 | 1105.5 | 1192.5 | 1239.5 | 1688.5 | 911 | 973 |
| **1522** | 1263 | 1348.5 | 1622.5 | 1451 | 846 | 1601 | 1387 | 959 | 1499.5 |
| **1238** | 1491.5 | 911.5 | 737 | 610.5 | 1107.5 | 1350.5 | 1053.5 | 1136 | 1454.5 |
| **1222.5** | 1388 | 957 | 905.5 | 1189.5 | 1215 | 886.5 | 1119.5 | 1481.5 | 1093.5 |
| **811** | 1181.5 | 1040 | 1208.5 | 1422.5 | 1226 | 1200.5 | 879 | 1229 | 944.5 |
| **1679** | 833.5 | 1241.5 | 1325.5 | 1061 | 839.5 | 1088 | 914.5 | 1649.5 | 997 |
| **1385** | 1407 | 1334 | 1304 | 1112.5 | 1404 | 1158.5 | 1644.5 | 1362 | 1105.5 |
| **1396** | 1251.5 | 1119 | 1023.5 | 1102 | 1067.5 | 1170 | 1148.5 | 1288.5 | 1793 |
| **1260** | 1254 | 1072.5 | 985.5 | 1088 | 1255.5 | 1325.5 | 1140 | 1250.5 | 1204.5 |
| **1148** | 1096 | 1303 | 1295.5 | 1401.5 | 1305.5 | 761 | 1408.5 | 816.5 | 1103.5 |
| **1279.5** | 1154.5 | 1711 | 1189.5 | 786 | 1254.5 | 1411.5 | 1327.5 | 1065.5 | 1195 |
| **1562.5** | 1247 | 1256 | 1106.5 | 1208 | 1263 | 1339.5 | 1289.5 | 624 | 1493 |
| **944.5** | 1344 | 1737.5 | 1388 | 1026.5 | 880 | 1492.5 | 1085 | 1208.5 | 1596 |
| **1670** | 974 | 925.5 | 1245 | 1489 | 1024 | 1430.5 | 1239 | 1348.5 | 1539.5 |
| **1440.5** | 724 | 1181.5 | 946.5 | 1737 | 1178 | 1187 | 1446 | 961.5 | 809.5 |
| **1003** | 1206 | 1093 | 1331.5 | 1233 | 1313 | 933 | 1082.5 | 1420 | 1140.5 |
| **1031** | 964.5 | 905.5 | 1501.5 | 1297.5 | 656 | 1458.5 | 1388 | 1218 | 1416 |
| **895.5** | 962.5 | 1577.5 | 1482 | 1076 | 1246 | 1380.5 | 829.5 | 984.5 | 1080 |
| **1492.5** | 1200.5 | 1190 | 1576 | 1706.5 | 1017 | 704 | 1489 | 1537 | 865 |
| **845** | 1056 | 1024 | 1054.5 | 799.5 | 1659 | 1228 | 1477.5 | 936 | 1126.5 |
| **1593.5** | 1372.5 | 1457 | 1250 | 873.5 | 1453.5 | 1208.5 | 1595.5 | 1361 | 1148.5 |
| **1251** | 1578.5 | 1103.5 | 1228.5 | 1088 | 1462 | 1371.5 | 802 | 1125 | 1231.5 |

| | | | | | | | | | |
|---|---|---|---|---|---|---|---|---|---|
| **1605.5** | **1210.5** | **1178.5** | **1328** | **1234.5** | **951** | **1090.5** | **1687.5** | **1388** | **1504** |
| **1231** | 934 | 1030 | 1260 | 667 | 1482.5 | 1038 | 1489 | 1158.5 | 1122.5 |
| **1180** | 1241 | 1171 | 1288 | 1362 | 1552.5 | 1366 | 1189 | 1058.5 | 1255.5 |
| **1055** | 1079 | 1714.5 | 882 | 960 | 780 | 1509.5 | 960.5 | 1113.5 | 970 |
| **1295.5** | 1418.5 | 967 | 1242.5 | 1297.5 | 992 | 933 | 1482 | 1549.5 | 1538.5 |
| **1303.5** | 990.5 | 895 | 1368.5 | 669.5 | 1341.5 | 1061 | 1280.5 | 1377.5 | 1419.5 |
| **1051** | 1575.5 | 1395.5 | 1429.5 | 836 | 1086.5 | 1550 | 1352.5 | 1088 | 1612.5 |
| **838.5** | 1625 | 1235 | 1651 | 952.5 | 1341.5 | 1456.5 | 1154.5 | 1083 | 1253 |
| **1757.5** | 773.5 | 1678.5 | 1286 | 1661.5 | 774 | 1734.5 | 931 | 1317 | 1261 |
| **1234** | 1387 | 1124.5 | 1155 | 1255.5 | 1369 | 1062.5 | 1358.5 | 1591 | 1084 |
| **1017.5** | 1222 | 861.5 | 1010.5 | 1275.5 | 1103.5 | 1334.5 | 910.5 | 1361.5 | 1431 |
| **1182.5** | 1506.5 | 1077.5 | 811.5 | 1457.5 | 1263 | 1221 | 734.5 | 1054.5 | 840.5 |
| **1017** | 1284.5 | 1308 | 1212.5 | 1116.5 | 1290 | 1280.5 | 963.5 | 1168.5 | 1374.5 |
| **1013** | 1102 | 1339 | 1230.5 | 941.5 | 1282.5 | 1320.5 | 1323 | 1334.5 | 1153 |
| **1486** | 1297.5 | 915 | 1564.5 | 778 | 881.5 | 1148.5 | 1232.5 | 1129.5 | 1162.5 |
| **1255.5** | 1746.5 | 1056 | 869.5 | 1048.5 | 1020 | 1091.5 | 1253 | 1457.5 | 1173.5 |
| **894** | 1217 | 1194.5 | 831 | 946 | 890 | 624.5 | 1315.5 | 907 | 907 |
| **1234** | 967.5 | 1196 | 1654.5 | 1144.5 | 1389.5 | 1449 | 974.5 | 1133 | 1672 |
| **1016.5** | 1216.5 | 1125.5 | 1529 | 959.5 | 1019.5 | 1597 | 701 | 1184.5 | 1293.5 |
| **1175** | 910.5 | 930.5 | 1121 | 1150 | 1158 | 866 | 971.5 | 708.5 | 1240 |
| **1123.5** | 908 | 945.5 | 1469 | 1204 | 1347 | 982 | 1712 | 1154.5 | 855 |
| **1504** | 749 | 1495.5 | 1434 | 1367.5 | 1194.5 | 1203 | 1545.5 | 1171 | 1147.5 |
| **811** | 1214.5 | 1254.5 | 1196 | 1445.5 | 1022.5 | 1183 | 1580.5 | 1362 | 1262 |
| **1674.5** | 1244.5 | 1557.5 | 1382.5 | 1071 | 1132.5 | 1193 | 1244.5 | 1086.5 | 1405 |
| **1588** | 1222 | 1781 | 808.5 | 637 | 1224.5 | 1304 | 853.5 | 1160.5 | 1139.5 |
| **1244.5** | 1603.5 | 1171.5 | 1543.5 | 1356.5 | 1399.5 | 1527.5 | 1287.5 | 1343 | 1533 |
| **1605** | 1419.5 | 960.5 | 1221 | 1233.5 | 1167.5 | 1118 | 1191.5 | 1697.5 | 1637 |
| **1030.5** | 704.5 | 1003.5 | 1456 | 1011 | 807.5 | 1738 | 1220.5 | 952 | 1308.5 |
| **988** | 1415 | 956 | 1358.5 | 1027.5 | 1732 | 1240 | 1115 | 822.5 | 1651.5 |
| **775.5** | 1660.5 | 980.5 | 843 | 1285.5 | 1182 | 1045 | 1415 | 1004 | 1573 |
| **923.5** | 1090.5 | 1466 | 1005.5 | 1691 | 1403.5 | 1106 | 1159.5 | 1502 | 935 |
| **1275** | 954.5 | 1165.5 | 762.5 | 1267 | 1477.5 | 1263 | 1275 | 834.5 | 1199.5 |
| **1250.5** | 839.5 | 1133 | 1156.5 | 951.5 | 1266 | 984.5 | 1472.5 | 1502.5 | 871.5 |
| **822.5** | 1101.5 | 907 | 731 | 1232.5 | 957.5 | 1456 | 1302.5 | 890.5 | 1324.5 |
| **1280.5** | 1269.5 | 1289.5 | 725 | 906 | 987 | 1011 | 1353 | 1293 | 1248 |

| 1605.5 | 1210.5 | 1178.5 | 1328 | 1234.5 | 951 | 1090.5 | 1687.5 | 1388 | 1504 |
|---|---|---|---|---|---|---|---|---|---|
| **940** | 989.5 | 1220.5 | 1192 | 1089.5 | 1675 | 1667 | 1537.5 | 1376 | 1309 |
| **1253** | 1256 | 1050 | 1326 | 1002.5 | 1522 | 1005.5 | 1149 | 1099.5 | 1067 |
| **1576** | 1271 | 1569.5 | 750.5 | 859.5 | 1489.5 | 1138.5 | 1225.5 | 1222.5 | 870 |
| **1561.5** | 1153 | 671.5 | 1184 | 1470 | 1196.5 | 1253.5 | 1341.5 | 1136 | 1120 |
| **987.5** | 1592.5 | 1421.5 | 1219 | 714.5 | 1396 | 804.5 | 1016.5 | 1702.5 | 1224.5 |
| **715.5** | 1385.5 | 1300.5 | 1748 | 964.5 | 1339 | 1004 | 747.5 | 1535.5 | 831.5 |
| **1269** | 1326 | 1633.5 | 1565 | 993.5 | 730 | 1198.5 | 1405 | 1636 | 1296 |
| **968** | 1021 | 769.5 | 1464 | 734.5 | 996 | 1019 | 1094.5 | 997.5 | 1436.5 |
| **1118** | 983 | 1083 | 1151.5 | 1502 | 939 | 1179.5 | 1120 | 976.5 | 816 |
| **727** | 962.5 | 1187.5 | 991.5 | 992.5 | 1060 | 872 | 1193 | 1271.5 | 1001.5 |
| **1271** | 1757 | 943 | 1243 | 1557 | 1429.5 | 1092 | 1154 | 1342 | 1250.5 |
| **1253** | 662.5 | 751.5 | 1314 | 1159.5 | 1722 | 1441 | 1713.5 | 1197.5 | 1223 |
| **1434.5** | 1265 | 740.5 | 1189 | 1439 | 1470 | 1576 | 1052.5 | 989.5 | 1331 |
| **941.5** | 1571 | 1152 | 1323.5 | 1125.5 | 1718.5 | 1723 | 1638.5 | 1468.5 | 1492 |
| **1430.5** | 1553.5 | 1306.5 | 1170.5 | 1435 | 1081 | 1314.5 | 1016.5 | 1137.5 | 1324.5 |
| **1321.5** | 1027.5 | 1006 | 860 | 942.5 | 1312 | 1694 | 1402 | 1232 | 1177 |
| **1123** | 1579.5 | 1325 | 929.5 | 941 | 1279.5 | 1128 | 1122.5 | 1158 | 1107.5 |
| **1415.5** | 1377 | 1514 | 1755 | 932.5 | 827 | 1519 | 1349.5 | 1216 | 802.5 |
| **996** | 1383.5 | 1435.5 | 1331.5 | 1107 | 1348 | 1387.5 | 949.5 | 776.5 | 963.5 |

Death tolls user defined

| **74** | **74** | **74** | **74** | **74** | **74** | **74** | **74** | **74** | **74** |
|---|---|---|---|---|---|---|---|---|---|
| **74** | 74 | 74 | 74 | 74 | 74 | 74 | 74 | 74 | 74 |
| **74** | 74 | 74 | 74 | 74 | 74 | 74 | 74 | 74 | 74 |
| **74** | 74 | 74 | 74 | 74 | 74 | 74 | 74 | 74 | 74 |
| **74** | 74 | 74 | 74 | 74 | 74 | 74 | 74 | 74 | 74 |
| **74** | 74 | 74 | 74 | 74 | 74 | 74 | 74 | 74 | 74 |
| **74** | 74 | 74 | 74 | 74 | 74 | 74 | 74 | 74 | 74 |
| **74** | 74 | 74 | 74 | 74 | 74 | 74 | 74 | 74 | 74 |
| **74** | 74 | 74 | 74 | 74 | 74 | 74 | 74 | 74 | 74 |
| **74** | 74 | 74 | 74 | 74 | 74 | 74 | 74 | 74 | 74 |
| **74** | 74 | 74 | 74 | 74 | 74 | 74 | 74 | 74 | 74 |
| **74** | 74 | 74 | 74 | 74 | 74 | 74 | 74 | 74 | 74 |
| **74** | 74 | 74 | 74 | 74 | 74 | 74 | 74 | 74 | 74 |
| **74** | 74 | 74 | 74 | 74 | 74 | 74 | 74 | 74 | 74 |
| **74** | 74 | 74 | 74 | 74 | 74 | 74 | 74 | 74 | 74 |
| **74** | 74 | 74 | 74 | 74 | 74 | 74 | 74 | 74 | 74 |
| **74** | 74 | 74 | 74 | 74 | 74 | 74 | 74 | 74 | 74 |
| **74** | 74 | 74 | 74 | 74 | 74 | 74 | 74 | 74 | 74 |
| **74** | 74 | 74 | 74 | 74 | 74 | 74 | 74 | 74 | 74 |
| **74** | 74 | 74 | 74 | 74 | 74 | 74 | 74 | 74 | 74 |
| **74** | 74 | 74 | 74 | 74 | 74 | 74 | 74 | 74 | 74 |
| **74** | 74 | 74 | 74 | 74 | 74 | 74 | 74 | 74 | 74 |
| **74** | 74 | 74 | 74 | 74 | 74 | 74 | 74 | 74 | 74 |
| **74** | 74 | 74 | 74 | 74 | 74 | 74 | 74 | 74 | 74 |
| **74** | 74 | 74 | 74 | 74 | 74 | 74 | 74 | 74 | 74 |
| **74** | 74 | 74 | 74 | 74 | 74 | 74 | 74 | 74 | 74 |
| **74** | 74 | 74 | 74 | 74 | 74 | 74 | 74 | 74 | 74 |
| **74** | 74 | 74 | 74 | 74 | 74 | 74 | 74 | 74 | 74 |
| **74** | 74 | 74 | 74 | 74 | 74 | 74 | 74 | 74 | 74 |
| **74** | 74 | 74 | 74 | 74 | 74 | 74 | 74 | 74 | 74 |
| **74** | 74 | 74 | 74 | 74 | 74 | 74 | 74 | 74 | 74 |
| **74** | 74 | 74 | 74 | 74 | 74 | 74 | 74 | 74 | 74 |
| **74** | 74 | 74 | 74 | 74 | 74 | 74 | 74 | 74 | 74 |

| **74** | **74** | **74** | **74** | **74** | **74** | **74** | **74** | **74** | **74** |
|---|---|---|---|---|---|---|---|---|---|
| **74** | 74 | 74 | 74 | 74 | 74 | 74 | 74 | 74 | 74 |
| **74** | 74 | 74 | 74 | 74 | 74 | 74 | 74 | 74 | 74 |
| **74** | 74 | 74 | 74 | 74 | 74 | 74 | 74 | 74 | 74 |
| **74** | 74 | 74 | 74 | 74 | 74 | 74 | 74 | 74 | 74 |
| **74** | 74 | 74 | 74 | 74 | 74 | 74 | 74 | 74 | 74 |
| **74** | 74 | 74 | 74 | 74 | 74 | 74 | 74 | 74 | 74 |
| **74** | 74 | 74 | 74 | 74 | 74 | 74 | 74 | 74 | 74 |
| **74** | 74 | 74 | 74 | 74 | 74 | 74 | 74 | 74 | 74 |
| **74** | 74 | 74 | 74 | 74 | 74 | 74 | 74 | 74 | 74 |
| **74** | 74 | 74 | 74 | 74 | 74 | 74 | 74 | 74 | 74 |
| **74** | 74 | 74 | 74 | 74 | 74 | 74 | 74 | 74 | 74 |
| **74** | 74 | 74 | 74 | 74 | 74 | 74 | 74 | 74 | 74 |
| **74** | 74 | 74 | 74 | 74 | 74 | 74 | 74 | 74 | 74 |
| **74** | 74 | 74 | 74 | 74 | 74 | 74 | 74 | 74 | 74 |
| **74** | 74 | 74 | 74 | 74 | 74 | 74 | 74 | 74 | 74 |
| **74** | 74 | 74 | 74 | 74 | 74 | 74 | 74 | 74 | 74 |
| **74** | 74 | 74 | 74 | 74 | 74 | 74 | 74 | 74 | 74 |
| **74** | 74 | 74 | 74 | 74 | 74 | 74 | 74 | 74 | 74 |
| **74** | 74 | 74 | 74 | 74 | 74 | 74 | 74 | 74 | 74 |
| **74** | 74 | 74 | 74 | 74 | 74 | 74 | 74 | 74 | 74 |
| **74** | 74 | 74 | 74 | 74 | 74 | 74 | 74 | 74 | 74 |
| **74** | 74 | 74 | 74 | 74 | 74 | 74 | 74 | 74 | 74 |
| **74** | 74 | 74 | 74 | 74 | 74 | 74 | 74 | 74 | 74 |
| **74** | 74 | 74 | 74 | 74 | 74 | 74 | 74 | 74 | 74 |
| **74** | 74 | 74 | 74 | 74 | 74 | 74 | 74 | 74 | 74 |
| **74** | 74 | 74 | 74 | 74 | 74 | 74 | 74 | 74 | 74 |
| **74** | 74 | 74 | 74 | 74 | 74 | 74 | 74 | 74 | 74 |
| **74** | 74 | 74 | 74 | 74 | 74 | 74 | 74 | 74 | 74 |
| **74** | 74 | 74 | 74 | 74 | 74 | 74 | 74 | 74 | 74 |
| **74** | 74 | 74 | 74 | 74 | 74 | 74 | 74 | 74 | 74 |
| **74** | 74 | 74 | 74 | 74 | 74 | 74 | 74 | 74 | 74 |
| **74** | 74 | 74 | 74 | 74 | 74 | 74 | 74 | 74 | 74 |
| **74** | 74 | 74 | 74 | 74 | 74 | 74 | 74 | 74 | 74 |

| 74 | 74 | 74 | 74 | 74 | 74 | 74 | 74 | 74 | 74 |
|---|---|---|---|---|---|---|---|---|---|
| **74** | 74 | 74 | 74 | 74 | 74 | 74 | 74 | 74 | 74 |
| **74** | 74 | 74 | 74 | 74 | 74 | 74 | 74 | 74 | 74 |
| **74** | 74 | 74 | 74 | 74 | 74 | 74 | 74 | 74 | 74 |
| **74** | 74 | 74 | 74 | 74 | 74 | 74 | 74 | 74 | 74 |
| **74** | 74 | 74 | 74 | 74 | 74 | 74 | 74 | 74 | 74 |
| **74** | 74 | 74 | 74 | 74 | 74 | 74 | 74 | 74 | 74 |
| **74** | 74 | 74 | 74 | 74 | 74 | 74 | 74 | 74 | 74 |
| **74** | 74 | 74 | 74 | 74 | 74 | 74 | 74 | 74 | 74 |
| **74** | 74 | 74 | 74 | 74 | 74 | 74 | 74 | 74 | 74 |
| **74** | 74 | 74 | 74 | 74 | 74 | 74 | 74 | 74 | 74 |
| **74** | 74 | 74 | 74 | 74 | 74 | 74 | 74 | 74 | 74 |
| **74** | 74 | 74 | 74 | 74 | 74 | 74 | 74 | 74 | 74 |
| **74** | 74 | 74 | 74 | 74 | 74 | 74 | 74 | 74 | 74 |
| **74** | 74 | 74 | 74 | 74 | 74 | 74 | 74 | 74 | 74 |
| **74** | 74 | 74 | 74 | 74 | 74 | 74 | 74 | 74 | 74 |
| **74** | 74 | 74 | 74 | 74 | 74 | 74 | 74 | 74 | 74 |
| **74** | 74 | 74 | 74 | 74 | 74 | 74 | 74 | 74 | 74 |
| **74** | 74 | 74 | 74 | 74 | 74 | 74 | 74 | 74 | 74 |
| **74** | 74 | 74 | 74 | 74 | 74 | 74 | 74 | 74 | 74 |
| **74** | 74 | 74 | 74 | 74 | 74 | 74 | 74 | 74 | 74 |
| **74** | 74 | 74 | 74 | 74 | 74 | 74 | 74 | 74 | 74 |
| **74** | 74 | 74 | 74 | 74 | 74 | 74 | 74 | 74 | 74 |
| **74** | 74 | 74 | 74 | 74 | 74 | 74 | 74 | 74 | 74 |
| **74** | 74 | 74 | 74 | 74 | 74 | 74 | 74 | 74 | 74 |
| **74** | 74 | 74 | 74 | 74 | 74 | 74 | 74 | 74 | 74 |
| **74** | 74 | 74 | 74 | 74 | 74 | 74 | 74 | 74 | 74 |
| **74** | 74 | 74 | 74 | 74 | 74 | 74 | 74 | 74 | 74 |
| **74** | 74 | 74 | 74 | 74 | 74 | 74 | 74 | 74 | 74 |
| **74** | 74 | 74 | 74 | 74 | 74 | 74 | 74 | 74 | 74 |
| **74** | 74 | 74 | 74 | 74 | 74 | 74 | 74 | 74 | 74 |
| **74** | 74 | 74 | 74 | 74 | 74 | 74 | 74 | 74 | 74 |
| **74** | 74 | 74 | 74 | 74 | 74 | 74 | 74 | 74 | 74 |
| **74** | 74 | 74 | 74 | 74 | 74 | 74 | 74 | 74 | 74 |

| 74 | 74 | 74 | 74 | 74 | 74 | 74 | 74 | 74 | 74 |
|---|---|---|---|---|---|---|---|---|---|
| **74** | 74 | 74 | 74 | 74 | 74 | 74 | 74 | 74 | 74 |
| **74** | 74 | 74 | 74 | 74 | 74 | 74 | 74 | 74 | 74 |
| **74** | 74 | 74 | 74 | 74 | 74 | 74 | 74 | 74 | 74 |
| **74** | 74 | 74 | 74 | 74 | 74 | 74 | 74 | 74 | 74 |
| **74** | 74 | 74 | 74 | 74 | 74 | 74 | 74 | 74 | 74 |
| **74** | 74 | 74 | 74 | 74 | 74 | 74 | 74 | 74 | 74 |
| **74** | 74 | 74 | 74 | 74 | 74 | 74 | 74 | 74 | 74 |
| **74** | 74 | 74 | 74 | 74 | 74 | 74 | 74 | 74 | 74 |
| **74** | 74 | 74 | 74 | 74 | 74 | 74 | 74 | 74 | 74 |
| **74** | 74 | 74 | 74 | 74 | 74 | 74 | 74 | 74 | 74 |
| **74** | 74 | 74 | 74 | 74 | 74 | 74 | 74 | 74 | 74 |
| **74** | 74 | 74 | 74 | 74 | 74 | 74 | 74 | 74 | 74 |
| **74** | 74 | 74 | 74 | 74 | 74 | 74 | 74 | 74 | 74 |
| **74** | 74 | 74 | 74 | 74 | 74 | 74 | 74 | 74 | 74 |
| **74** | 74 | 74 | 74 | 74 | 74 | 74 | 74 | 74 | 74 |
| **74** | 74 | 74 | 74 | 74 | 74 | 74 | 74 | 74 | 74 |
| **74** | 74 | 74 | 74 | 74 | 74 | 74 | 74 | 74 | 74 |
| **74** | 74 | 74 | 74 | 74 | 74 | 74 | 74 | 74 | 74 |
| **74** | 74 | 74 | 74 | 74 | 74 | 74 | 74 | 74 | 74 |
| **74** | 74 | 74 | 74 | 74 | 74 | 74 | 74 | 74 | 74 |
| **74** | 74 | 74 | 74 | 74 | 74 | 74 | 74 | 74 | 74 |

Death tolls mean 300 normal distribution

| **566** | **370** | **452** | **366** | **283** | **259** | **257** | **321** | **124** | **308** |
|---|---|---|---|---|---|---|---|---|---|
| **252** | 376 | 169 | 395 | 387 | 434 | 309 | 307 | 324 | 291 |
| **335** | 172 | 392 | 214 | 545 | 201 | 334 | 456 | 286 | 18 |
| **359** | 542 | 296 | 211 | 110 | 187 | 183 | 482 | 346 | 154 |
| **362** | 517 | 332 | 370 | 399 | 245 | 346 | 584 | 358 | 144 |
| **287** | 492 | 138 | 194 | 484 | 356 | 112 | 145 | 369 | 304 |
| **284** | 193 | 263 | 578 | 302 | 149 | 117 | 223 | 372 | 445 |
| **446** | 298 | 128 | 565 | 372 | 418 | 517 | 37 | 339 | 214 |
| **511** | 137 | 395 | 194 | 519 | 327 | 319 | 333 | 247 | 362 |
| **485** | 320 | 226 | 86 | 381 | 389 | 338 | 409 | 249 | 407 |
| **466** | 249 | 173 | 497 | 108 | 187 | 427 | 242 | 329 | 87 |
| **349** | 366 | 257 | 306 | 333 | 480 | 229 | 279 | 265 | 284 |
| **307** | 194 | 115 | 265 | 332 | 174 | 332 | 312 | 340 | 460 |
| **477** | 276 | 590 | 370 | 403 | 134 | 263 | 505 | 295 | 523 |
| **192** | 345 | 409 | 206 | 565 | 347 | 398 | 242 | 118 | 507 |
| **417** | 258 | 338 | 285 | 326 | 222 | 543 | 458 | 328 | 272 |
| **147** | 223 | 453 | 424 | 459 | 213 | 387 | 209 | 430 | 243 |
| **485** | 429 | 321 | 266 | 435 | 463 | 230 | 378 | 376 | 466 |
| **308** | 260 | 242 | 46 | 163 | 276 | 323 | 462 | 370 | 378 |
| **441** | 318 | 405 | 245 | 294 | 189 | 177 | 203 | 320 | 314 |
| **301** | 141 | 421 | 189 | 294 | 222 | 505 | 487 | 346 | 424 |
| **328** | 344 | 494 | 508 | 422 | 316 | 318 | 212 | 538 | 41 |
| **442** | 254 | 173 | 311 | 403 | 256 | 568 | 363 | 425 | 215 |
| **387** | 65 | 518 | 257 | 158 | 195 | 547 | 390 | 324 | 125 |
| **197** | 214 | 447 | 176 | 454 | 261 | 391 | 396 | 110 | 198 |
| **273** | 230 | 304 | 137 | 294 | 318 | 444 | 281 | 420 | 271 |
| **328** | 483 | 319 | 112 | 255 | 471 | 318 | 418 | 207 | 367 |
| **433** | 307 | 449 | 146 | 134 | 171 | 204 | 453 | 311 | 264 |
| **53** | 332 | 141 | 98 | 460 | 400 | 87 | 489 | 377 | 210 |
| **51** | 302 | 396 | 279 | 349 | 244 | 486 | 386 | 519 | 479 |
| **278** | 485 | 241 | 456 | 473 | 392 | 297 | 526 | 369 | 393 |
| **360** | 304 | 270 | 307 | 296 | 317 | 357 | 254 | 325 | 181 |
| **526** | 157 | 335 | 361 | 391 | 525 | 89 | 337 | 448 | 253 |

| 566 | 370 | 452 | 366 | 283 | 259 | 257 | 321 | 124 | 308 |
|---|---|---|---|---|---|---|---|---|---|
| **451** | 503 | 275 | 193 | 318 | 537 | 450 | 496 | 46 | 383 |
| **421** | 317 | 474 | 369 | 460 | 320 | 454 | 290 | 267 | 249 |
| **475** | 309 | 284 | 459 | 313 | 253 | 418 | 486 | 199 | 257 |
| **44** | 325 | 101 | 368 | 374 | 193 | 421 | 133 | 494 | 146 |
| **239** | 445 | 154 | 376 | 536 | 506 | 316 | 306 | 297 | 312 |
| **123** | 189 | 342 | 524 | 187 | 384 | 238 | 435 | 76 | 433 |
| **296** | 291 | 276 | 246 | 407 | 234 | 163 | 176 | 301 | 377 |
| **271** | 280 | 530 | 319 | 490 | 100 | 256 | 41 | 192 | 412 |
| **323** | 518 | 477 | 397 | 226 | 354 | 205 | 385 | 389 | 396 |
| **300** | 259 | 33 | 274 | 71 | 367 | 349 | 229 | 319 | 211 |
| **454** | 529 | 251 | 230 | 239 | 238 | 390 | 288 | 257 | 457 |
| **381** | 302 | 538 | 306 | 210 | 363 | 303 | 213 | 403 | 316 |
| **215** | 460 | 212 | 224 | 313 | 289 | 190 | 465 | 187 | 231 |
| **391** | 287 | 255 | 245 | 360 | 315 | 528 | 281 | 420 | 273 |
| **395** | 139 | 197 | 264 | 74 | 334 | 198 | 300 | 474 | 383 |
| **283** | 160 | 495 | 228 | 389 | 490 | 355 | 336 | 348 | 337 |
| **246** | 329 | 467 | 337 | 152 | 389 | 79 | 228 | 260 | 291 |
| **150** | 260 | 435 | 51 | 305 | 246 | 403 | 431 | 481 | 338 |
| **264** | 206 | 178 | 117 | 212 | 428 | 294 | 525 | 323 | 312 |
| **121** | 370 | 428 | 198 | 150 | 230 | 277 | 357 | 71 | 357 |
| **217** | 179 | 325 | 333 | 444 | 529 | 148 | 275 | 486 | 281 |
| **336** | 487 | 305 | 322 | 446 | 519 | 126 | 363 | 555 | 309 |
| **592** | 180 | 498 | 517 | 390 | 280 | 60 | 414 | 192 | 188 |
| **491** | 247 | 475 | 475 | 231 | 451 | 304 | 287 | 541 | 163 |
| **371** | 338 | 441 | 173 | 332 | 388 | 481 | 321 | 134 | 278 |
| **424** | 252 | 134 | 312 | 347 | 303 | 482 | 212 | 550 | 293 |
| **362** | 127 | 30 | 526 | 331 | 315 | 480 | 320 | 55 | 286 |
| **258** | 395 | 242 | 222 | 11 | 118 | 276 | 126 | 363 | 61 |
| **253** | 405 | 323 | 110 | 455 | 491 | 357 | 314 | 424 | 357 |
| **300** | 210 | 174 | 428 | 115 | 392 | 428 | 285 | 222 | 220 |
| **524** | 300 | 123 | 476 | 74 | 439 | 221 | 339 | 232 | 342 |
| **209** | 473 | 285 | 378 | 408 | 110 | 281 | 375 | 311 | 338 |
| **445** | 274 | 332 | 253 | 267 | 78 | 381 | 303 | 211 | 561 |

| 566 | 370 | 452 | 366 | 283 | 259 | 257 | 321 | 124 | 308 |
|---|---|---|---|---|---|---|---|---|---|
| 424 | 417 | 282 | 523 | 331 | 380 | 293 | 43 | 195 | 413 |
| 464 | 281 | 27 | 330 | 334 | 290 | 159 | 405 | 189 | 454 |
| 336 | 364 | 430 | 225 | 159 | 319 | 249 | 490 | 169 | 146 |
| 426 | 320 | 134 | 238 | 438 | 49 | 425 | 270 | 169 | 243 |
| 311 | 32 | 369 | 60 | 530 | 188 | 228 | 212 | 356 | 277 |
| 397 | 218 | 491 | 274 | 280 | 553 | 86 | 246 | 453 | 71 |
| 240 | 444 | 506 | 223 | 483 | 306 | 258 | 349 | 101 | 355 |
| 398 | 272 | 400 | 257 | 457 | 405 | 127 | 226 | 193 | 114 |
| 415 | 273 | 314 | 341 | 258 | 140 | 217 | 359 | 271 | 419 |
| 401 | 392 | 465 | 202 | 408 | 347 | 370 | 205 | 368 | 379 |
| 374 | 121 | 421 | 335 | 361 | 446 | 266 | 420 | 151 | 525 |
| 86 | 311 | 310 | 288 | 186 | 92 | 335 | 224 | 333 | 342 |
| 429 | 437 | 302 | 412 | 460 | 372 | 253 | 253 | 471 | 374 |
| 206 | 476 | 298 | 347 | 302 | 290 | 288 | 405 | 457 | 24 |
| 447 | 39 | 209 | 351 | 417 | 310 | 312 | 418 | 514 | 283 |
| 370 | 176 | 392 | 325 | 109 | 420 | 360 | 249 | 74 | 429 |
| 337 | 257 | 429 | 477 | 275 | 171 | 288 | 303 | 335 | 462 |
| 390 | 294 | 137 | 332 | 254 | 284 | 265 | 276 | 279 | 337 |
| 557 | 384 | 424 | 299 | 445 | 46 | 136 | 161 | 287 | 181 |
| 340 | 465 | 381 | 440 | 429 | 280 | 468 | 237 | 206 | 136 |
| 424 | 224 | 251 | 503 | 414 | 338 | 161 | 153 | 232 | 324 |
| 154 | 375 | 285 | 248 | 510 | 361 | 518 | 297 | 339 | 124 |
| 296 | 498 | 179 | 277 | 351 | 229 | 578 | 95 | 125 | 132 |
| 244 | 190 | 41 | 84 | 519 | 357 | 333 | 379 | 241 | 395 |
| 502 | 479 | 282 | 256 | 63 | 132 | 260 | 412 | 113 | 193 |
| 190 | 402 | 150 | 477 | 144 | 401 | 305 | 515 | 318 | 224 |
| 457 | 182 | 295 | 415 | 109 | 334 | 408 | 169 | 243 | 347 |
| 546 | 294 | 49 | 296 | 512 | 181 | 361 | 321 | 258 | 251 |
| 453 | 232 | 273 | 493 | 12 | 292 | 175 | 219 | 293 | 231 |
| 185 | 206 | 316 | 257 | 362 | 253 | 184 | 230 | 398 | 480 |
| 470 | 317 | 526 | 291 | 404 | 269 | 546 | 371 | 511 | 239 |
| 265 | 474 | 333 | 129 | 513 | 384 | 523 | 45 | 157 | 495 |
| 512 | 273 | 310 | 301 | 275 | 311 | 266 | 221 | 139 | 415 |

| **566** | **370** | **452** | **366** | **283** | **259** | **257** | **321** | **124** | **308** |
|---|---|---|---|---|---|---|---|---|---|
| **401** | 160 | 150 | 426 | 399 | 248 | 470 | 263 | 352 | 559 |
| **333** | 222 | 130 | 184 | 576 | 352 | 291 | 375 | 508 | 220 |
| **271** | 303 | 422 | 499 | 379 | 263 | 308 | 385 | 183 | 408 |
| **314** | 367 | 185 | 201 | 221 | 403 | 257 | 295 | 433 | 201 |
| **330** | 197 | 181 | 113 | 240 | 350 | 291 | 157 | 555 | 140 |
| **97** | 172 | 251 | 274 | 446 | 534 | 396 | 216 | 281 | 315 |
| **452** | 336 | 309 | 258 | 172 | 378 | 350 | 206 | 60 | 99 |
| **203** | 146 | 385 | 288 | 237 | 322 | 78 | 291 | 302 | 341 |
| **155** | 446 | 230 | 227 | 325 | 465 | 215 | 438 | 60 | 180 |
| **511** | 131 | 383 | 79 | 323 | 331 | 431 | 144 | 386 | 551 |
| **133** | 473 | 333 | 346 | 160 | 469 | 257 | 519 | 327 | 317 |
| **221** | 374 | 388 | 302 | 315 | 165 | 266 | 252 | 358 | 211 |
| **258** | 484 | 474 | 185 | 182 | 257 | 356 | 397 | 265 | 307 |
| **239** | 282 | 212 | 187 | 132 | 47 | 301 | 218 | 385 | 370 |
| **282** | 314 | 205 | 322 | 282 | 128 | 572 | 115 | 299 | 314 |
| **466** | 304 | 217 | 432 | 301 | 398 | 137 | 578 | 131 | 355 |
| **84** | 259 | 304 | 332 | 508 | 63 | 391 | 508 | 219 | 295 |
| **390** | 326 | 318 | 361 | 445 | 51 | 288 | 377 | 466 | 509 |
| **290** | 278 | 247 | 428 | 423 | 446 | 221 | 464 | 257 | 171 |
| **282** | 449 | 313 | 246 | 371 | 314 | 103 | 338 | 239 | 338 |
| **464** | 126 | 556 | 321 | 275 | 245 | 347 | 66 | 339 | 456 |

Results (user defined death toll and paste in N2021)

| 320 | 331 | 295 | 356 | 335 | 264 | 284 | 306 | 383 | 289 |
|---|---|---|---|---|---|---|---|---|---|
| **277** | 309 | 312 | 331 | 283 | 341 | 328 | 431 | 319 | 267 |
| **268** | 306 | 318 | 261 | 375 | 354 | 383 | 328 | 346 | 367 |
| **415** | 329 | 337 | 280 | 264 | 343 | 296 | 387 | 416 | 317 |
| **363** | 338 | 280 | 337 | 263 | 297 | 289 | 297 | 248 | 304 |
| **361** | 274 | 335 | 287 | 334 | 298 | 287 | 271 | 312 | 324 |
| **340** | 338 | 321 | 337 | 291 | 393 | 317 | 316 | 421 | 319 |
| **352** | 318 | 311 | 341 | 274 | 407 | 388 | 380 | 393 | 310 |
| **417** | 303 | 334 | 406 | 340 | 312 | 387 | 333 | 347 | 393 |
| **324** | 289 | 369 | 323 | 293 | 406 | 372 | 272 | 351 | 399 |
| **304** | 298 | 289 | 322 | 284 | 374 | 270 | 366 | 255 | 253 |
| **330** | 286 | 340 | 300 | 396 | 378 | 336 | 266 | 361 | 419 |
| **302** | 326 | 369 | 347 | 382 | 364 | 374 | 424 | 360 | 334 |
| **255** | 298 | 346 | 363 | 381 | 314 | 274 | 334 | 343 | 405 |
| **300** | 382 | 290 | 365 | 360 | 315 | 326 | 401 | 383 | 366 |
| **374** | 313 | 359 | 397 | 272 | 391 | 398 | 348 | 278 | 294 |
| **319** | 277 | 347 | 378 | 273 | 273 | 354 | 358 | 293 | 278 |
| **389** | 295 | 343 | 274 | 325 | 287 | 283 | 323 | 325 | 336 |
| **275** | 263 | 361 | 304 | 326 | 266 | 260 | 347 | 270 | 343 |
| **264** | 380 | 390 | 378 | 365 | 311 | 329 | 320 | 314 | 275 |
| **340** | 336 | 296 | 361 | 388 | 357 | 318 | 380 | 264 | 396 |
| **327** | 310 | 328 | 279 | 403 | 267 | 369 | 304 | 275 | 291 |
| **272** | 271 | 396 | 354 | 290 | 385 | 332 | 348 | 360 | 340 |
| **285** | 370 | 307 | 265 | 387 | 281 | 392 | 351 | 367 | 294 |
| **281** | 351 | 390 | 310 | 313 | 350 | 284 | 276 | 367 | 312 |
| **311** | 263 | 317 | 351 | 263 | 384 | 390 | 329 | 355 | 274 |
| **306** | 328 | 311 | 373 | 349 | 252 | 383 | 368 | 366 | 299 |
| **334** | 372 | 328 | 328 | 266 | 362 | 292 | 359 | 348 | 290 |
| **337** | 373 | 311 | 306 | 279 | 259 | 322 | 323 | 338 | 309 |
| **261** | 396 | 313 | 313 | 324 | 255 | 328 | 368 | 350 | 397 |
| **370** | 356 | 291 | 258 | 379 | 303 | 327 | 301 | 361 | 282 |
| **310** | 360 | 378 | 340 | 399 | 366 | 317 | 355 | 258 | 309 |
| **388** | 313 | 320 | 357 | 287 | 359 | 273 | 347 | 424 | 305 |

| 320 | 331 | 295 | 356 | 335 | 264 | 284 | 306 | 383 | 289 |
|---|---|---|---|---|---|---|---|---|---|
| **366** | 307 | 358 | 349 | 261 | 315 | 376 | 331 | 275 | 365 |
| **368** | 372 | 306 | 289 | 330 | 257 | 281 | 338 | 263 | 397 |
| **292** | 351 | 329 | 362 | 289 | 297 | 284 | 299 | 357 | 288 |
| **362** | 339 | 364 | 296 | 336 | 426 | 318 | 383 | 331 | 288 |
| **376** | 423 | 382 | 298 | 357 | 379 | 313 | 274 | 388 | 333 |
| **345** | 293 | 348 | 344 | 364 | 273 | 278 | 258 | 354 | 338 |
| **266** | 398 | 310 | 354 | 237 | 299 | 361 | 313 | 362 | 298 |
| **368** | 338 | 341 | 254 | 380 | 262 | 367 | 297 | 353 | 292 |
| **382** | 409 | 327 | 374 | 423 | 305 | 378 | 271 | 347 | 407 |
| **311** | 335 | 313 | 367 | 272 | 293 | 268 | 290 | 331 | 292 |
| **273** | 383 | 270 | 348 | 315 | 322 | 345 | 287 | 253 | 330 |
| **321** | 318 | 286 | 330 | 277 | 271 | 347 | 351 | 310 | 388 |
| **333** | 394 | 365 | 328 | 320 | 269 | 343 | 338 | 261 | 317 |
| **349** | 267 | 296 | 297 | 243 | 273 | 292 | 328 | 263 | 274 |
| **361** | 402 | 256 | 330 | 291 | 267 | 243 | 396 | 368 | 362 |
| **345** | 376 | 315 | 317 | 289 | 269 | 274 | 314 | 387 | 406 |
| **287** | 276 | 291 | 341 | 309 | 312 | 335 | 317 | 330 | 317 |
| **352** | 314 | 313 | 339 | 241 | 369 | 253 | 384 | 357 | 423 |
| **378** | 456 | 300 | 261 | 342 | 311 | 357 | 300 | 360 | 306 |
| **288** | 336 | 398 | 365 | 391 | 321 | 344 | 359 | 341 | 343 |
| **347** | 356 | 413 | 275 | 298 | 373 | 316 | 362 | 359 | 321 |
| **409** | 311 | 357 | 347 | 279 | 398 | 326 | 362 | 304 | 395 |
| **351** | 321 | 386 | 342 | 265 | 348 | 308 | 309 | 385 | 411 |
| **294** | 358 | 365 | 336 | 350 | 350 | 309 | 305 | 319 | 332 |
| **368** | 342 | 277 | 390 | 348 | 275 | 274 | 375 | 388 | 302 |
| **395** | 332 | 299 | 356 | 310 | 428 | 288 | 325 | 312 | 293 |
| **370** | 317 | 306 | 278 | 364 | 343 | 351 | 381 | 318 | 313 |
| **312** | 367 | 383 | 347 | 380 | 322 | 345 | 255 | 292 | 369 |
| **267** | 369 | 370 | 286 | 290 | 371 | 329 | 340 | 325 | 260 |
| **338** | 403 | 387 | 309 | 295 | 247 | 272 | 307 | 294 | 267 |
| **306** | 286 | 267 | 287 | 316 | 274 | 279 | 357 | 273 | 277 |
| **343** | 293 | 284 | 332 | 386 | 279 | 264 | 309 | 285 | 316 |
| **239** | 360 | 421 | 269 | 338 | 291 | 329 | 438 | 295 | 367 |

| 320 | 331 | 295 | 356 | 335 | 264 | 284 | 306 | 383 | 289 |
|---|---|---|---|---|---|---|---|---|---|
| **298** | 393 | 331 | 392 | 339 | 397 | 284 | 370 | 335 | 316 |
| **275** | 340 | 333 | 370 | 323 | 312 | 384 | 322 | 360 | 264 |
| **277** | 332 | 303 | 287 | 284 | 404 | 331 | 276 | 269 | 410 |
| **340** | 371 | 321 | 282 | 405 | 281 | 360 | 405 | 240 | 302 |
| **266** | 354 | 383 | 265 | 370 | 427 | 265 | 257 | 391 | 368 |
| **364** | 313 | 269 | 357 | 275 | 255 | 274 | 296 | 259 | 255 |
| **353** | 283 | 379 | 297 | 337 | 311 | 372 | 300 | 344 | 347 |
| **329** | 309 | 410 | 402 | 367 | 308 | 308 | 403 | 365 | 321 |
| **278** | 327 | 376 | 303 | 322 | 365 | 362 | 315 | 270 | 339 |
| **302** | 388 | 302 | 343 | 364 | 389 | 319 | 320 | 284 | 344 |
| **303** | 325 | 250 | 293 | 280 | 386 | 283 | 358 | 243 | 319 |
| **255** | 304 | 258 | 397 | 379 | 350 | 391 | 368 | 328 | 346 |
| **293** | 382 | 339 | 249 | 296 | 375 | 325 | 280 | 384 | 345 |
| **283** | 304 | 367 | 285 | 299 | 289 | 286 | 334 | 293 | 325 |
| **300** | 370 | 361 | 389 | 339 | 287 | 304 | 305 | 403 | 374 |
| **335** | 394 | 288 | 339 | 313 | 372 | 360 | 393 | 335 | 282 |
| **385** | 278 | 363 | 365 | 288 | 263 | 398 | 384 | 263 | 346 |
| **332** | 275 | 316 | 313 | 313 | 360 | 380 | 420 | 356 | 347 |
| **262** | 278 | 297 | 322 | 377 | 314 | 360 | 263 | 367 | 362 |
| **403** | 368 | 256 | 275 | 330 | 279 | 268 | 363 | 265 | 369 |
| **313** | 350 | 383 | 273 | 266 | 346 | 364 | 360 | 289 | 266 |
| **351** | 358 | 421 | 316 | 380 | 354 | 267 | 372 | 289 | 345 |
| **312** | 378 | 311 | 380 | 352 | 300 | 301 | 329 | 347 | 358 |
| **327** | 334 | 301 | 363 | 408 | 391 | 388 | 303 | 371 | 325 |
| **327** | 322 | 323 | 354 | 318 | 389 | 330 | 350 | 373 | 397 |
| **409** | 282 | 270 | 255 | 384 | 296 | 279 | 329 | 371 | 303 |
| **271** | 246 | 299 | 410 | 411 | 276 | 368 | 286 | 279 | 409 |
| **284** | 364 | 368 | 281 | 355 | 346 | 271 | 280 | 287 | 306 |
| **343** | 303 | 282 | 255 | 364 | 296 | 333 | 378 | 314 | 385 |
| **313** | 345 | 335 | 307 | 352 | 291 | 367 | 278 | 385 | 273 |
| **402** | 330 | 375 | 276 | 343 | 348 | 338 | 314 | 323 | 375 |
| **303** | 287 | 331 | 347 | 274 | 353 | 367 | 322 | 329 | 325 |
| **370** | 276 | 273 | 264 | 353 | 390 | 316 | 285 | 347 | 301 |

| 320 | 331 | 295 | 356 | 335 | 264 | 284 | 306 | 383 | 289 |
|---|---|---|---|---|---|---|---|---|---|
| **275** | 290 | 371 | 310 | 358 | 393 | 297 | 366 | 361 | 382 |
| **334** | 306 | 368 | 322 | 304 | 296 | 355 | 383 | 286 | 283 |
| **378** | 393 | 368 | 300 | 271 | 327 | 279 | 342 | 353 | 327 |
| **393** | 327 | 383 | 406 | 405 | 275 | 345 | 359 | 279 | 350 |
| **378** | 355 | 301 | 311 | 265 | 425 | 309 | 270 | 337 | 384 |
| **297** | 275 | 376 | 393 | 307 | 405 | 414 | 384 | 315 | 342 |
| **389** | 322 | 380 | 423 | 340 | 278 | 320 | 351 | 376 | 275 |
| **373** | 332 | 331 | 317 | 379 | 363 | 339 | 276 | 275 | 339 |
| **291** | 292 | 328 | 353 | 307 | 253 | 360 | 311 | 327 | 305 |
| **317** | 291 | 348 | 383 | 301 | 366 | 345 | 272 | 307 | 307 |
| **258** | 371 | 328 | 363 | 373 | 359 | 268 | 296 | 379 | 259 |
| **318** | 391 | 259 | 298 | 380 | 303 | 332 | 264 | 362 | 277 |
| **330** | 311 | 378 | 381 | 388 | 369 | 324 | 296 | 306 | 347 |
| **359** | 324 | 273 | 309 | 359 | 295 | 286 | 361 | 303 | 298 |
| **278** | 299 | 295 | 344 | 317 | 267 | 390 | 351 | 261 | 379 |
| **320** | 418 | 307 | 340 | 287 | 350 | 397 | 329 | 317 | 332 |
| **413** | 321 | 276 | 376 | 309 | 392 | 341 | 319 | 351 | 285 |
| **449** | 340 | 294 | 377 | 320 | 342 | 402 | 321 | 340 | 404 |
| **377** | 327 | 369 | 352 | 316 | 304 | 351 | 320 | 434 | 384 |
| **310** | 393 | 347 | 263 | 337 | 388 | 362 | 371 | 295 | 308 |
| **339** | 319 | 328 | 337 | 292 | 300 | 380 | 290 | 384 | 339 |

Results (traditional census method)

| 320 | 331 | 295 | 356 | 335 | 264 | 284 | 306 | 383 | 289 |
|---|---|---|---|---|---|---|---|---|---|
| **277** | 309 | 312 | 331 | 283 | 341 | 328 | 431 | 319 | 267 |
| **268** | 306 | 318 | 261 | 375 | 354 | 383 | 328 | 346 | 367 |
| **415** | 329 | 337 | 280 | 264 | 343 | 296 | 387 | 416 | 317 |
| **363** | 338 | 280 | 337 | 263 | 297 | 289 | 297 | 248 | 304 |
| **361** | 274 | 335 | 287 | 334 | 298 | 287 | 271 | 312 | 324 |
| **340** | 338 | 321 | 337 | 291 | 393 | 317 | 316 | 421 | 319 |
| **352** | 318 | 311 | 341 | 274 | 407 | 388 | 380 | 393 | 310 |
| **417** | 303 | 334 | 406 | 340 | 312 | 387 | 333 | 347 | 393 |
| **324** | 289 | 369 | 323 | 293 | 406 | 372 | 272 | 351 | 399 |
| **304** | 298 | 289 | 322 | 284 | 374 | 270 | 366 | 255 | 253 |
| **330** | 286 | 340 | 300 | 396 | 378 | 336 | 266 | 361 | 419 |
| **302** | 326 | 369 | 347 | 382 | 364 | 374 | 424 | 360 | 334 |
| **255** | 298 | 346 | 363 | 381 | 314 | 274 | 334 | 343 | 405 |
| **300** | 382 | 290 | 365 | 360 | 315 | 326 | 401 | 383 | 366 |
| **374** | 313 | 359 | 397 | 272 | 391 | 398 | 348 | 278 | 294 |
| **319** | 277 | 347 | 378 | 273 | 273 | 354 | 358 | 293 | 278 |
| **389** | 295 | 343 | 274 | 325 | 287 | 283 | 323 | 325 | 336 |
| **275** | 263 | 361 | 304 | 326 | 266 | 260 | 347 | 270 | 343 |
| **264** | 380 | 390 | 378 | 365 | 311 | 329 | 320 | 314 | 275 |
| **340** | 336 | 296 | 361 | 388 | 357 | 318 | 380 | 264 | 396 |
| **327** | 310 | 328 | 279 | 403 | 267 | 369 | 304 | 275 | 291 |
| **272** | 271 | 396 | 354 | 290 | 385 | 332 | 348 | 360 | 340 |
| **285** | 370 | 307 | 265 | 387 | 281 | 392 | 351 | 367 | 294 |
| **281** | 351 | 390 | 310 | 313 | 350 | 284 | 276 | 367 | 312 |
| **311** | 263 | 317 | 351 | 263 | 384 | 390 | 329 | 355 | 274 |
| **306** | 328 | 311 | 373 | 349 | 252 | 383 | 368 | 366 | 299 |
| **334** | 372 | 328 | 328 | 266 | 362 | 292 | 359 | 348 | 290 |
| **337** | 373 | 311 | 306 | 279 | 259 | 322 | 323 | 338 | 309 |
| **261** | 396 | 313 | 313 | 324 | 255 | 328 | 368 | 350 | 397 |
| **370** | 356 | 291 | 258 | 379 | 303 | 327 | 301 | 361 | 282 |
| **310** | 360 | 378 | 340 | 399 | 366 | 317 | 355 | 258 | 309 |
| **388** | 313 | 320 | 357 | 287 | 359 | 273 | 347 | 424 | 305 |

| 320 | 331 | 295 | 356 | 335 | 264 | 284 | 306 | 383 | 289 |
|---|---|---|---|---|---|---|---|---|---|
| **366** | 307 | 358 | 349 | 261 | 315 | 376 | 331 | 275 | 365 |
| **368** | 372 | 306 | 289 | 330 | 257 | 281 | 338 | 263 | 397 |
| **292** | 351 | 329 | 362 | 289 | 297 | 284 | 299 | 357 | 288 |
| **362** | 339 | 364 | 296 | 336 | 426 | 318 | 383 | 331 | 288 |
| **376** | 423 | 382 | 298 | 357 | 379 | 313 | 274 | 388 | 333 |
| **345** | 293 | 348 | 344 | 364 | 273 | 278 | 258 | 354 | 338 |
| **266** | 398 | 310 | 354 | 237 | 299 | 361 | 313 | 362 | 298 |
| **368** | 338 | 341 | 254 | 380 | 262 | 367 | 297 | 353 | 292 |
| **382** | 409 | 327 | 374 | 423 | 305 | 378 | 271 | 347 | 407 |
| **311** | 335 | 313 | 367 | 272 | 293 | 268 | 290 | 331 | 292 |
| **273** | 383 | 270 | 348 | 315 | 322 | 345 | 287 | 253 | 330 |
| **321** | 318 | 286 | 330 | 277 | 271 | 347 | 351 | 310 | 388 |
| **333** | 394 | 365 | 328 | 320 | 269 | 343 | 338 | 261 | 317 |
| **349** | 267 | 296 | 297 | 243 | 273 | 292 | 328 | 263 | 274 |
| **361** | 402 | 256 | 330 | 291 | 267 | 243 | 396 | 368 | 362 |
| **345** | 376 | 315 | 317 | 289 | 269 | 274 | 314 | 387 | 406 |
| **287** | 276 | 291 | 341 | 309 | 312 | 335 | 317 | 330 | 317 |
| **352** | 314 | 313 | 339 | 241 | 369 | 253 | 384 | 357 | 423 |
| **378** | 456 | 300 | 261 | 342 | 311 | 357 | 300 | 360 | 306 |
| **288** | 336 | 398 | 365 | 391 | 321 | 344 | 359 | 341 | 343 |
| **347** | 356 | 413 | 275 | 298 | 373 | 316 | 362 | 359 | 321 |
| **409** | 311 | 357 | 347 | 279 | 398 | 326 | 362 | 304 | 395 |
| **351** | 321 | 386 | 342 | 265 | 348 | 308 | 309 | 385 | 411 |
| **294** | 358 | 365 | 336 | 350 | 350 | 309 | 305 | 319 | 332 |
| **368** | 342 | 277 | 390 | 348 | 275 | 274 | 375 | 388 | 302 |
| **395** | 332 | 299 | 356 | 310 | 428 | 288 | 325 | 312 | 293 |
| **370** | 317 | 306 | 278 | 364 | 343 | 351 | 381 | 318 | 313 |
| **312** | 367 | 383 | 347 | 380 | 322 | 345 | 255 | 292 | 369 |
| **267** | 369 | 370 | 286 | 290 | 371 | 329 | 340 | 325 | 260 |
| **338** | 403 | 387 | 309 | 295 | 247 | 272 | 307 | 294 | 267 |
| **306** | 286 | 267 | 287 | 316 | 274 | 279 | 357 | 273 | 277 |
| **343** | 293 | 284 | 332 | 386 | 279 | 264 | 309 | 285 | 316 |
| **239** | 360 | 421 | 269 | 338 | 291 | 329 | 438 | 295 | 367 |

| 320 | 331 | 295 | 356 | 335 | 264 | 284 | 306 | 383 | 289 |
|---|---|---|---|---|---|---|---|---|---|
| **298** | 393 | 331 | 392 | 339 | 397 | 284 | 370 | 335 | 316 |
| **275** | 340 | 333 | 370 | 323 | 312 | 384 | 322 | 360 | 264 |
| **277** | 332 | 303 | 287 | 284 | 404 | 331 | 276 | 269 | 410 |
| **340** | 371 | 321 | 282 | 405 | 281 | 360 | 405 | 240 | 302 |
| **266** | 354 | 383 | 265 | 370 | 427 | 265 | 257 | 391 | 368 |
| **364** | 313 | 269 | 357 | 275 | 255 | 274 | 296 | 259 | 255 |
| **353** | 283 | 379 | 297 | 337 | 311 | 372 | 300 | 344 | 347 |
| **329** | 309 | 410 | 402 | 367 | 308 | 308 | 403 | 365 | 321 |
| **278** | 327 | 376 | 303 | 322 | 365 | 362 | 315 | 270 | 339 |
| **302** | 388 | 302 | 343 | 364 | 389 | 319 | 320 | 284 | 344 |
| **303** | 325 | 250 | 293 | 280 | 386 | 283 | 358 | 243 | 319 |
| **255** | 304 | 258 | 397 | 379 | 350 | 391 | 368 | 328 | 346 |
| **293** | 382 | 339 | 249 | 296 | 375 | 325 | 280 | 384 | 345 |
| **283** | 304 | 367 | 285 | 299 | 289 | 286 | 334 | 293 | 325 |
| **300** | 370 | 361 | 389 | 339 | 287 | 304 | 305 | 403 | 374 |
| **335** | 394 | 288 | 339 | 313 | 372 | 360 | 393 | 335 | 282 |
| **385** | 278 | 363 | 365 | 288 | 263 | 398 | 384 | 263 | 346 |
| **332** | 275 | 316 | 313 | 313 | 360 | 380 | 420 | 356 | 347 |
| **262** | 278 | 297 | 322 | 377 | 314 | 360 | 263 | 367 | 362 |
| **403** | 368 | 256 | 275 | 330 | 279 | 268 | 363 | 265 | 369 |
| **313** | 350 | 383 | 273 | 266 | 346 | 364 | 360 | 289 | 266 |
| **351** | 358 | 421 | 316 | 380 | 354 | 267 | 372 | 289 | 345 |
| **312** | 378 | 311 | 380 | 352 | 300 | 301 | 329 | 347 | 358 |
| **327** | 334 | 301 | 363 | 408 | 391 | 388 | 303 | 371 | 325 |
| **327** | 322 | 323 | 354 | 318 | 389 | 330 | 350 | 373 | 397 |
| **409** | 282 | 270 | 255 | 384 | 296 | 279 | 329 | 371 | 303 |
| **271** | 246 | 299 | 410 | 411 | 276 | 368 | 286 | 279 | 409 |
| **284** | 364 | 368 | 281 | 355 | 346 | 271 | 280 | 287 | 306 |
| **343** | 303 | 282 | 255 | 364 | 296 | 333 | 378 | 314 | 385 |
| **313** | 345 | 335 | 307 | 352 | 291 | 367 | 278 | 385 | 273 |
| **402** | 330 | 375 | 276 | 343 | 348 | 338 | 314 | 323 | 375 |
| **303** | 287 | 331 | 347 | 274 | 353 | 367 | 322 | 329 | 325 |
| **370** | 276 | 273 | 264 | 353 | 390 | 316 | 285 | 347 | 301 |

| **320** | **331** | **295** | **356** | **335** | **264** | **284** | **306** | **383** | **289** |
|---|---|---|---|---|---|---|---|---|---|
| **275** | 290 | 371 | 310 | 358 | 393 | 297 | 366 | 361 | 382 |
| **334** | 306 | 368 | 322 | 304 | 296 | 355 | 383 | 286 | 283 |
| **378** | 393 | 368 | 300 | 271 | 327 | 279 | 342 | 353 | 327 |
| **393** | 327 | 383 | 406 | 405 | 275 | 345 | 359 | 279 | 350 |
| **378** | 355 | 301 | 311 | 265 | 425 | 309 | 270 | 337 | 384 |
| **297** | 275 | 376 | 393 | 307 | 405 | 414 | 384 | 315 | 342 |
| **389** | 322 | 380 | 423 | 340 | 278 | 320 | 351 | 376 | 275 |
| **373** | 332 | 331 | 317 | 379 | 363 | 339 | 276 | 275 | 339 |
| **291** | 292 | 328 | 353 | 307 | 253 | 360 | 311 | 327 | 305 |
| **317** | 291 | 348 | 383 | 301 | 366 | 345 | 272 | 307 | 307 |
| **258** | 371 | 328 | 363 | 373 | 359 | 268 | 296 | 379 | 259 |
| **318** | 391 | 259 | 298 | 380 | 303 | 332 | 264 | 362 | 277 |
| **330** | 311 | 378 | 381 | 388 | 369 | 324 | 296 | 306 | 347 |
| **359** | 324 | 273 | 309 | 359 | 295 | 286 | 361 | 303 | 298 |
| **278** | 299 | 295 | 344 | 317 | 267 | 390 | 351 | 261 | 379 |
| **320** | 418 | 307 | 340 | 287 | 350 | 397 | 329 | 317 | 332 |
| **413** | 321 | 276 | 376 | 309 | 392 | 341 | 319 | 351 | 285 |
| **449** | 340 | 294 | 377 | 320 | 342 | 402 | 321 | 340 | 404 |
| **377** | 327 | 369 | 352 | 316 | 304 | 351 | 320 | 434 | 384 |
| **310** | 393 | 347 | 263 | 337 | 388 | 362 | 371 | 295 | 308 |
| **339** | 319 | 328 | 337 | 292 | 300 | 380 | 290 | 384 | 339 |

Results (new census method)

| 597 | 412 | 446 | 401 | 469 | 350 | 392 | 672 | 402 | 652 | 407 |
|---|---|---|---|---|---|---|---|---|---|---|
| **315** | 442 | 195 | 215 | 726 | 473 | 551 | 340 | 438 | 443 | 132 |
| **592** | 397 | 485 | 433 | 245 | 493 | 199 | 430 | 347 | 413 | 0.00 |
| **343** | 392 | 536 | 667 | 495 | 597 | 497 | 580 | 617 | 489 | 0.13 |
| **327** | 598 | 632 | 328 | 463 | 492 | 360 | 650 | 539 | 342 | 0.36 |
| **356** | 564 | 562 | 443 | 596 | 483 | 632 | 448 | 502 | 527 | |
| **255** | 545 | 375 | 410 | 311 | 194 | 435 | 204 | 340 | 296 | |
| **396** | 531 | 337 | 430 | 523 | 202 | 290 | 260 | 314 | 469 | |
| **269** | 698 | 410 | 210 | 468 | 342 | 329 | 443 | 573 | 162 | |
| **400** | 240 | 471 | 302 | 437 | 233 | 394 | 306 | 586 | 435 | |
| **455** | 356 | 365 | 642 | 563 | 279 | 479 | 351 | 581 | 582 | |
| **557** | 526 | 342 | 382 | 120 | 208 | 297 | 494 | 346 | 457 | |
| **581** | 710 | 380 | 357 | 245 | 348 | 522 | 472 | 198 | 664 | |
| **213** | 449 | 330 | 531 | 225 | 405 | 621 | 407 | 299 | 439 | |
| **412** | 302 | 222 | 411 | 256 | 361 | 523 | 404 | 329 | 473 | |
| **455** | 455 | 278 | 217 | 243 | 472 | 186 | 257 | 393 | 263 | |
| **461** | 367 | 729 | 401 | 589 | 491 | 426 | 304 | 326 | 254 | |
| **424** | 585 | 509 | 505 | 385 | 493 | 368 | 425 | 164 | 362 | |
| **152** | 384 | 248 | 490 | 418 | 444 | 570 | 495 | 652 | 349 | |
| **539** | 298 | 102 | 291 | 525 | 254 | 358 | 510 | 507 | 343 | |
| **575** | 441 | 356 | 428 | 199 | 566 | 577 | 378 | 424 | 469 | |
| **500** | 644 | 214 | 340 | 401 | 751 | 276 | 213 | 473 | 672 | |
| **486** | 194 | 414 | 292 | 342 | 540 | 382 | 307 | 335 | 243 | |
| **397** | 561 | 319 | 385 | 430 | 346 | 531 | 405 | 190 | 286 | |
| **635** | 213 | 586 | 590 | 368 | 298 | 174 | 605 | 339 | 301 | |
| **580** | 287 | 387 | 534 | 382 | 351 | 139 | 210 | 314 | 439 | |
| **200** | 359 | 514 | 386 | 264 | 562 | 413 | 362 | 236 | 560 | |
| **390** | 163 | 452 | 225 | 450 | 382 | 494 | 482 | 395 | 464 | |
| **304** | 448 | 413 | 440 | 788 | 693 | 760 | 426 | 314 | 384 | |
| **551** | 285 | 606 | 666 | 447 | 440 | 491 | 247 | 408 | 259 | |
| **413** | 410 | 541 | 592 | 254 | 593 | 352 | 292 | 558 | 671 | |
| **465** | 131 | 555 | 498 | 325 | 360 | 463 | 217 | 567 | 511 | |
| **342** | 311 | 395 | 447 | 532 | 258 | 660 | 294 | 504 | 300 | |
| **245** | 494 | 254 | 405 | 189 | 475 | 323 | 376 | 376 | 110 | |
| **407** | 324 | 545 | 487 | 232 | 306 | 598 | 424 | 492 | 432 | |
| **397** | 436 | 507 | 310 | 396 | 352 | 561 | 463 | 409 | 500 | |
| **379** | 522 | 147 | 646 | 299 | 238 | 521 | 246 | 244 | 398 | |

| 597 | 412 | 446 | 401 | 469 | 350 | 392 | 672 | 402 | 652 | 407 |
|---|---|---|---|---|---|---|---|---|---|---|
| **536** | 332 | 315 | 309 | 309 | 192 | 402 | 462 | 358 | 448 | |
| **348** | 385 | 510 | 162 | 173 | 341 | 428 | 438 | 483 | 364 | |
| **304** | 242 | 225 | 282 | 405 | 294 | 403 | 483 | 364 | 689 | |
| **268** | 596 | 328 | 653 | 239 | 321 | 424 | 409 | 332 | 616 | |
| **360** | 286 | 502 | 279 | 232 | 538 | 365 | 392 | 418 | 365 | |
| **535** | 531 | 531 | 296 | 457 | 586 | 335 | 489 | 386 | 398 | |
| **376** | 285 | 510 | 569 | 483 | 423 | 388 | 636 | 401 | 581 | |
| **420** | 643 | 631 | 398 | 270 | 249 | 323 | 299 | 496 | 318 | |
| **597** | 518 | 503 | 382 | 349 | 473 | 396 | 570 | 313 | 291 | |
| **467** | 498 | 538 | 644 | 608 | 268 | 658 | 463 | 355 | 633 | |
| **325** | 453 | 297 | 174 | 130 | 430 | 570 | 262 | 304 | 488 | |
| **391** | 441 | 292 | 276 | 454 | 503 | 316 | 420 | 422 | 288 | |
| **277** | 427 | 349 | 342 | 567 | 508 | 386 | 249 | 389 | 297 | |
| **609** | 224 | 480 | 447 | 393 | 191 | 420 | 191 | 550 | 273 | |
| **375** | 444 | 505 | 523 | 326 | 532 | 390 | 677 | 485 | 499 | |
| **619** | 412 | 322 | 301 | 290 | 348 | 369 | 291 | 440 | 596 | |
| **406** | 390 | 266 | 346 | 349 | 336 | 459 | 328 | 429 | 442 | |
| **308** | 340 | 383 | 397 | 580 | 394 | 199 | 443 | 209 | 284 | |
| **385** | 425 | 552 | 376 | 255 | 406 | 527 | 491 | 319 | 335 | |
| **696** | 358 | 401 | 372 | 371 | 358 | 593 | 479 | 147 | 555 | |
| **233** | 439 | 794 | 393 | 321 | 292 | 620 | 382 | 313 | 614 | |
| **523** | 271 | 290 | 356 | 528 | 275 | 601 | 507 | 454 | 605 | |
| **464** | 203 | 436 | 330 | 581 | 381 | 341 | 416 | 278 | 218 | |
| **313** | 306 | 298 | 429 | 322 | 411 | 250 | 420 | 626 | 326 | |
| **373** | 258 | 302 | 659 | 521 | 108 | 464 | 483 | 413 | 607 | |
| **240** | 231 | 466 | 575 | 341 | 519 | 577 | 238 | 316 | 411 | |
| **552** | 428 | 496 | 664 | 648 | 369 | 172 | 509 | 622 | 271 | |
| **251** | 345 | 340 | 389 | 167 | 703 | 493 | 524 | 316 | 356 | |
| **701** | 416 | 474 | 535 | 211 | 582 | 393 | 499 | 520 | 329 | |
| **459** | 466 | 350 | 342 | 297 | 457 | 591 | 176 | 318 | 469 | |
| **461** | 284 | 294 | 409 | 150 | 664 | 241 | 488 | 330 | 439 | |
| **458** | 479 | 398 | 487 | 595 | 504 | 466 | 442 | 385 | 362 | |
| **311** | 385 | 676 | 309 | 223 | 209 | 481 | 225 | 430 | 316 | |
| **534** | 525 | 223 | 519 | 406 | 242 | 338 | 630 | 471 | 501 | |
| **424** | 293 | 300 | 422 | 183 | 574 | 397 | 477 | 577 | 606 | |
| **301** | 653 | 407 | 552 | 227 | 348 | 499 | 520 | 337 | 556 | |
| **226** | 632 | 390 | 477 | 248 | 604 | 591 | 301 | 282 | 424 | |

| **597** | **412** | **446** | **401** | **469** | **350** | **392** | **672** | **402** | **652** | **407** |
|---|---|---|---|---|---|---|---|---|---|---|
| **749** | 231 | 526 | 483 | 692 | 169 | 585 | 331 | 524 | 413 | |
| **524** | 410 | 412 | 370 | 357 | 450 | 310 | 503 | 660 | 319 | |
| **348** | 471 | 304 | 346 | 490 | 278 | 496 | 227 | 576 | 477 | |
| **467** | 579 | 418 | 161 | 459 | 396 | 357 | 168 | 397 | 257 | |
| **372** | 342 | 431 | 483 | 404 | 360 | 472 | 356 | 304 | 487 | |
| **357** | 488 | 400 | 502 | 306 | 498 | 496 | 408 | 516 | 363 | |
| **587** | 604 | 211 | 561 | 176 | 280 | 421 | 454 | 294 | 306 | |
| **442** | 596 | 365 | 194 | 321 | 301 | 295 | 385 | 481 | 438 | |
| **198** | 458 | 340 | 177 | 323 | 276 | 123 | 354 | 320 | 290 | |
| **397** | 329 | 404 | 681 | 422 | 458 | 446 | 241 | 293 | 580 | |
| **382** | 512 | 447 | 563 | 208 | 323 | 536 | 203 | 343 | 351 | |
| **355** | 269 | 331 | 427 | 358 | 509 | 289 | 270 | 218 | 396 | |
| **348** | 216 | 249 | 631 | 470 | 501 | 264 | 531 | 461 | 300 | |
| **476** | 145 | 490 | 506 | 442 | 359 | 493 | 480 | 416 | 339 | |
| **246** | 332 | 436 | 401 | 465 | 348 | 384 | 547 | 431 | 448 | |
| **587** | 459 | 627 | 434 | 334 | 296 | 335 | 465 | 264 | 506 | |
| **624** | 413 | 655 | 192 | 122 | 373 | 466 | 250 | 309 | 297 | |
| **343** | 656 | 461 | 684 | 446 | 510 | 597 | 422 | 443 | 579 | |
| **688** | 603 | 317 | 334 | 327 | 455 | 271 | 447 | 717 | 488 | |
| **352** | 127 | 244 | 549 | 285 | 212 | 705 | 488 | 311 | 464 | |
| **246** | 547 | 345 | 561 | 257 | 736 | 456 | 304 | 232 | 663 | |
| **236** | 694 | 286 | 247 | 413 | 407 | 278 | 586 | 250 | 677 | |
| **206** | 342 | 434 | 381 | 570 | 458 | 420 | 427 | 552 | 266 | |
| **447** | 314 | 349 | 184 | 497 | 449 | 366 | 463 | 214 | 448 | |
| **399** | 269 | 402 | 455 | 273 | 354 | 403 | 585 | 480 | 268 | |
| **264** | 400 | 330 | 176 | 340 | 234 | 566 | 419 | 205 | 451 | |
| **462** | 497 | 334 | 181 | 282 | 337 | 278 | 374 | 528 | 453 | |
| **213** | 240 | 368 | 419 | 396 | 685 | 739 | 498 | 442 | 398 | |
| **359** | 437 | 278 | 398 | 240 | 687 | 299 | 308 | 383 | 280 | |
| **449** | 374 | 582 | 211 | 275 | 450 | 383 | 497 | 425 | 204 | |
| **645** | 464 | 117 | 390 | 525 | 324 | 385 | 400 | 384 | 405 | |
| **267** | 684 | 445 | 346 | 158 | 580 | 263 | 293 | 541 | 517 | |
| **132** | 491 | 482 | 731 | 220 | 371 | 326 | 208 | 636 | 206 | |
| **501** | 513 | 539 | 499 | 310 | 224 | 340 | 509 | 569 | 527 | |
| **357** | 379 | 213 | 418 | 219 | 299 | 284 | 441 | 336 | 528 | |
| **437** | 219 | 348 | 373 | 420 | 245 | 442 | 421 | 286 | 280 | |
| **192** | 237 | 465 | 345 | 308 | 349 | 266 | 467 | 367 | 348 | |

| **597** | **412** | **446** | **401** | **469** | **350** | **392** | **672** | **402** | **652** | **407** |
|---|---|---|---|---|---|---|---|---|---|---|
| **400** | 671 | 250 | 357 | 476 | 391 | 375 | 437 | 472 | 454 | |
| **348** | 137 | 246 | 501 | 309 | 723 | 590 | 602 | 434 | 425 | |
| **613** | 455 | 198 | 361 | 505 | 653 | 463 | 336 | 329 | 371 | |
| **275** | 537 | 379 | 449 | 398 | 611 | 522 | 560 | 531 | 524 | |
| **436** | 569 | 527 | 369 | 519 | 283 | 448 | 317 | 317 | 518 | |
| **388** | 294 | 345 | 200 | 308 | 445 | 522 | 504 | 393 | 371 | |
| **317** | 631 | 378 | 267 | 291 | 483 | 336 | 420 | 321 | 338 | |
| **498** | 395 | 509 | 798 | 229 | 195 | 477 | 398 | 440 | 227 | |
| **299** | 466 | 525 | 437 | 415 | 512 | 383 | 305 | 189 | 259 | |

Pairs of values N2022 v H user-defined

| 320 | 74 |
|---|---|
| 277 | 74 |
| 268 | 74 |
| 415 | 74 |
| 363 | 74 |
| 361 | 74 |
| 340 | 74 |
| 352 | 74 |
| 417 | 74 |
| 324 | 74 |
| 304 | 74 |
| 330 | 74 |
| 302 | 74 |
| 255 | 74 |
| 300 | 74 |
| 374 | 74 |
| 319 | 74 |
| 389 | 74 |
| 275 | 74 |
| 264 | 74 |
| 340 | 74 |
| 327 | 74 |
| 272 | 74 |
| 285 | 74 |
| 281 | 74 |
| 311 | 74 |
| 306 | 74 |
| 334 | 74 |
| 337 | 74 |
| 261 | 74 |
| 370 | 74 |

| 320 | 74 |
|---|---|
| 310 | 74 |
| 388 | 74 |
| 366 | 74 |
| 368 | 74 |
| 292 | 74 |
| 362 | 74 |
| 376 | 74 |
| 345 | 74 |
| 266 | 74 |
| 368 | 74 |
| 382 | 74 |
| 311 | 74 |
| 273 | 74 |
| 321 | 74 |
| 333 | 74 |
| 349 | 74 |
| 361 | 74 |
| 345 | 74 |
| 287 | 74 |
| 352 | 74 |
| 378 | 74 |
| 288 | 74 |
| 347 | 74 |
| 409 | 74 |
| 351 | 74 |
| 294 | 74 |
| 368 | 74 |
| 395 | 74 |
| 370 | 74 |
| 312 | 74 |
| 267 | 74 |

| | |
|---|---|
| **320** | **74** |
| **338** | 74 |
| **306** | 74 |
| **343** | 74 |
| **239** | 74 |
| **298** | 74 |
| **275** | 74 |
| **277** | 74 |
| **340** | 74 |
| **266** | 74 |
| **364** | 74 |
| **353** | 74 |
| **329** | 74 |
| **278** | 74 |
| **302** | 74 |
| **303** | 74 |
| **255** | 74 |
| **293** | 74 |
| **283** | 74 |
| **300** | 74 |
| **335** | 74 |
| **385** | 74 |
| **332** | 74 |
| **262** | 74 |
| **403** | 74 |
| **313** | 74 |
| **351** | 74 |
| **312** | 74 |
| **327** | 74 |
| **327** | 74 |
| **409** | 74 |
| **271** | 74 |

| 320 | 74 |
|---|---|
| 284 | 74 |
| 343 | 74 |
| 313 | 74 |
| 402 | 74 |
| 303 | 74 |
| 370 | 74 |
| 275 | 74 |
| 334 | 74 |
| 378 | 74 |
| 393 | 74 |
| 378 | 74 |
| 297 | 74 |
| 389 | 74 |
| 373 | 74 |
| 291 | 74 |
| 317 | 74 |
| 258 | 74 |
| 318 | 74 |
| 330 | 74 |
| 359 | 74 |
| 278 | 74 |
| 320 | 74 |
| 413 | 74 |
| 449 | 74 |
| 377 | 74 |
| 310 | 74 |
| 339 | 74 |
| 331 | 74 |
| 309 | 74 |
| 306 | 74 |
| 329 | 74 |

| | |
|---|---|
| **320** | **74** |
| **338** | 74 |
| **274** | 74 |
| **338** | 74 |
| **318** | 74 |
| **303** | 74 |
| **289** | 74 |
| **298** | 74 |
| **286** | 74 |
| **326** | 74 |
| **298** | 74 |
| **382** | 74 |
| **313** | 74 |
| **277** | 74 |
| **295** | 74 |
| **263** | 74 |
| **380** | 74 |
| **336** | 74 |
| **310** | 74 |
| **271** | 74 |
| **370** | 74 |
| **351** | 74 |
| **263** | 74 |
| **328** | 74 |
| **372** | 74 |
| **373** | 74 |
| **396** | 74 |
| **356** | 74 |
| **360** | 74 |
| **313** | 74 |
| **307** | 74 |
| **372** | 74 |

| 320 | 74 |
|---|---|
| 351 | 74 |
| 339 | 74 |
| 423 | 74 |
| 293 | 74 |
| 398 | 74 |
| 338 | 74 |
| 409 | 74 |
| 335 | 74 |
| 383 | 74 |
| 318 | 74 |
| 394 | 74 |
| 267 | 74 |
| 402 | 74 |
| 376 | 74 |
| 276 | 74 |
| 314 | 74 |
| 456 | 74 |
| 336 | 74 |
| 356 | 74 |
| 311 | 74 |
| 321 | 74 |
| 358 | 74 |
| 342 | 74 |
| 332 | 74 |
| 317 | 74 |
| 367 | 74 |
| 369 | 74 |
| 403 | 74 |
| 286 | 74 |
| 293 | 74 |
| 360 | 74 |

| | |
|---|---|
| **320** | **74** |
| **393** | 74 |
| **340** | 74 |
| **332** | 74 |
| **371** | 74 |
| **354** | 74 |
| **313** | 74 |
| **283** | 74 |
| **309** | 74 |
| **327** | 74 |
| **388** | 74 |
| **325** | 74 |
| **304** | 74 |
| **382** | 74 |
| **304** | 74 |
| **370** | 74 |
| **394** | 74 |
| **278** | 74 |
| **275** | 74 |
| **278** | 74 |
| **368** | 74 |
| **350** | 74 |
| **358** | 74 |
| **378** | 74 |
| **334** | 74 |
| **322** | 74 |
| **282** | 74 |
| **246** | 74 |
| **364** | 74 |
| **303** | 74 |
| **345** | 74 |
| **330** | 74 |

| 320 | 74 |
|---|---|
| 287 | 74 |
| 276 | 74 |
| 290 | 74 |
| 306 | 74 |
| 393 | 74 |
| 327 | 74 |
| 355 | 74 |
| 275 | 74 |
| 322 | 74 |
| 332 | 74 |
| 292 | 74 |
| 291 | 74 |
| 371 | 74 |
| 391 | 74 |
| 311 | 74 |
| 324 | 74 |
| 299 | 74 |
| 418 | 74 |
| 321 | 74 |
| 340 | 74 |
| 327 | 74 |
| 393 | 74 |
| 319 | 74 |
| 295 | 74 |
| 312 | 74 |
| 318 | 74 |
| 337 | 74 |
| 280 | 74 |
| 335 | 74 |
| 321 | 74 |
| 311 | 74 |

| | |
|---|---|
| **320** | **74** |
| **334** | 74 |
| **369** | 74 |
| **289** | 74 |
| **340** | 74 |
| **369** | 74 |
| **346** | 74 |
| **290** | 74 |
| **359** | 74 |
| **347** | 74 |
| **343** | 74 |
| **361** | 74 |
| **390** | 74 |
| **296** | 74 |
| **328** | 74 |
| **396** | 74 |
| **307** | 74 |
| **390** | 74 |
| **317** | 74 |
| **311** | 74 |
| **328** | 74 |
| **311** | 74 |
| **313** | 74 |
| **291** | 74 |
| **378** | 74 |
| **320** | 74 |
| **358** | 74 |
| **306** | 74 |
| **329** | 74 |
| **364** | 74 |
| **382** | 74 |
| **348** | 74 |

| 320 | 74 |
|---|---|
| 310 | 74 |
| 341 | 74 |
| 327 | 74 |
| 313 | 74 |
| 270 | 74 |
| 286 | 74 |
| 365 | 74 |
| 296 | 74 |
| 256 | 74 |
| 315 | 74 |
| 291 | 74 |
| 313 | 74 |
| 300 | 74 |
| 398 | 74 |
| 413 | 74 |
| 357 | 74 |
| 386 | 74 |
| 365 | 74 |
| 277 | 74 |
| 299 | 74 |
| 306 | 74 |
| 383 | 74 |
| 370 | 74 |
| 387 | 74 |
| 267 | 74 |
| 284 | 74 |
| 421 | 74 |
| 331 | 74 |
| 333 | 74 |
| 303 | 74 |
| 321 | 74 |

| 320 | 74 |
|---|---|
| 383 | 74 |
| 269 | 74 |
| 379 | 74 |
| 410 | 74 |
| 376 | 74 |
| 302 | 74 |
| 250 | 74 |
| 258 | 74 |
| 339 | 74 |
| 367 | 74 |
| 361 | 74 |
| 288 | 74 |
| 363 | 74 |
| 316 | 74 |
| 297 | 74 |
| 256 | 74 |
| 383 | 74 |
| 421 | 74 |
| 311 | 74 |
| 301 | 74 |
| 323 | 74 |
| 270 | 74 |
| 299 | 74 |
| 368 | 74 |
| 282 | 74 |
| 335 | 74 |
| 375 | 74 |
| 331 | 74 |
| 273 | 74 |
| 371 | 74 |
| 368 | 74 |

| | |
|---|---|
| **320** | **74** |
| **368** | 74 |
| **383** | 74 |
| **301** | 74 |
| **376** | 74 |
| **380** | 74 |
| **331** | 74 |
| **328** | 74 |
| **348** | 74 |
| **328** | 74 |
| **259** | 74 |
| **378** | 74 |
| **273** | 74 |
| **295** | 74 |
| **307** | 74 |
| **276** | 74 |
| **294** | 74 |
| **369** | 74 |
| **347** | 74 |
| **328** | 74 |
| **356** | 74 |
| **331** | 74 |
| **261** | 74 |
| **280** | 74 |
| **337** | 74 |
| **287** | 74 |
| **337** | 74 |
| **341** | 74 |
| **406** | 74 |
| **323** | 74 |
| **322** | 74 |
| **300** | 74 |

| | |
|---|---|
| **320** | **74** |
| **347** | 74 |
| **363** | 74 |
| **365** | 74 |
| **397** | 74 |
| **378** | 74 |
| **274** | 74 |
| **304** | 74 |
| **378** | 74 |
| **361** | 74 |
| **279** | 74 |
| **354** | 74 |
| **265** | 74 |
| **310** | 74 |
| **351** | 74 |
| **373** | 74 |
| **328** | 74 |
| **306** | 74 |
| **313** | 74 |
| **258** | 74 |
| **340** | 74 |
| **357** | 74 |
| **349** | 74 |
| **289** | 74 |
| **362** | 74 |
| **296** | 74 |
| **298** | 74 |
| **344** | 74 |
| **354** | 74 |
| **254** | 74 |
| **374** | 74 |
| **367** | 74 |

| | |
|---|---|
| **320** | **74** |
| **348** | 74 |
| **330** | 74 |
| **328** | 74 |
| **297** | 74 |
| **330** | 74 |
| **317** | 74 |
| **341** | 74 |
| **339** | 74 |
| **261** | 74 |
| **365** | 74 |
| **275** | 74 |
| **347** | 74 |
| **342** | 74 |
| **336** | 74 |
| **390** | 74 |
| **356** | 74 |
| **278** | 74 |
| **347** | 74 |
| **286** | 74 |
| **309** | 74 |
| **287** | 74 |
| **332** | 74 |
| **269** | 74 |
| **392** | 74 |
| **370** | 74 |
| **287** | 74 |
| **282** | 74 |
| **265** | 74 |
| **357** | 74 |
| **297** | 74 |
| **402** | 74 |

| | |
|---|---|
| **320** | **74** |
| **303** | 74 |
| **343** | 74 |
| **293** | 74 |
| **397** | 74 |
| **249** | 74 |
| **285** | 74 |
| **389** | 74 |
| **339** | 74 |
| **365** | 74 |
| **313** | 74 |
| **322** | 74 |
| **275** | 74 |
| **273** | 74 |
| **316** | 74 |
| **380** | 74 |
| **363** | 74 |
| **354** | 74 |
| **255** | 74 |
| **410** | 74 |
| **281** | 74 |
| **255** | 74 |
| **307** | 74 |
| **276** | 74 |
| **347** | 74 |
| **264** | 74 |
| **310** | 74 |
| **322** | 74 |
| **300** | 74 |
| **406** | 74 |
| **311** | 74 |
| **393** | 74 |

| | |
|---|---|
| **320** | **74** |
| **423** | 74 |
| **317** | 74 |
| **353** | 74 |
| **383** | 74 |
| **363** | 74 |
| **298** | 74 |
| **381** | 74 |
| **309** | 74 |
| **344** | 74 |
| **340** | 74 |
| **376** | 74 |
| **377** | 74 |
| **352** | 74 |
| **263** | 74 |
| **337** | 74 |
| **335** | 74 |
| **283** | 74 |
| **375** | 74 |
| **264** | 74 |
| **263** | 74 |
| **334** | 74 |
| **291** | 74 |
| **274** | 74 |
| **340** | 74 |
| **293** | 74 |
| **284** | 74 |
| **396** | 74 |
| **382** | 74 |
| **381** | 74 |
| **360** | 74 |
| **272** | 74 |

| 320 | 74 |
|---|---|
| 273 | 74 |
| 325 | 74 |
| 326 | 74 |
| 365 | 74 |
| 388 | 74 |
| 403 | 74 |
| 290 | 74 |
| 387 | 74 |
| 313 | 74 |
| 263 | 74 |
| 349 | 74 |
| 266 | 74 |
| 279 | 74 |
| 324 | 74 |
| 379 | 74 |
| 399 | 74 |
| 287 | 74 |
| 261 | 74 |
| 330 | 74 |
| 289 | 74 |
| 336 | 74 |
| 357 | 74 |
| 364 | 74 |
| 237 | 74 |
| 380 | 74 |
| 423 | 74 |
| 272 | 74 |
| 315 | 74 |
| 277 | 74 |
| 320 | 74 |
| 243 | 74 |

| 320 | 74 |
|---|---|
| 291 | 74 |
| 289 | 74 |
| 309 | 74 |
| 241 | 74 |
| 342 | 74 |
| 391 | 74 |
| 298 | 74 |
| 279 | 74 |
| 265 | 74 |
| 350 | 74 |
| 348 | 74 |
| 310 | 74 |
| 364 | 74 |
| 380 | 74 |
| 290 | 74 |
| 295 | 74 |
| 316 | 74 |
| 386 | 74 |
| 338 | 74 |
| 339 | 74 |
| 323 | 74 |
| 284 | 74 |
| 405 | 74 |
| 370 | 74 |
| 275 | 74 |
| 337 | 74 |
| 367 | 74 |
| 322 | 74 |
| 364 | 74 |
| 280 | 74 |
| 379 | 74 |

| 320 | 74 |
| --- | --- |
| 296 | 74 |
| 299 | 74 |
| 339 | 74 |
| 313 | 74 |
| 288 | 74 |
| 313 | 74 |
| 377 | 74 |
| 330 | 74 |
| 266 | 74 |
| 380 | 74 |
| 352 | 74 |
| 408 | 74 |
| 318 | 74 |
| 384 | 74 |
| 411 | 74 |
| 355 | 74 |
| 364 | 74 |
| 352 | 74 |
| 343 | 74 |
| 274 | 74 |
| 353 | 74 |
| 358 | 74 |
| 304 | 74 |
| 271 | 74 |
| 405 | 74 |
| 265 | 74 |
| 307 | 74 |
| 340 | 74 |
| 379 | 74 |
| 307 | 74 |
| 301 | 74 |

| | |
|---|---|
| **320** | **74** |
| **373** | 74 |
| **380** | 74 |
| **388** | 74 |
| **359** | 74 |
| **317** | 74 |
| **287** | 74 |
| **309** | 74 |
| **320** | 74 |
| **316** | 74 |
| **337** | 74 |
| **292** | 74 |
| **264** | 74 |
| **341** | 74 |
| **354** | 74 |
| **343** | 74 |
| **297** | 74 |
| **298** | 74 |
| **393** | 74 |
| **407** | 74 |
| **312** | 74 |
| **406** | 74 |
| **374** | 74 |
| **378** | 74 |
| **364** | 74 |
| **314** | 74 |
| **315** | 74 |
| **391** | 74 |
| **273** | 74 |
| **287** | 74 |
| **266** | 74 |
| **311** | 74 |

| | |
|---|---|
| **320** | **74** |
| **357** | 74 |
| **267** | 74 |
| **385** | 74 |
| **281** | 74 |
| **350** | 74 |
| **384** | 74 |
| **252** | 74 |
| **362** | 74 |
| **259** | 74 |
| **255** | 74 |
| **303** | 74 |
| **366** | 74 |
| **359** | 74 |
| **315** | 74 |
| **257** | 74 |
| **297** | 74 |
| **426** | 74 |
| **379** | 74 |
| **273** | 74 |
| **299** | 74 |
| **262** | 74 |
| **305** | 74 |
| **293** | 74 |
| **322** | 74 |
| **271** | 74 |
| **269** | 74 |
| **273** | 74 |
| **267** | 74 |
| **269** | 74 |
| **312** | 74 |
| **369** | 74 |

| | |
|---|---|
| **320** | **74** |
| **311** | 74 |
| **321** | 74 |
| **373** | 74 |
| **398** | 74 |
| **348** | 74 |
| **350** | 74 |
| **275** | 74 |
| **428** | 74 |
| **343** | 74 |
| **322** | 74 |
| **371** | 74 |
| **247** | 74 |
| **274** | 74 |
| **279** | 74 |
| **291** | 74 |
| **397** | 74 |
| **312** | 74 |
| **404** | 74 |
| **281** | 74 |
| **427** | 74 |
| **255** | 74 |
| **311** | 74 |
| **308** | 74 |
| **365** | 74 |
| **389** | 74 |
| **386** | 74 |
| **350** | 74 |
| **375** | 74 |
| **289** | 74 |
| **287** | 74 |
| **372** | 74 |

| 320 | 74 |
|---|---|
| 263 | 74 |
| 360 | 74 |
| 314 | 74 |
| 279 | 74 |
| 346 | 74 |
| 354 | 74 |
| 300 | 74 |
| 391 | 74 |
| 389 | 74 |
| 296 | 74 |
| 276 | 74 |
| 346 | 74 |
| 296 | 74 |
| 291 | 74 |
| 348 | 74 |
| 353 | 74 |
| 390 | 74 |
| 393 | 74 |
| 296 | 74 |
| 327 | 74 |
| 275 | 74 |
| 425 | 74 |
| 405 | 74 |
| 278 | 74 |
| 363 | 74 |
| 253 | 74 |
| 366 | 74 |
| 359 | 74 |
| 303 | 74 |
| 369 | 74 |
| 295 | 74 |

| 320 | 74 |
|---|---|
| 267 | 74 |
| 350 | 74 |
| 392 | 74 |
| 342 | 74 |
| 304 | 74 |
| 388 | 74 |
| 300 | 74 |
| 284 | 74 |
| 328 | 74 |
| 383 | 74 |
| 296 | 74 |
| 289 | 74 |
| 287 | 74 |
| 317 | 74 |
| 388 | 74 |
| 387 | 74 |
| 372 | 74 |
| 270 | 74 |
| 336 | 74 |
| 374 | 74 |
| 274 | 74 |
| 326 | 74 |
| 398 | 74 |
| 354 | 74 |
| 283 | 74 |
| 260 | 74 |
| 329 | 74 |
| 318 | 74 |
| 369 | 74 |
| 332 | 74 |
| 392 | 74 |

| | |
|---|---|
| **320** | **74** |
| **284** | 74 |
| **390** | 74 |
| **383** | 74 |
| **292** | 74 |
| **322** | 74 |
| **328** | 74 |
| **327** | 74 |
| **317** | 74 |
| **273** | 74 |
| **376** | 74 |
| **281** | 74 |
| **284** | 74 |
| **318** | 74 |
| **313** | 74 |
| **278** | 74 |
| **361** | 74 |
| **367** | 74 |
| **378** | 74 |
| **268** | 74 |
| **345** | 74 |
| **347** | 74 |
| **343** | 74 |
| **292** | 74 |
| **243** | 74 |
| **274** | 74 |
| **335** | 74 |
| **253** | 74 |
| **357** | 74 |
| **344** | 74 |
| **316** | 74 |
| **326** | 74 |

| | |
|---|---|
| **320** | **74** |
| **308** | 74 |
| **309** | 74 |
| **274** | 74 |
| **288** | 74 |
| **351** | 74 |
| **345** | 74 |
| **329** | 74 |
| **272** | 74 |
| **279** | 74 |
| **264** | 74 |
| **329** | 74 |
| **284** | 74 |
| **384** | 74 |
| **331** | 74 |
| **360** | 74 |
| **265** | 74 |
| **274** | 74 |
| **372** | 74 |
| **308** | 74 |
| **362** | 74 |
| **319** | 74 |
| **283** | 74 |
| **391** | 74 |
| **325** | 74 |
| **286** | 74 |
| **304** | 74 |
| **360** | 74 |
| **398** | 74 |
| **380** | 74 |
| **360** | 74 |
| **268** | 74 |

| 320 | 74 |
|---|---|
| 364 | 74 |
| 267 | 74 |
| 301 | 74 |
| 388 | 74 |
| 330 | 74 |
| 279 | 74 |
| 368 | 74 |
| 271 | 74 |
| 333 | 74 |
| 367 | 74 |
| 338 | 74 |
| 367 | 74 |
| 316 | 74 |
| 297 | 74 |
| 355 | 74 |
| 279 | 74 |
| 345 | 74 |
| 309 | 74 |
| 414 | 74 |
| 320 | 74 |
| 339 | 74 |
| 360 | 74 |
| 345 | 74 |
| 268 | 74 |
| 332 | 74 |
| 324 | 74 |
| 286 | 74 |
| 390 | 74 |
| 397 | 74 |
| 341 | 74 |
| 402 | 74 |

| | |
|---|---|
| **320** | **74** |
| **351** | 74 |
| **362** | 74 |
| **380** | 74 |
| **306** | 74 |
| **431** | 74 |
| **328** | 74 |
| **387** | 74 |
| **297** | 74 |
| **271** | 74 |
| **316** | 74 |
| **380** | 74 |
| **333** | 74 |
| **272** | 74 |
| **366** | 74 |
| **266** | 74 |
| **424** | 74 |
| **334** | 74 |
| **401** | 74 |
| **348** | 74 |
| **358** | 74 |
| **323** | 74 |
| **347** | 74 |
| **320** | 74 |
| **380** | 74 |
| **304** | 74 |
| **348** | 74 |
| **351** | 74 |
| **276** | 74 |
| **329** | 74 |
| **368** | 74 |
| **359** | 74 |

| | |
|---|---|
| **320** | **74** |
| **323** | 74 |
| **368** | 74 |
| **301** | 74 |
| **355** | 74 |
| **347** | 74 |
| **331** | 74 |
| **338** | 74 |
| **299** | 74 |
| **383** | 74 |
| **274** | 74 |
| **258** | 74 |
| **313** | 74 |
| **297** | 74 |
| **271** | 74 |
| **290** | 74 |
| **287** | 74 |
| **351** | 74 |
| **338** | 74 |
| **328** | 74 |
| **396** | 74 |
| **314** | 74 |
| **317** | 74 |
| **384** | 74 |
| **300** | 74 |
| **359** | 74 |
| **362** | 74 |
| **362** | 74 |
| **309** | 74 |
| **305** | 74 |
| **375** | 74 |
| **325** | 74 |

| 320 | 74 |
|---|---|
| 381 | 74 |
| 255 | 74 |
| 340 | 74 |
| 307 | 74 |
| 357 | 74 |
| 309 | 74 |
| 438 | 74 |
| 370 | 74 |
| 322 | 74 |
| 276 | 74 |
| 405 | 74 |
| 257 | 74 |
| 296 | 74 |
| 300 | 74 |
| 403 | 74 |
| 315 | 74 |
| 320 | 74 |
| 358 | 74 |
| 368 | 74 |
| 280 | 74 |
| 334 | 74 |
| 305 | 74 |
| 393 | 74 |
| 384 | 74 |
| 420 | 74 |
| 263 | 74 |
| 363 | 74 |
| 360 | 74 |
| 372 | 74 |
| 329 | 74 |
| 303 | 74 |

| | |
|---|---|
| **320** | **74** |
| **350** | 74 |
| **329** | 74 |
| **286** | 74 |
| **280** | 74 |
| **378** | 74 |
| **278** | 74 |
| **314** | 74 |
| **322** | 74 |
| **285** | 74 |
| **366** | 74 |
| **383** | 74 |
| **342** | 74 |
| **359** | 74 |
| **270** | 74 |
| **384** | 74 |
| **351** | 74 |
| **276** | 74 |
| **311** | 74 |
| **272** | 74 |
| **296** | 74 |
| **264** | 74 |
| **296** | 74 |
| **361** | 74 |
| **351** | 74 |
| **329** | 74 |
| **319** | 74 |
| **321** | 74 |
| **320** | 74 |
| **371** | 74 |
| **290** | 74 |
| **383** | 74 |

| 320 | 74 |
|---|---|
| 319 | 74 |
| 346 | 74 |
| 416 | 74 |
| 248 | 74 |
| 312 | 74 |
| 421 | 74 |
| 393 | 74 |
| 347 | 74 |
| 351 | 74 |
| 255 | 74 |
| 361 | 74 |
| 360 | 74 |
| 343 | 74 |
| 383 | 74 |
| 278 | 74 |
| 293 | 74 |
| 325 | 74 |
| 270 | 74 |
| 314 | 74 |
| 264 | 74 |
| 275 | 74 |
| 360 | 74 |
| 367 | 74 |
| 367 | 74 |
| 355 | 74 |
| 366 | 74 |
| 348 | 74 |
| 338 | 74 |
| 350 | 74 |
| 361 | 74 |
| 258 | 74 |

| | |
|---|---|
| **320** | **74** |
| **424** | 74 |
| **275** | 74 |
| **263** | 74 |
| **357** | 74 |
| **331** | 74 |
| **388** | 74 |
| **354** | 74 |
| **362** | 74 |
| **353** | 74 |
| **347** | 74 |
| **331** | 74 |
| **253** | 74 |
| **310** | 74 |
| **261** | 74 |
| **263** | 74 |
| **368** | 74 |
| **387** | 74 |
| **330** | 74 |
| **357** | 74 |
| **360** | 74 |
| **341** | 74 |
| **359** | 74 |
| **304** | 74 |
| **385** | 74 |
| **319** | 74 |
| **388** | 74 |
| **312** | 74 |
| **318** | 74 |
| **292** | 74 |
| **325** | 74 |
| **294** | 74 |

| 320 | 74 |
| --- | --- |
| 273 | 74 |
| 285 | 74 |
| 295 | 74 |
| 335 | 74 |
| 360 | 74 |
| 269 | 74 |
| 240 | 74 |
| 391 | 74 |
| 259 | 74 |
| 344 | 74 |
| 365 | 74 |
| 270 | 74 |
| 284 | 74 |
| 243 | 74 |
| 328 | 74 |
| 384 | 74 |
| 293 | 74 |
| 403 | 74 |
| 335 | 74 |
| 263 | 74 |
| 356 | 74 |
| 367 | 74 |
| 265 | 74 |
| 289 | 74 |
| 289 | 74 |
| 347 | 74 |
| 371 | 74 |
| 373 | 74 |
| 371 | 74 |
| 279 | 74 |
| 287 | 74 |

| | |
|---|---|
| **320** | **74** |
| **314** | 74 |
| **385** | 74 |
| **323** | 74 |
| **329** | 74 |
| **347** | 74 |
| **361** | 74 |
| **286** | 74 |
| **353** | 74 |
| **279** | 74 |
| **337** | 74 |
| **315** | 74 |
| **376** | 74 |
| **275** | 74 |
| **327** | 74 |
| **307** | 74 |
| **379** | 74 |
| **362** | 74 |
| **306** | 74 |
| **303** | 74 |
| **261** | 74 |
| **317** | 74 |
| **351** | 74 |
| **340** | 74 |
| **434** | 74 |
| **295** | 74 |
| **384** | 74 |
| **289** | 74 |
| **267** | 74 |
| **367** | 74 |
| **317** | 74 |
| **304** | 74 |

| | |
|---|---|
| **320** | **74** |
| **324** | 74 |
| **319** | 74 |
| **310** | 74 |
| **393** | 74 |
| **399** | 74 |
| **253** | 74 |
| **419** | 74 |
| **334** | 74 |
| **405** | 74 |
| **366** | 74 |
| **294** | 74 |
| **278** | 74 |
| **336** | 74 |
| **343** | 74 |
| **275** | 74 |
| **396** | 74 |
| **291** | 74 |
| **340** | 74 |
| **294** | 74 |
| **312** | 74 |
| **274** | 74 |
| **299** | 74 |
| **290** | 74 |
| **309** | 74 |
| **397** | 74 |
| **282** | 74 |
| **309** | 74 |
| **305** | 74 |
| **365** | 74 |
| **397** | 74 |
| **288** | 74 |

| | |
|---|---|
| **320** | **74** |
| **288** | 74 |
| **333** | 74 |
| **338** | 74 |
| **298** | 74 |
| **292** | 74 |
| **407** | 74 |
| **292** | 74 |
| **330** | 74 |
| **388** | 74 |
| **317** | 74 |
| **274** | 74 |
| **362** | 74 |
| **406** | 74 |
| **317** | 74 |
| **423** | 74 |
| **306** | 74 |
| **343** | 74 |
| **321** | 74 |
| **395** | 74 |
| **411** | 74 |
| **332** | 74 |
| **302** | 74 |
| **293** | 74 |
| **313** | 74 |
| **369** | 74 |
| **260** | 74 |
| **267** | 74 |
| **277** | 74 |
| **316** | 74 |
| **367** | 74 |
| **316** | 74 |

| | |
|---|---|
| **320** | **74** |
| **264** | 74 |
| **410** | 74 |
| **302** | 74 |
| **368** | 74 |
| **255** | 74 |
| **347** | 74 |
| **321** | 74 |
| **339** | 74 |
| **344** | 74 |
| **319** | 74 |
| **346** | 74 |
| **345** | 74 |
| **325** | 74 |
| **374** | 74 |
| **282** | 74 |
| **346** | 74 |
| **347** | 74 |
| **362** | 74 |
| **369** | 74 |
| **266** | 74 |
| **345** | 74 |
| **358** | 74 |
| **325** | 74 |
| **397** | 74 |
| **303** | 74 |
| **409** | 74 |
| **306** | 74 |
| **385** | 74 |
| **273** | 74 |
| **375** | 74 |
| **325** | 74 |

| | |
|---|---|
| **320** | **74** |
| **301** | 74 |
| **382** | 74 |
| **283** | 74 |
| **327** | 74 |
| **350** | 74 |
| **384** | 74 |
| **342** | 74 |
| **275** | 74 |
| **339** | 74 |
| **305** | 74 |
| **307** | 74 |
| **259** | 74 |
| **277** | 74 |
| **347** | 74 |
| **298** | 74 |
| **379** | 74 |
| **332** | 74 |
| **285** | 74 |
| **404** | 74 |
| **384** | 74 |
| **308** | 74 |
| **339** | 74 |
