## Supplementary material for "Uncertainty and precaution in hunting wolves twice in a year": Supp Info S3

Deaths uniform distribution between 0.38-0.56

| **RANDBETWEEN(38,56)÷100** |
|---|

| RANDBETWEEN(38,56)÷100 |
|---|

Packs with pups normal centered 0.72 (range 0.55-0.89)

| (((RANDBETWEEN(1,17)+RANDBETWEEN(1,17))−2)÷100)+0.52 |
|---|

**(((RANDBETWEEN(1,17)+RANDBETWEEN(1,17))−2)÷100)+0.52**

Litter size 4.8 (3-6) normal centered on 4.8

| TRUNC(((RANDBETWEEN(3,6)+RANDBETWEEN(3,6))÷2)+0.3) |
|---|

**TRUNC(((RANDBETWEEN(3,6)+RANDBETWEEN(3,6))÷2)+0.3)**

Pup survival to 3-9 months normal, long right tail, mean 0.2 (0.05-0.72)

**(RANDBETWEEN(0,16)+RANDBETWEEN(0,16)+IF(RAND()<0.75,0,RANDBETWEEN(3,40)))÷100**

(RANDBETWEEN(0,16)+RANDBETWEEN(0,16)+IF(RAND()<0.75,0,RANDBETWEEN(3,40)))÷100

Number of breeding packs 74-167 uniform random

| **RANDBETWEEN(74,167)** |
|---|

| RANDBETWEEN(74,167) |
|---|

N2021 traditional =695-751 uniform

| **RANDBETWEEN(695,751)** |
|---|

| **RANDBETWEEN(695,751)** |
|---|

| RANDBETWEEN(695,751) |
|---|

| RANDBETWEEN(695,751) |
|---|

N2021new =1195 (957-1573 normal) The 218 killed in Feb 2021 wolf-hunt are deducted in subsequent steps.

| **1075+(RANDBETWEEN(−496,756)+RANDBETWEEN(−496,756))÷2** |
|---|

**1075+(RANDBETWEEN(−496,756)+RANDBETWEEN(−496,756))÷2**

**1075+(RANDBETWEEN(−496,756)+RANDBETWEEN(−496,756))÷2**

| 1075+(RANDBETWEEN(–496,756)+RANDBETWEEN(–496,756))÷2 |
|---|

Death tolls user defined

| **Any value from 0-600 could be entered here** |
|---|

| **Any value from 0-600 could be entered here** |
|---|

| Any value from 0-600 could be entered here |
|---|

| **Any value from 0-600 could be entered here** |
|---|

Death tolls mean 300 normal distribution

**(RANDBETWEEN(0,300)+RANDBETWEEN(0,300))**

**(RANDBETWEEN(0,300)+RANDBETWEEN(0,300))**

| **(RANDBETWEEN(0,300)+RANDBETWEEN(0,300))** |
|---|

| (RANDBETWEEN(0,300)+RANDBETWEEN(0,300)) |
|---|

Results (user defined death toll and paste in N2021)

| ('N2021 traditional =695-751 uniform'::A1+('Number of breeding packs 74-167 uniform random'::A1×'Packs with pups normal centered 0.72 (range 0.55-0.89)'::A1×'Litter size 4.8 (3-6) normal centered on 4.8'::A1×'Pup survival to 3-9 months normal, long right tail, mean 0.2 (0.05-0.72)'::A1)×0.5)×(1−'Deaths uniform distribution between 0.38-0.56'::A1)−Death tolls user defined::A1 |
|---|

('N2021 traditional =695-751 uniform'::A1+('Number of breeding packs 74-167 uniform random'::A1×'Packs with pups normal centered 0.72 (range 0.55-0.89)'::A1×'Litter size 4.8 (3-6) normal centered on 4.8'::A1×'Pup survival to 3-9 months normal, long right tail, mean 0.2 (0.05-0.72)'::A1)×0.5)×(1−'Deaths uniform distribution between  0.38-0.56'::A1)−Death tolls user defined::A1

| ('N2021 traditional =695-751 uniform'::A1+('Number of breeding packs 74-167 uniform random'::A1×'Packs with pups normal centered 0.72 (range 0.55-0.89)'::A1×'Litter size 4.8 (3-6) normal centered on 4.8'::A1×'Pup survival to 3-9 months normal, long right tail, mean 0.2 (0.05-0.72)'::A1)×0.5)×(1−'Deaths uniform distribution between 0.38-0.56'::A1)−Death tolls user defined::A1 |
|---|

| ('N2021 traditional =695-751 uniform'::A1+('Number of breeding packs 74-167 uniform random'::A1×'Packs with pups normal centered 0.72 (range 0.55-0.89)'::A1×'Litter size 4.8 (3-6) normal centered on 4.8'::A1×'Pup survival to 3-9 months normal, long right tail, mean 0.2 (0.05-0.72)'::A1)×0.5)×(1−'Deaths uniform distribution between 0.38-0.56'::A1)−Death tolls user defined::A1 |
|---|

Results (traditional census method)

| ('N2021 traditional =695-751 uniform'::A1+('Number of breeding packs 74-167 uniform random'::A1×'Packs with pups normal centered 0.72 (range 0.55-0.89)'::A1×'Litter size 4.8 (3-6) normal centered on 4.8'::A1×'Pup survival to 3-9 months normal, long right tail, mean 0.2 (0.05-0.72)'::A1)×0.5)×(1−'Deaths uniform distribution between 0.38-0.56'::A1)−Death tolls user defined::A1 |
|---|

**('N2021 traditional =695-751 uniform'::A1+('Number of breeding packs 74-167 uniform random'::A1×'Packs with pups normal centered 0.72 (range 0.55-0.89)'::A1×'Litter size 4.8 (3-6) normal centered on 4.8'::A1×'Pup survival to 3-9 months normal, long right tail, mean 0.2 (0.05-0.72)'::A1)×0.5)×(1−'Deaths uniform distribution between  0.38-0.56'::A1)−Death tolls user defined::A1**

**('N2021 traditional =695-751 uniform'::A1+('Number of breeding packs 74-167 uniform random'::A1×'Packs with pups normal centered 0.72 (range 0.55-0.89)'::A1×'Litter size 4.8 (3-6) normal centered on 4.8'::A1×'Pup survival to 3-9 months normal, long right tail, mean 0.2 (0.05-0.72)'::A1)×0.5)×(1−'Deaths uniform distribution between  0.38-0.56'::A1)−Death tolls user defined::A1**

**('N2021 traditional =695-751 uniform'::A1+('Number of breeding packs 74-167 uniform random'::A1×'Packs with pups normal centered 0.72 (range 0.55-0.89)'::A1×'Litter size 4.8 (3-6) normal centered on 4.8'::A1×'Pup survival to 3-9 months normal, long right tail, mean 0.2 (0.05-0.72)'::A1)×0.5)×(1−'Deaths uniform distribution between 0.38-0.56'::A1)−Death tolls user defined::A1**

Results (new census method)

**('N2021new =1195 (957-1573 normal) The 218 killed in Feb 2021 wolf-hunt are deducted in subsequent steps.'::A1−218+('Number of breeding packs 74-167 uniform random'::A1×'Packs with pups normal centered 0.72 (range 0.55-0.89)'::A1×'Litter size 4.8 (3-6) normal centered on 4.8'::A1×'Pup survival to 3-9 months normal, long right tail, mean 0.2 (0.05-0.72)'::A1)×0.5)×'Deaths uniform distribution between 0.38-0.56'::A1−Death tolls user defined::A1**

**('N2021new =1195 (957-1573 normal) The 218 killed in Feb 2021 wolf-hunt are deducted in subsequent steps.'::A1−218+('Number of breeding packs 74-167 uniform random'::A1×'Packs with pups normal centered 0.72 (range 0.55-0.89)'::A1×'Litter size 4.8 (3-6) normal centered on 4.8'::A1×'Pup survival to 3-9 months normal, long right tail, mean 0.2 (0.05-0.72)'::A1)×0.5)×'Deaths uniform distribution between 0.38-0.56'::A1−Death tolls user defined::A1**

**('N2021new =1195 (957-1573 normal) The 218 killed in Feb 2021 wolf-hunt are deducted in subsequent steps.'::A1−218+('Number of breeding packs 74-167 uniform random'::A1×'Packs with pups normal centered 0.72 (range 0.55-0.89)'::A1×'Litter size 4.8 (3-6) normal centered on 4.8'::A1×'Pup survival to 3-9 months normal, long right tail, mean 0.2 (0.05-0.72)'::A1)×0.5)×'Deaths uniform distribution between  0.38-0.56'::A1−Death tolls user defined::A1**

| ('N2021new =1195 (957-1573 normal) The 218 killed in Feb 2021 wolf-hunt are deducted in subsequent steps.'::A1−218+('Number of breeding packs 74-167 uniform random'::A1×'Packs with pups normal centered 0.72 (range 0.55-0.89)'::A1×'Litter size 4.8 (3-6) normal centered on 4.8'::A1×'Pup survival to 3-9 months normal, long right tail, mean 0.2 (0.05-0.72)'::A1)×0.5)×'Deaths uniform distribution between 0.38-0.56'::A1−Death tolls user defined::A1 |
|---|

Pairs of values N2022 v H user-defined

| ('N2021 traditional =695-751 uniform'::A1+('Number of breeding packs 74-167 uniform random'::A1×'Packs with pups normal centered 0.72 (range 0.55-0.89)'::A1×'Litter size 4.8 (3-6) normal centered on 4.8'::A1×'Pup survival to 3-9 months normal, long right tail, mean 0.2 (0.05-0.72)'::A1)×0.5)×(1−'Deaths uniform distribution between 0.38-0.56'::A1)−Death tolls user defined::A1 | Any value from 0-600 could be entered here |
|---:|---:|
| **0** | 0 |
| **0** | 0 |
| **0** | 0 |
| **0** | 0 |
| **0** | 0 |
| **0** | 0 |
| **0** | 0 |
| **0** | 0 |
| **0** | 0 |
| **0** | 0 |
| **0** | 0 |
| **0** | 0 |
| **0** | 0 |
| **0** | 0 |
| **0** | 0 |
| **0** | 0 |
| **0** | 0 |
| **0** | 0 |
| **0** | 0 |
| **0** | 0 |
| **0** | 0 |
| **0** | 0 |
| **0** | 0 |
| **0** | 0 |
| **0** | 0 |
| **0** | 0 |

| ('N2021 traditional =695-751 uniform'::A1+('Number of breeding packs 74-167 uniform random'::A1×'Packs with pups normal centered 0.72 (range 0.55-0.89)'::A1×'Litter size 4.8 (3-6) normal centered on 4.8'::A1×'Pup survival to 3-9 months normal, long right tail, mean 0.2 (0.05-0.72)'::A1)×0.5)×(1−'Deaths uniform distribution between 0.38-0.56'::A1)−Death tolls user defined::A1 | Any value from 0-600 could be entered here |
|---|---|
| **0** | 0 |
| **0** | 0 |
| **0** | 0 |
| **0** | 0 |
| **0** | 0 |
| **0** | 0 |
| **0** | 0 |
| **0** | 0 |
| **0** | 0 |
| **0** | 0 |
| **0** | 0 |
| **0** | 0 |
| **0** | 0 |
| **0** | 0 |
| **0** | 0 |
| **0** | 0 |
| **0** | 0 |
| **0** | 0 |
| **0** | 0 |
| **0** | 0 |
| **0** | 0 |
| **0** | 0 |
| **0** | 0 |
| **0** | 0 |
| **0** | 0 |
| **0** | 0 |
| **0** | 0 |

| ('N2021 traditional =695-751 uniform'::A1+('Number of breeding packs 74-167 uniform random'::A1×'Packs with pups normal centered 0.72 (range 0.55-0.89)'::A1×'Litter size 4.8 (3-6) normal centered on 4.8'::A1×'Pup survival to 3-9 months normal, long right tail, mean 0.2 (0.05-0.72)'::A1)×0.5)×(1−'Deaths uniform distribution between 0.38-0.56'::A1)−Death tolls user defined::A1 | Any value from 0-600 could be entered here |
|---:|---:|
| **0** | 0 |
| **0** | 0 |
| **0** | 0 |
| **0** | 0 |
| **0** | 0 |
| **0** | 0 |
| **0** | 0 |
| **0** | 0 |
| **0** | 0 |
| **0** | 0 |
| **0** | 0 |
| **0** | 0 |
| **0** | 0 |
| **0** | 0 |
| **0** | 0 |
| **0** | 0 |
| **0** | 0 |
| **0** | 0 |
| **0** | 0 |
| **0** | 0 |
| **0** | 0 |
| **0** | 0 |
| **0** | 0 |
| **0** | 0 |
| **0** | 0 |
| **0** | 0 |
| **0** | 0 |

| ('N2021 traditional =695-751 uniform'::A1+('Number of breeding packs 74-167 uniform random'::A1×'Packs with pups normal centered 0.72 (range 0.55-0.89)'::A1×'Litter size 4.8 (3-6) normal centered on 4.8'::A1×'Pup survival to 3-9 months normal, long right tail, mean 0.2 (0.05-0.72)'::A1)×0.5)×(1−'Deaths uniform distribution between 0.38-0.56'::A1)−Death tolls user defined::A1 | Any value from 0-600 could be entered here |
|---|---|
| **0** | 0 |
| **0** | 0 |
| **0** | 0 |
| **0** | 0 |
| **0** | 0 |
| **0** | 0 |
| **0** | 0 |
| **0** | 0 |
| **0** | 0 |
| **0** | 0 |
| **0** | 0 |
| **0** | 0 |
| **0** | 0 |
| **0** | 0 |
| **0** | 0 |
| **0** | 0 |
| **0** | 0 |
| **0** | 0 |
| **0** | 0 |
| **0** | 0 |
| **0** | 0 |
| **0** | 0 |
| **0** | 0 |
| **0** | 0 |

| ('N2021 traditional =695-751 uniform'::A1+('Number of breeding packs 74-167 uniform random'::A1×'Packs with pups normal centered 0.72 (range 0.55-0.89)'::A1×'Litter size 4.8 (3-6) normal centered on 4.8'::A1×'Pup survival to 3-9 months normal, long right tail, mean 0.2 (0.05-0.72)'::A1)×0.5)×(1−'Deaths uniform distribution between 0.38-0.56'::A1)−Death tolls user defined::A1 | Any value from 0-600 could be entered here |
|---|---|

| ('N2021 traditional =695-751 uniform'::A1+('Number of breeding packs 74-167 uniform random'::A1×'Packs with pups normal centered 0.72 (range 0.55-0.89)'::A1×'Litter size 4.8 (3-6) normal centered on 4.8'::A1×'Pup survival to 3-9 months normal, long right tail, mean 0.2 (0.05-0.72)'::A1)×0.5)×(1−'Deaths uniform distribution between 0.38-0.56'::A1)−Death tolls user defined::A1 | Any value from 0-600 could be entered here |
|---|---|

| ('N2021 traditional =695-751 uniform'::A1+('Number of breeding packs 74-167 uniform random'::A1×'Packs with pups normal centered 0.72 (range 0.55-0.89)'::A1×'Litter size 4.8 (3-6) normal centered on 4.8'::A1×'Pup survival to 3-9 months normal, long right tail, mean 0.2 (0.05-0.72)'::A1)×0.5)×(1−'Deaths uniform distribution between 0.38-0.56'::A1)−Death tolls user defined::A1 | Any value from 0-600 could be entered here |
|---|---|

| ('N2021 traditional =695-751 uniform'::A1+('Number of breeding packs 74-167 uniform random'::A1×'Packs with pups normal centered 0.72 (range 0.55-0.89)'::A1×'Litter size 4.8 (3-6) normal centered on 4.8'::A1×'Pup survival to 3-9 months normal, long right tail, mean 0.2 (0.05-0.72)'::A1)×0.5)×(1−'Deaths uniform distribution between 0.38-0.56'::A1)−Death tolls user defined::A1 | Any value from 0-600 could be entered here |
|---|---|

| ('N2021 traditional =695-751 uniform'::A1+('Number of breeding packs 74-167 uniform random'::A1×'Packs with pups normal centered 0.72 (range 0.55-0.89)'::A1×'Litter size 4.8 (3-6) normal centered on 4.8'::A1×'Pup survival to 3-9 months normal, long right tail, mean 0.2 (0.05-0.72)'::A1)×0.5)×(1−'Deaths uniform distribution between 0.38-0.56'::A1)−Death tolls user defined::A1 | Any value from 0-600 could be entered here |
|---|---|

| ('N2021 traditional =695-751 uniform'::A1+('Number of breeding packs 74-167 uniform random'::A1×'Packs with pups normal centered 0.72 (range 0.55-0.89)'::A1×'Litter size 4.8 (3-6) normal centered on 4.8'::A1×'Pup survival to 3-9 months normal, long right tail, mean 0.2 (0.05-0.72)'::A1)×0.5)×(1−'Deaths uniform distribution between 0.38-0.56'::A1)−Death tolls user defined::A1 | Any value from 0-600 could be entered here |
|---|---|

| ('N2021 traditional =695-751 uniform'::A1+('Number of breeding packs 74-167 uniform random'::A1×'Packs with pups normal centered 0.72 (range 0.55-0.89)'::A1×'Litter size 4.8 (3-6) normal centered on 4.8'::A1×'Pup survival to 3-9 months normal, long right tail, mean 0.2 (0.05-0.72)'::A1)×0.5)×(1−'Deaths uniform distribution between 0.38-0.56'::A1)−Death tolls user defined::A1 | Any value from 0-600 could be entered here |
|---|---|

| ('N2021 traditional =695-751 uniform'::A1+('Number of breeding packs 74-167 uniform random'::A1×'Packs with pups normal centered 0.72 (range 0.55-0.89)'::A1×'Litter size 4.8 (3-6) normal centered on 4.8'::A1×'Pup survival to 3-9 months normal, long right tail, mean 0.2 (0.05-0.72)'::A1)×0.5)×(1−'Deaths uniform distribution between 0.38-0.56'::A1)−Death tolls user defined::A1 | Any value from 0-600 could be entered here |
|---|---|

| ('N2021 traditional =695-751 uniform'::A1+('Number of breeding packs 74-167 uniform random'::A1×'Packs with pups normal centered 0.72 (range 0.55-0.89)'::A1×'Litter size 4.8 (3-6) normal centered on 4.8'::A1×'Pup survival to 3-9 months normal, long right tail, mean 0.2 (0.05-0.72)'::A1)×0.5)×(1−'Deaths uniform distribution between 0.38-0.56'::A1)−Death tolls user defined::A1 | Any value from 0-600 could be entered here |
|---|---|

| ('N2021 traditional =695-751 uniform'::A1+('Number of breeding packs 74-167 uniform random'::A1×'Packs with pups normal centered 0.72 (range 0.55-0.89)'::A1×'Litter size 4.8 (3-6) normal centered on 4.8'::A1×'Pup survival to 3-9 months normal, long right tail, mean 0.2 (0.05-0.72)'::A1)×0.5)×(1−'Deaths uniform distribution between 0.38-0.56'::A1)−Death tolls user defined::A1 | Any value from 0-600 could be entered here |
|---|---|

| ('N2021 traditional =695-751 uniform'::A1+('Number of breeding packs 74-167 uniform random'::A1×'Packs with pups normal centered 0.72 (range 0.55-0.89)'::A1×'Litter size 4.8 (3-6) normal centered on 4.8'::A1×'Pup survival to 3-9 months normal, long right tail, mean 0.2 (0.05-0.72)'::A1)×0.5)×(1−'Deaths uniform distribution between 0.38-0.56'::A1)−Death tolls user defined::A1 | Any value from 0-600 could be entered here |
|---|---|

| ('N2021 traditional =695-751 uniform'::A1+('Number of breeding packs 74-167 uniform random'::A1×'Packs with pups normal centered 0.72 (range 0.55-0.89)'::A1×'Litter size 4.8 (3-6) normal centered on 4.8'::A1×'Pup survival to 3-9 months normal, long right tail, mean 0.2 (0.05-0.72)'::A1)×0.5)×(1−'Deaths uniform distribution between 0.38-0.56'::A1)−Death tolls user defined::A1 | Any value from 0-600 could be entered here |
|---|---|

| ('N2021 traditional =695-751 uniform'::A1+('Number of breeding packs 74-167 uniform random'::A1×'Packs with pups normal centered 0.72 (range 0.55-0.89)'::A1×'Litter size 4.8 (3-6) normal centered on 4.8'::A1×'Pup survival to 3-9 months normal, long right tail, mean 0.2 (0.05-0.72)'::A1)×0.5)×(1−'Deaths uniform distribution between 0.38-0.56'::A1)−Death tolls user defined::A1 | Any value from 0-600 could be entered here |
|---|---|

| ('N2021 traditional =695-751 uniform'::A1+('Number of breeding packs 74-167 uniform random'::A1×'Packs with pups normal centered 0.72 (range 0.55-0.89)'::A1×'Litter size 4.8 (3-6) normal centered on 4.8'::A1×'Pup survival to 3-9 months normal, long right tail, mean 0.2 (0.05-0.72)'::A1)×0.5)×(1−'Deaths uniform distribution between 0.38-0.56'::A1)−Death tolls user defined::A1 | Any value from 0-600 could be entered here |
|---|---|

| ('N2021 traditional =695-751 uniform'::A1+('Number of breeding packs 74-167 uniform random'::A1×'Packs with pups normal centered 0.72 (range 0.55-0.89)'::A1×'Litter size 4.8 (3-6) normal centered on 4.8'::A1×'Pup survival to 3-9 months normal, long right tail, mean 0.2 (0.05-0.72)'::A1)×0.5)×(1−'Deaths uniform distribution between 0.38-0.56'::A1)−Death tolls user defined::A1 | Any value from 0-600 could be entered here |
|---|---|

| **('N2021 traditional =695-751 uniform'::A1+('Number of breeding packs 74-167 uniform random'::A1×'Packs with pups normal centered 0.72 (range 0.55-0.89)'::A1×'Litter size 4.8 (3-6) normal centered on 4.8'::A1×'Pup survival to 3-9 months normal, long right tail, mean 0.2 (0.05-0.72)'::A1)×0.5)×(1−'Deaths uniform distribution between 0.38-0.56'::A1)−Death tolls user defined::A1** | **Any value from 0-600 could be entered here** |
| --- | --- |

| ('N2021 traditional =695-751 uniform'::A1+('Number of breeding packs 74-167 uniform random'::A1×'Packs with pups normal centered 0.72 (range 0.55-0.89)'::A1×'Litter size 4.8 (3-6) normal centered on 4.8'::A1×'Pup survival to 3-9 months normal, long right tail, mean 0.2 (0.05-0.72)'::A1)×0.5)×(1−'Deaths uniform distribution between 0.38-0.56'::A1)−Death tolls user defined::A1 | Any value from 0-600 could be entered here |
|---|---|

| ('N2021 traditional =695-751 uniform'::A1+('Number of breeding packs 74-167 uniform random'::A1×'Packs with pups normal centered 0.72 (range 0.55-0.89)'::A1×'Litter size 4.8 (3-6) normal centered on 4.8'::A1×'Pup survival to 3-9 months normal, long right tail, mean 0.2 (0.05-0.72)'::A1)×0.5)×(1−'Deaths uniform distribution between 0.38-0.56'::A1)−Death tolls user defined::A1 | Any value from 0-600 could be entered here |
|---|---|

| ('N2021 traditional =695-751 uniform'::A1+('Number of breeding packs 74-167 uniform random'::A1×'Packs with pups normal centered 0.72 (range 0.55-0.89)'::A1×'Litter size 4.8 (3-6) normal centered on 4.8'::A1×'Pup survival to 3-9 months normal, long right tail, mean 0.2 (0.05-0.72)'::A1)×0.5)×(1−'Deaths uniform distribution between 0.38-0.56'::A1)−Death tolls user defined::A1 | Any value from 0-600 could be entered here |
|---|---|

| ('N2021 traditional =695-751 uniform'::A1+('Number of breeding packs 74-167 uniform random'::A1×'Packs with pups normal centered 0.72 (range 0.55-0.89)'::A1×'Litter size 4.8 (3-6) normal centered on 4.8'::A1×'Pup survival to 3-9 months normal, long right tail, mean 0.2 (0.05-0.72)'::A1)×0.5)×(1−'Deaths uniform distribution between 0.38-0.56'::A1)−Death tolls user defined::A1 | Any value from 0-600 could be entered here |
|---|---|

| ('N2021 traditional =695-751 uniform'::A1+('Number of breeding packs 74-167 uniform random'::A1×'Packs with pups normal centered 0.72 (range 0.55-0.89)'::A1×'Litter size 4.8 (3-6) normal centered on 4.8'::A1×'Pup survival to 3-9 months normal, long right tail, mean 0.2 (0.05-0.72)'::A1)×0.5)×(1−'Deaths uniform distribution between 0.38-0.56'::A1)−Death tolls user defined::A1 | Any value from 0-600 could be entered here |
|---|---|

| ('N2021 traditional =695-751 uniform'::A1+('Number of breeding packs 74-167 uniform random'::A1×'Packs with pups normal centered 0.72 (range 0.55-0.89)'::A1×'Litter size 4.8 (3-6) normal centered on 4.8'::A1×'Pup survival to 3-9 months normal, long right tail, mean 0.2 (0.05-0.72)'::A1)×0.5)×(1−'Deaths uniform distribution between 0.38-0.56'::A1)−Death tolls user defined::A1 | Any value from 0-600 could be entered here |
|---|---|

| ('N2021 traditional =695-751 uniform'::A1+('Number of breeding packs 74-167 uniform random'::A1×'Packs with pups normal centered 0.72 (range 0.55-0.89)'::A1×'Litter size 4.8 (3-6) normal centered on 4.8'::A1×'Pup survival to 3-9 months normal, long right tail, mean 0.2 (0.05-0.72)'::A1)×0.5)×(1−'Deaths uniform distribution between 0.38-0.56'::A1)−Death tolls user defined::A1 | Any value from 0-600 could be entered here |
|---|---|

| ('N2021 traditional =695-751 uniform'::A1+('Number of breeding packs 74-167 uniform random'::A1×'Packs with pups normal centered 0.72 (range 0.55-0.89)'::A1×'Litter size 4.8 (3-6) normal centered on 4.8'::A1×'Pup survival to 3-9 months normal, long right tail, mean 0.2 (0.05-0.72)'::A1)×0.5)×(1−'Deaths uniform distribution between 0.38-0.56'::A1)−Death tolls user defined::A1 | Any value from 0-600 could be entered here |
|---|---|

| ('N2021 traditional =695-751 uniform'::A1+('Number of breeding packs 74-167 uniform random'::A1×'Packs with pups normal centered 0.72 (range 0.55-0.89)'::A1×'Litter size 4.8 (3-6) normal centered on 4.8'::A1×'Pup survival to 3-9 months normal, long right tail, mean 0.2 (0.05-0.72)'::A1)×0.5)×(1−'Deaths uniform distribution between 0.38-0.56'::A1)−Death tolls user defined::A1 | Any value from 0-600 could be entered here |
|---|---|

| ('N2021 traditional =695-751 uniform'::A1+('Number of breeding packs 74-167 uniform random'::A1×'Packs with pups normal centered 0.72 (range 0.55-0.89)'::A1×'Litter size 4.8 (3-6) normal centered on 4.8'::A1×'Pup survival to 3-9 months normal, long right tail, mean 0.2 (0.05-0.72)'::A1)×0.5)×(1−'Deaths uniform distribution between 0.38-0.56'::A1)−Death tolls user defined::A1 | Any value from 0-600 could be entered here |
|---|---|

| ('N2021 traditional =695-751 uniform'::A1+('Number of breeding packs 74-167 uniform random'::A1×'Packs with pups normal centered 0.72 (range 0.55-0.89)'::A1×'Litter size 4.8 (3-6) normal centered on 4.8'::A1×'Pup survival to 3-9 months normal, long right tail, mean 0.2 (0.05-0.72)'::A1)×0.5)×(1−'Deaths uniform distribution between 0.38-0.56'::A1)−Death tolls user defined::A1 | Any value from 0-600 could be entered here |
|---|---|

| ('N2021 traditional =695-751 uniform'::A1+('Number of breeding packs 74-167 uniform random'::A1×'Packs with pups normal centered 0.72 (range 0.55-0.89)'::A1×'Litter size 4.8 (3-6) normal centered on 4.8'::A1×'Pup survival to 3-9 months normal, long right tail, mean 0.2 (0.05-0.72)'::A1)×0.5)×(1−'Deaths uniform distribution between 0.38-0.56'::A1)−Death tolls user defined::A1 | Any value from 0-600 could be entered here |
|---|---|

| ('N2021 traditional =695-751 uniform'::A1+('Number of breeding packs 74-167 uniform random'::A1×'Packs with pups normal centered 0.72 (range 0.55-0.89)'::A1×'Litter size 4.8 (3-6) normal centered on 4.8'::A1×'Pup survival to 3-9 months normal, long right tail, mean 0.2 (0.05-0.72)'::A1)×0.5)×(1−'Deaths uniform distribution between 0.38-0.56'::A1)−Death tolls user defined::A1 | Any value from 0-600 could be entered here |
|---|---|

| ('N2021 traditional =695-751 uniform'::A1+('Number of breeding packs 74-167 uniform random'::A1×'Packs with pups normal centered 0.72 (range 0.55-0.89)'::A1×'Litter size 4.8 (3-6) normal centered on 4.8'::A1×'Pup survival to 3-9 months normal, long right tail, mean 0.2 (0.05-0.72)'::A1)×0.5)×(1−'Deaths uniform distribution between 0.38-0.56'::A1)−Death tolls user defined::A1 | Any value from 0-600 could be entered here |
|---|---|

| ('N2021 traditional =695-751 uniform'::A1+('Number of breeding packs 74-167 uniform random'::A1×'Packs with pups normal centered 0.72 (range 0.55-0.89)'::A1×'Litter size 4.8 (3-6) normal centered on 4.8'::A1×'Pup survival to 3-9 months normal, long right tail, mean 0.2 (0.05-0.72)'::A1)×0.5)×(1−'Deaths uniform distribution between 0.38-0.56'::A1)−Death tolls user defined::A1 | Any value from 0-600 could be entered here |
|---|---|

| ('N2021 traditional =695-751 uniform'::A1+('Number of breeding packs 74-167 uniform random'::A1×'Packs with pups normal centered 0.72 (range 0.55-0.89)'::A1×'Litter size 4.8 (3-6) normal centered on 4.8'::A1×'Pup survival to 3-9 months normal, long right tail, mean 0.2 (0.05-0.72)'::A1)×0.5)×(1−'Deaths uniform distribution between 0.38-0.56'::A1)−Death tolls user defined::A1 | Any value from 0-600 could be entered here |
|---|---|

| ('N2021 traditional =695-751 uniform'::A1+('Number of breeding packs 74-167 uniform random'::A1×'Packs with pups normal centered 0.72 (range 0.55-0.89)'::A1×'Litter size 4.8 (3-6) normal centered on 4.8'::A1×'Pup survival to 3-9 months normal, long right tail, mean 0.2 (0.05-0.72)'::A1)×0.5)×(1−'Deaths uniform distribution between 0.38-0.56'::A1)−Death tolls user defined::A1 | Any value from 0-600 could be entered here |
|---|---|

| ('N2021 traditional =695-751 uniform'::A1+('Number of breeding packs 74-167 uniform random'::A1×'Packs with pups normal centered 0.72 (range 0.55-0.89)'::A1×'Litter size 4.8 (3-6) normal centered on 4.8'::A1×'Pup survival to 3-9 months normal, long right tail, mean 0.2 (0.05-0.72)'::A1)×0.5)×(1−'Deaths uniform distribution between 0.38-0.56'::A1)−Death tolls user defined::A1 | Any value from 0-600 could be entered here |
|---|---|

| ('N2021 traditional =695-751 uniform'::A1+('Number of breeding packs 74-167 uniform random'::A1×'Packs with pups normal centered 0.72 (range 0.55-0.89)'::A1×'Litter size 4.8 (3-6) normal centered on 4.8'::A1×'Pup survival to 3-9 months normal, long right tail, mean 0.2 (0.05-0.72)'::A1)×0.5)×(1−'Deaths uniform distribution between 0.38-0.56'::A1)−Death tolls user defined::A1 | Any value from 0-600 could be entered here |
|---|---|

| ('N2021 traditional =695-751 uniform'::A1+('Number of breeding packs 74-167 uniform random'::A1×'Packs with pups normal centered 0.72 (range 0.55-0.89)'::A1×'Litter size 4.8 (3-6) normal centered on 4.8'::A1×'Pup survival to 3-9 months normal, long right tail, mean 0.2 (0.05-0.72)'::A1)×0.5)×(1−'Deaths uniform distribution between 0.38-0.56'::A1)−Death tolls user defined::A1 | Any value from 0-600 could be entered here |
|---|---|

| ('N2021 traditional =695-751 uniform'::A1+('Number of breeding packs 74-167 uniform random'::A1×'Packs with pups normal centered 0.72 (range 0.55-0.89)'::A1×'Litter size 4.8 (3-6) normal centered on 4.8'::A1×'Pup survival to 3-9 months normal, long right tail, mean 0.2 (0.05-0.72)'::A1)×0.5)×(1−'Deaths uniform distribution between 0.38-0.56'::A1)−Death tolls user defined::A1 | Any value from 0-600 could be entered here |
|---|---|

| ('N2021 traditional =695-751 uniform'::A1+('Number of breeding packs 74-167 uniform random'::A1×'Packs with pups normal centered 0.72 (range 0.55-0.89)'::A1×'Litter size 4.8 (3-6) normal centered on 4.8'::A1×'Pup survival to 3-9 months normal, long right tail, mean 0.2 (0.05-0.72)'::A1)×0.5)×(1−'Deaths uniform distribution between 0.38-0.56'::A1)−Death tolls user defined::A1 | Any value from 0-600 could be entered here |
| --- | --- |

| ('N2021 traditional =695-751 uniform'::A1+('Number of breeding packs 74-167 uniform random'::A1×'Packs with pups normal centered 0.72 (range 0.55-0.89)'::A1×'Litter size 4.8 (3-6) normal centered on 4.8'::A1×'Pup survival to 3-9 months normal, long right tail, mean 0.2 (0.05-0.72)'::A1)×0.5)×(1−'Deaths uniform distribution between 0.38-0.56'::A1)−Death tolls user defined::A1 | Any value from 0-600 could be entered here |
|---|---|

| ('N2021 traditional =695-751 uniform'::A1+('Number of breeding packs 74-167 uniform random'::A1×'Packs with pups normal centered 0.72 (range 0.55-0.89)'::A1×'Litter size 4.8 (3-6) normal centered on 4.8'::A1×'Pup survival to 3-9 months normal, long right tail, mean 0.2 (0.05-0.72)'::A1)×0.5)×(1−'Deaths uniform distribution between 0.38-0.56'::A1)−Death tolls user defined::A1 | Any value from 0-600 could be entered here |
|---|---|

| ('N2021 traditional =695-751 uniform'::A1+('Number of breeding packs 74-167 uniform random'::A1×'Packs with pups normal centered 0.72 (range 0.55-0.89)'::A1×'Litter size 4.8 (3-6) normal centered on 4.8'::A1×'Pup survival to 3-9 months normal, long right tail, mean 0.2 (0.05-0.72)'::A1)×0.5)×(1−'Deaths uniform distribution between 0.38-0.56'::A1)−Death tolls user defined::A1 | Any value from 0-600 could be entered here |
|---|---|
